# Hill-type models of skeletal muscle and neuromuscular actuators: a systematic review

**DOI:** 10.1101/2022.10.14.512218

**Authors:** Arnault H. Caillet, Andrew T.M. Phillips, Christopher Carty, Dario Farina, Luca Modenese

**Affiliations:** Department of Bioengineering, Imperial College London, SW7 2AZ, UK; Department of Civil and Environmental Engineering, Imperial College London, SW7 2AZ, UK; Griffith Centre of Biomedical and Rehabilitation Engineering (GCORE), Griffith University, Australia; School of Medicine and Dentistry, Griffith University, Australia; Department of Orthopaedics, Children’s Health Queensland Hospital and Health Service, Brisbane, Australia; Graduate School of Biomedical Engineering, University of New South Wales, Sydney, Australia

## Abstract

Backed by a century of research and development, Hill-type models of skeletal muscle, often including a muscle-tendon complex and neuromechanical interface, are widely used for countless applications. Lacking recent comprehensive reviews, the field of Hill-type modelling is, however, dense and hard-to-explore, with detrimental consequences on innovation. Here we present the first systematic review of Hill-type muscle modelling. It aims to clarify the literature by detailing its contents and critically discussing the state-of-the-art by identifying the latest advances, current gaps, and potential future directions in Hill-type modelling. For this purpose, fifty-seven criteria-abiding Hill-type models were assessed according to a completeness evaluation, which identified the modelled muscle properties, and a modelling evaluation, which considered the level of validation and reusability of the models, as well as their modelling strategy and calibration. It is concluded that most models (1) do not significantly advance beyond historical gold standards, (2) neglect the importance of parameter identification, (3) lack robust validation, and (4) are not reusable in other studies. Besides providing a convenient tool supported by extensive supplementary material for navigating the literature, the results of this review highlight the need for global recommendations in Hill-type modelling to optimize inter-study consistency, knowledge transfer, and model reusability.

## I. INTRODUCTION

Computational models of skeletal muscle mimic the dynamics of muscle contraction by predicting the time profile of relevant muscle-related quantities that are challenging or impossible to obtain in vivo [1]-[4], including muscle force, muscle-tendon interaction and fascicle length and velocity. Among these models, Hill-type models are globally more popular than other available solutions, such as analytical descriptions of the muscle twitch [5]-[7], Huxley-type cross-bridge models [8]-[13], continuum models [14]-[17], and other comprehensive models [18], for several reasons. With sarcomere-to-muscle multiscale simplifications and phenomenological modelling approaches, Hill-type models rely on fewer parameters and are computationally cheaper and conceptually simpler [13], [19]-[21] than Huxley-type or continuum models, while they are complex enough to describe the chain of neuromechanical events responsible for muscle force generation, which makes them more accurate, flexible and amenable to further refinements [20] than analytical models. Hill-type models [22], [23] build upon a century of advancing research, a historical excursus of which is provided in Supplementary Material 1 (SM1). The predictive accuracy of these models has been extensively validated for various modelling variants, species, subjects, muscles and tasks. Moreover, Hill-type models were gradually advanced to include a representation of the coupling between neural commands and force generation, in this review called “neuromechanical interface”, effectively becoming neuromuscular actuators. Previous neuromuscular models, for example, could predict muscle forces from experimental neural input such as bipolar [24]-[27] and high-density [28], [29] electromyograms (bEMG and HD-EMG) or Functional Electrical Stimulation (FES) [30]. Also, Hill-type models, for which some open-source implementations are available [29], [31]-[36], are embedded in user-friendly biomechanical simulation platforms, including OpenSim [37], [38], AnyBody [39], MyoSuite [40] and SCONE [41]. For these reasons, Hill-type models are more accessible in experimental data-driven musculoskeletal (MSK) investigations than other solutions. Hill-type models are therefore used in a wide range of applications, either (1) to investigate the muscle-tendon interplay [23], [42], muscle energetics [43]-[45], contractile mechanics [46], [47] and neuromechanics [48], and age-related and neuromuscular impairment-related effects [49]-[52]; (2) to explore motor control [40], [53]-[55], muscle reflex [56]-[58], and the response of the human body to perturbations [59]-[61]; (3) in MSK studies to assess the muscles’ interactions with surrounding structures, such as the muscles’ relative force contribution to joint torques, motion and contact forces [51], [62]-[65], the indeterminacy of the load sharing system [66], [67], or muscle-induced stress development in skeletal bodies [68]-[71]; (4) in clinical applications to predict the outcome of surgical procedures [72]-[74] or optimizing rehabilitation [75]-[78], or for the design and control of prosthetics [79]-[83]; or (5) in sports-related applications, such as for optimizing performance [84]-[87] or predicting the effects of muscle fatigue on performance [88]-[94].

Hill-type models are therefore flexible and can fulfil different objectives, with a suitable compromise between model complexity and accuracy. To choose the optimal Hill-type variant, one must have a clear understanding of the contents of the literature in Hill-type modelling. Yet, the field of Hill-type modelling is currently extremely dense, with several thousand models proposed in the past 50 years. The field is moreover hard-to-explore since the last available extensive narrative reviews are more than 20 years old, while the more recent studies that investigate the existing modelling approaches and assess the gaps in the literature are not exhaustive, as they focus on specific aspects of the field of muscle modelling [20], [23], [95]-[100]. This situation drastically limits knowledge transfer and makes it difficult and time-consuming for researchers to identify the ‘right’ Hill-type model in the literature. This situation also slows down the pace of models’ improvement, despite the important remaining limitations discussed in the following, as it is difficult to build upon an ill-defined state-of-the-art. This situation also promotes detrimental tendencies, such as ill-referencing, inter-study inconsistency in terminologies, definitions and notations, and overlooking most recent contributions. As advised in the 90s [98], contemporary literature and systematic reviews of the literature must be available to organize and clarify the contents and state-of-the-art of the field. This was recently done, for example, in skeletal muscle continuum modelling [101], multiscale neuromuscular modelling [16], advanced muscle modelling in MSK simulations [102], muscle modelling with motor unit resolution [103], cervical spine MSK modelling [104], myoelectric control of exoskeletons [105], and lower limb force modelling during gait [106].

In this respect, this study provides the first systematic review of the field of Hill-type skeletal muscle and neuromuscular modelling. Its contents and state-of-the-art are assessed with two scoring schemes, a completeness assessment, and a modelling evaluation, applied to a set of 57 eligible studies, that were extracted from a systematic search for published Hill-type models. The eligible studies were chosen to be representative of all the modelling approaches taken to describe 23 neuromuscular and musculotendon properties that underly the muscle-force generation process. The completeness assessment identifies which muscle properties are described in an eligible model, while the modelling assessment evaluates the level of validation and reusability of the model and attention given to its modelling strategy and calibration. The results of the scoring schemes are used to discuss the current trends and gold standards in Hill-type modelling, challenge them, investigate the state-of-the art, identify the current gaps and lines of improvement, and open the discussion on the necessity of global recommendations to promote model reusability and knowledge transfer against the current scarcity of open-source models and lack of consensus on good practices. This study is supported by extensive supplementary material and can be used as a convenient time-saving tool to navigate through the literature, since any published model can be retrieved by combining the modelling approaches proposed in the 57 eligible studies, and to develop novel Hill-type models answering current limitations.

## II. METHODS

This systematic review was performed following the PRISMA guidelines [107] (see the commented checklist in SM2) by applying two scoring schemes - a completeness assessment and a modelling evaluation – to criteria-abiding Hill-type modelling studies extracted from an online systematic search (Table 1).

**Table 1:** Definition of the four keyterm concepts C1-C4 for the systematic search (see SM3 for details). Definition of the eligibility criteria EC1-EC6.

|  |  |
| --- | --- |
| C1 | The Hill-type musculotendon model |
| C2 | The muscle, the tendon |
| C3 | The model, the actuator, the simulation |
| C4 | The smooth muscle, the cardiac muscles |
| EC1 | The studies are Journal articles in English |
| EC2 | Only skeletal muscles are considered |
| EC3 | The muscle model is of the Hill-type |
| EC4 | An explicit description of the properties of at least a 'baseline' Hill-type model is provided (Fig. 1) |
| EC5 | The dynamics of the model rely on length- and force-dependent state variables |
| EC6 | The model is 'innovative' |

### A. Terminology, definitions, and acronyms

The definition of a Hill-type model has been continuously refined and extended since Hill’s preliminary Nobel prize-winning works [22], [108], [109], as detailed in the historical excursus in SM1. A Hill-type model refers today, consistently with the terminology proposed in milestone reviews in the field [23], [95], [98], to any rheological arrangement of an active force generator (FG), which builds the active muscle force according to a contraction dynamics, and in-series and/or in-parallel passive elements, which passively transmit force through the stretch-resisting dynamics of their constitutive material. Examples are displayed in Fig. 1. These generic FG and passive elements can appropriately describe the process of force generation at multiple scales in sarcomeres, fibres or entire muscles with appropriate ‘multiscale’ modelling assumptions and simplifications, that are usually present in Hill-type actuators and discussed in the following. In the literature, the behaviour of these active and passive rheological elements is governed by mathematical expressions that describe up to 23 phenomenological active or passive neuromuscular properties, which are displayed along with their acronyms in Table 2 and Fig. 2.

**Table 2:**
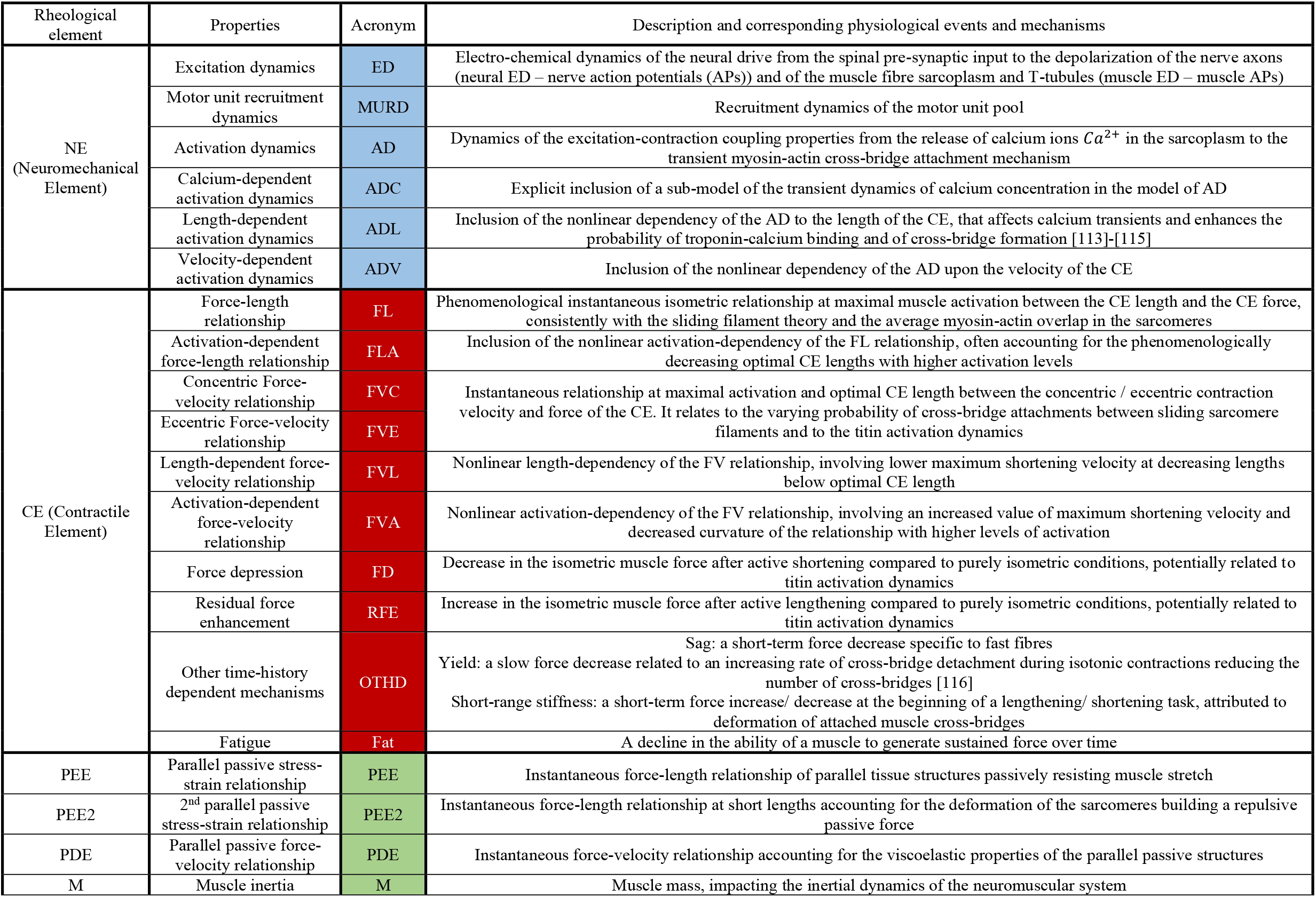
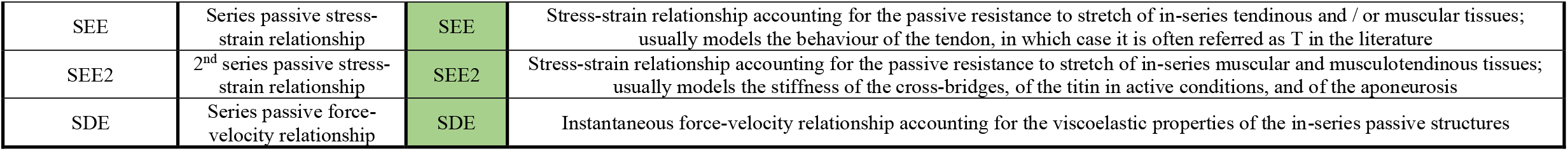
Hill-type models are described with 9 possible rheological elements and 23 possible phenomenological properties. This table provides for these elements and properties brief descriptions and the acronyms that are used in this review. ADL, ADV, FLA, FVL and FVA refer to a nonlinear dependency of the model activation dynamics (AD) and Force-Length (FL) and Force-Velocity (FV) relationships to the model active (A), length (L) and velocity (V) states.

**Fig. 1.**
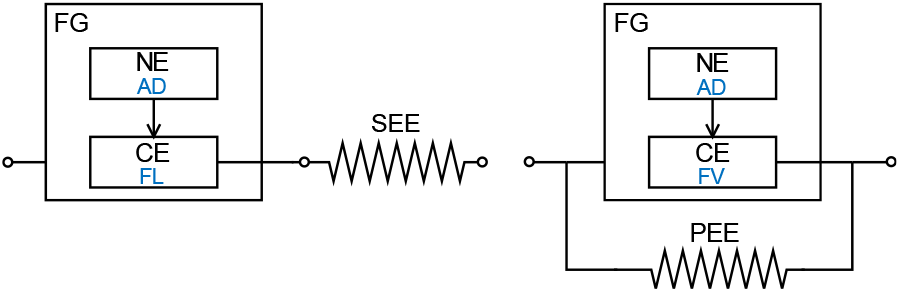
Example of two ‘baseline’ Hill-type models. They include one passive element (Left: SEE and Right: PEE), and an FG that involves a NE, described here by one AD property, and a CE, described here by one FL (Left) or FV (Right) relationship. The acronyms are defined in Table 2.

**Fig. 2.**
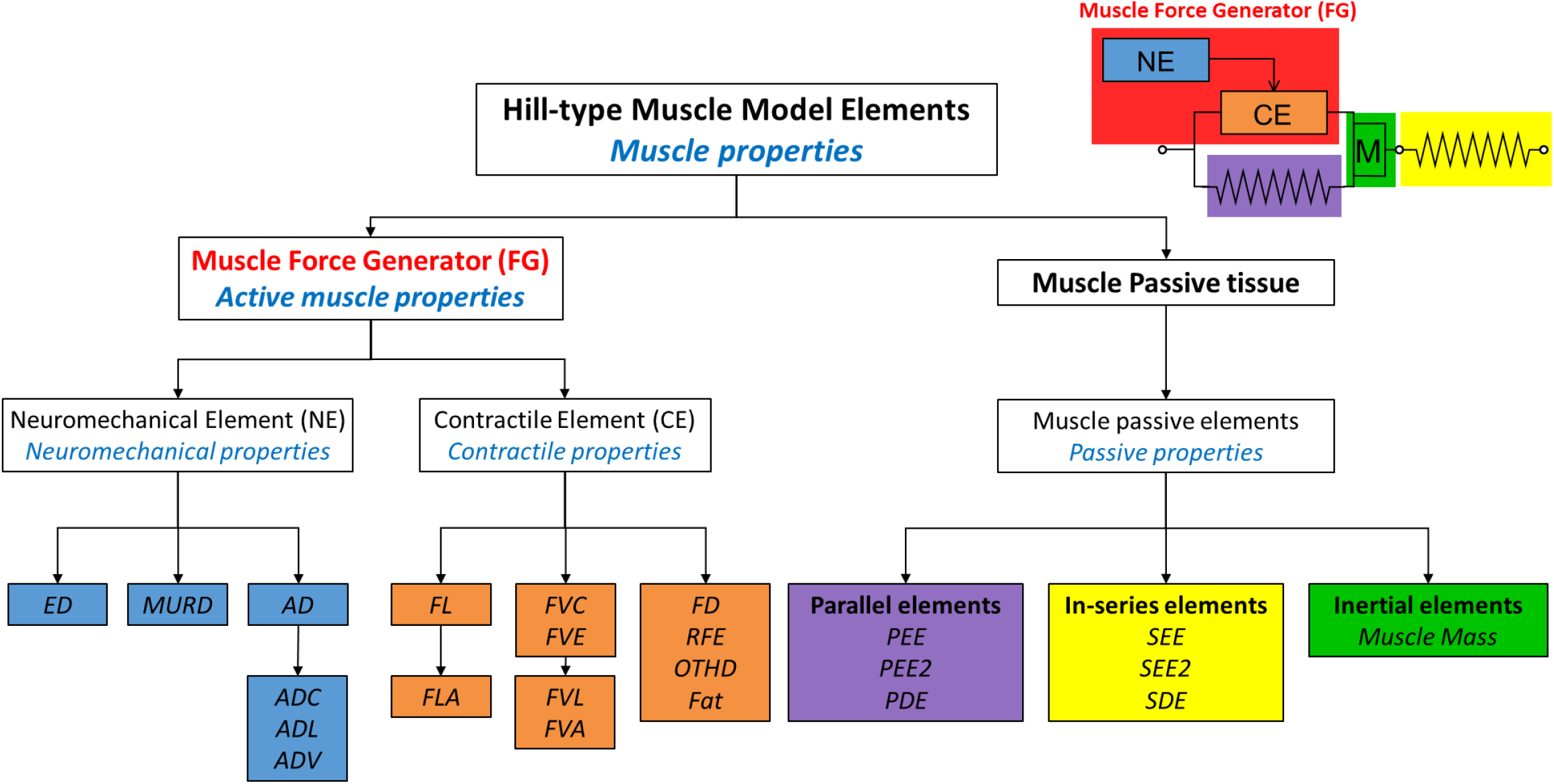
Hill-type models typically include passive elements and a muscle Force Generator (FG). In the FG, the Neuromechanical Element describes the muscle’s excitation-contraction coupling properties (light-blue boxes) and drives the muscle Contractile Element, described by the contractile properties (orange boxes) and responsible for force generation. Some properties are nonlinearly dependent to the level of activation (FLA, FVA), length (ADL, FVL) and contraction velocity (ADV) of the FG. Besides inertial elements (green box), Hill-type models include passive elements in-parallel (purple box) and in-series (yellow box) with the FG. The acronyms for the 9 rheological elements and the 23 neuromuscular properties are defined in Table 2.

In this review, the FG’s behaviour is defined by the interaction of the contractile element (CE) and the neuromechanical element (NE) (Fig. 2, Table 2). The CE lumps any of the 10 CE-related contractile properties (Table 2) that describe the multiscale mechanical events that occur during the active contraction dynamics of the FG. The NE groups the electrochemical processes involved in the neuromechanical interface between the central nervous system and the muscle machinery; it accounts for the excitation, activation, and motor unit recruitment dynamics of the neuromuscular system, that define the neural control to the FG.

For future purposes, a ‘baseline’ Hill-type model (Fig. 1) refers to any rheological arrangement of one passive element, one NE and one CE, the behaviour of which is respectively described by one passive property, one CE-related property, and one NE-related property. In the following, a Hill-type model is termed ‘innovative’ if it provided a methodological contribution to the field at the time of its publication, and more precisely a novel modelling approach for at least one of the 23 properties reported in Fig. 2, that could not be retrieved from a previously published paper identified from the systematic search. For conciseness and simplicity in this review, proposing novel rheological arrangements, new mathematical descriptions for previously published methodological approaches, and new ways of identifying the parameter values scaling those mathematical expressions were chosen not to be criteria to identify a model as ‘innovative’.

If not stated otherwise, this review only refers to ‘generic’ models [23], that are defined by properties normalized by the muscle-specific slack length of the Series Elastic Element (SEE), the optimal length of the CE, the maximum isometric force of the FG, and the maximum contraction velocity of the CE. For simplicity and conciseness, muscle pennation is disregarded in this review, despite limitations for highly-pennated muscles [23], [110]-[112]. Consistently with the literature, the passive elements and their respective constitutive property are described with the acronyms defined in Fig. 2 in the following.

### B. Systematic search for eligible studies

The research studies published between 1938 and 01/09/2024 that involved a Hill-type model were systematically searched for on four computational databases: PubMed, Web of Science, MedLine via Web of Science and EMBase via Ovid. Specific keyterms, listed in the Boolean search string in SM3, were searched for in the titles and abstracts of the research studies. The keyterms were grouped into four concepts (C1 to C4) listed in Table 1 with OR Boolean operators. The four groups of keyterms were then joined into a unique search string of the form ‘C1 AND C2 AND C3 NOT C4’. Details on the search strategy are provided in SM3. The results of the systematic search were stored by year of publication in the reference manager Mendeley.

The final set of ‘eligible’ studies retained for scoring assessment was identified according to six eligibility criteria (EC1 to EC6) listed in Table 1, aiming for milestone studies that provide detailed descriptions of ‘complete’ Hill-type models representative of the genesis and the state-of-the-art in Hill-type modelling. After removing duplicates, the studies that did not meet EC1 to EC5 were chronologically removed based on title screening, abstract screening, and if necessary, screening of the core text of the paper. The filtering was then repeated with the reference lists of the papers that passed EC1 to EC5, to identify further potentially eligible studies that were missed by the systematic search. The final set of studies that met EC1 to EC5 were chronologically compared for application of EC6. To identify the ‘innovative’ studies, inheritance diagrams (provided in Supplementary Material) were built for the AD, FL, FV, PEE and SEE properties to map over the past fifty years the chronological chain of references between studies. The inheritance diagrams distinguish the seminal studies that proposed novel modelling approaches to the field from the subsequent studies that adopted those new features or models.

### C. Scoring assessments

#### 1) Completeness assessment

Each eligible model was assessed for ‘completeness’ by scoring 1 for each of the 23 properties (Fig. 2) that was included in the Hill-type model, yielding a ‘completeness’ score out of 23, with a minimum of 3 according to EC4. According to Table 2, the modelling of the NE, CE and passive elements can respectively score 6, 10 and 7, yielding a balanced significance between these three types of elements. The completeness score enables an inter-study assessment of the level of complexity and the relative modelling strengths and weaknesses of the eligible models for mimicking the true muscle neuromechanical interface, mechanical events, and passive dynamics. The average score across eligible studies for each property was also calculated to identify the most common practices and gaps in Hill-type modelling.

#### 2) Modelling assessment

Four methodological practices (I-IV) considered by the reviewers to be crucial for a suitable transmission of incremental knowledge in the field of Hill-type modelling were reviewed in a ‘modelling assessment’ of the eligible studies. With the 10 questions (Q1 to Q10) listed in Table 3, it was evaluated the strategy taken by the eligible studies for

**Table 3:** The 10 questions constituting the modelling assessment. The eligible studies can score 0, 1 or 2 to these questions according to a detailed scoring scheme provided in SM4.

| Practices | Q# | Questions |
| --- | --- | --- |
| (I) Model Validation | 1 | Is the accuracy of the predictions obtained from the Hill-type model validated against ad hoc experimental data? |
|  | 2 | Are the Hill-type model’s predictions validated against reliable muscle-specific experimental quantities? |
|  | 3 | Is the accuracy of the Hill-type model evaluated using objective quantitative metrics? |
| (II) Model reusability | 4 | Is enough information provided to reimplement the Hill-type model? |
|  | 5 | Is the Hill-type model optimized for numerical stability and computational speed? |
| (III) Modelling choices and strategy | 6 | Are the simplifications adopted in modelling the muscle, both physiological and across relevant scales (e.g. sarcomere, fibre, entire muscle), clearly stated? |
|  | 7 | Are the modelling choices for modelling the rheological model clearly stated and / or justified with respect to the scope and objectives of the study? |
|  | 8 | Is there a discussion about the strengths and limitations of the normalized Hill-type model? |
| (IV) Model calibration | 9 | Are the mathematical expressions describing the properties of the generic (normalized) Hill-type model parametrized from experimental data consistent with the scope of the study? |
|  | 10 | Is the generic Hill-type model scaled (denormalized) with species- patient- or muscle-specific architectural scaling parameters? |

I. model validation, with a focus on quantitative and objective validation of the Hill-type predictions against reliable experimental measurements,
II. promoting model reusability, with a focus on the feasibility of model reimplementation and computational efficiency,
III. detailing the modelling choices, strategy, and limitations, with a focus on justified decisions for the taken physiological, multiscale, and rheological simplifications and modelling approaches, and
IV. model calibration, with a focus on how the constitutive properties and relationships of the generic (normalised) Hill-type model were obtained, and then scaled (denormalized) using species-, muscle-, and potentially subject-specific parameters.

Q1 to Q10 were scored 0, 1 or 2, which indicate no, incomplete and satisfying methodological practice respectively, and yield a ‘modelling score’ out of 20. To avoid bias and subjectivity, a detailed scoring grid was built for each question and is provided in SM4.

#### 3) Global score

For each eligible paper, the completeness and modelling scores were normalized by 23 and 20 respectively and averaged to yield a final global score in percentage. The global score measures the global degree of completeness, validation and reusability of an eligible model and the level of attention given to its modelling strategy and calibration. The global score is not an indicator of the overall quality of the eligible studies, considering that some of them were not modelling studies, with focus and contributions beyond the scope of the scoring assessments.

## III. RESULTS

### A. Systematic search for eligible studies and scoring process

The systematic search returned 57 eligible studies, listed in Table 4, that meet EC1 to EC6 (Table 1). As displayed in the flowchart in Fig. 3 (see SM5 for additional details), the three databases returned 2585 journal articles in English without duplicates, among which 127 were identified by the reviewers to meet EC2 to EC5. After screening the reference lists and identifying 38 additional studies that met EC2 to EC5, 165 studies were carefully screened in several iterations to apply EC6, using the inheritance diagrams stored in supplementary material as support. Fifty-seven eligible models met EC6 for the reasons listed in SM6, while the 108 excluded studies were either replicates or a combination of the 57 eligible models. As displayed in Fig. 3, 64 non-eligible studies were retained and listed and briefly analysed in SM7; 33 of them proposed models, also assessed for completeness in SM8, that were innovative (EC6) but did not meet EC4, while the 31 others brought significant contributions to the field of Hill-type modelling, including narrative reviews, sensitivity analyses, or comparisons between published models. The 57 eligible (listed in Table 4) studies and the 33 non-eligible models that met EC6 (listed in SM7) are considered to be representative of all the modelling approaches proposed to date in the field of Hill-type modelling for the properties listed in Table 2.

**Table 4:** List of the 57 eligible models and the primary Hill-type-related aim of their related study. Aims are either (1) Advancing the field of Hill-type modelling, (2) Developing methods for including Hill-type models in neuromusculoskeletal simulations, or (3) Investigating physiological mechanisms using Hill-type models. For conciseness, the studies are referred to ‘First Author Year of publication’ in this table and in the figures of this review.

| # | Study | Reference | Primary aim |
| --- | --- | --- | --- |
| 1 | Bahler 1968 | [118] | (1) |
| 2 | Hatze 1977 | [119] | (1) |
| 3 | Hatze 1978 | [120] | (1) |
| 4 | Wyss 1984 | [121] | (2) |
| 5 | Pierrynowski 1985 | [122] | (2) |
| 6 | Otten 1987 | [123] | (2) |
| 7 | Bobbert 1990 | [124] | (3) |
| 8 | Giat 1993 | [88] | (1) |
| 9 | Van Soest 1993 | [60] | (3) |
| 10 | Happee 1994 | [125] | (2) |
| 11 | Durfee 1994 | [126] | (1) |
| 12 | Winters 1995 | [56] | (2) |
| 13 | Caldwell 1995 | [42] | (3) |
| 14 | Van Zandwijk 1996 | [127] | (1) |
| 15 | Dorgan 1997 | [30] | (2) |
| 16 | Shue 1998 | [128] | (1) |
| 17 | Nussbaum 1998 | [129] | (1) |
| 18 | Curtin 1998 | [130] | (1) |
| 19 | Brown 1999 | [116] | (1) |
| 20 | Cheng 2000 | [31] | (1) |
| 21 | Ettema 2000 | [131] | (1) |
| 22 | De Woody 2001 | [132] | (3) |
| 23 | Gallucci 2002 | [133] | (3) |
| 24 | Lloyd 2003 | [26] | (1) |
| 25 | Thelen 2003 | [49] | (1) |
| 26 | McLean 2003 | [62] | (3) |
| 27 | Stelzer 2006 | [134] | (2) |
| 28 | Kistemaker 2006 | [135] | (2) |
| 29 | Günther 2007 | [136] | (1) |
| 30 | Mavritsaki 2007 | [137] | (1) |
| 31 | Gömmel 2008 | [138] | (3) |
| 32 | Song 2008 | [32] | (1) |
| 33 | Menegaldo 2009 | [139] | (1) |
| 34 | Moody 2009 | [68] | (3) |
| 35 | Rengifo 2010 | [140] | (3) |
| 36 | Proctor 2010 | [141] | (2) |
| 37 | Tsianos 2011 | [142] | (1) |
| 38 | Mörl 2012 | [48] | (1) |
| 39 | Wang 2012 | [143] | (2) |
| 40 | Blümel 2012 | [144] | (1) |
| 41 | Maceri 2012 | [145] | (1) |
| 42 | Callahan 2013 | [146] | (1) |
| 43 | Millard 2013 | [33] | (1) |
| 44 | Chand 2013 | [61] | (3) |
| 45 | Lee 2013 | [147] | (1) |
| 46 | Elias 2014 | [148] | (1) |
| 47 | Hamouda 2016 | [149] | (1) |
| 48 | Dick 2017 | [3] | (1) |
| 49 | Marcucci 2017 | [150] | (3) |
| 50 | Ross 2018 | [151] | (1) |
| 51 | Kim 2015, 2018 | [34], [117] | (1) |
| 52 | Siebert 2018 | [152] | (1) |
| 53 | Rockenfeller 2020 | [153] | (1) |
| 54 | Warner 2020 | [154] | (2) |
| 55 | Hussein 2022 | [155] | (1) |
| 56 | Millard 2023 | [35] | (1) |
| 57 | Caillet 2023 | [29] | (1) |

**Fig. 3.**
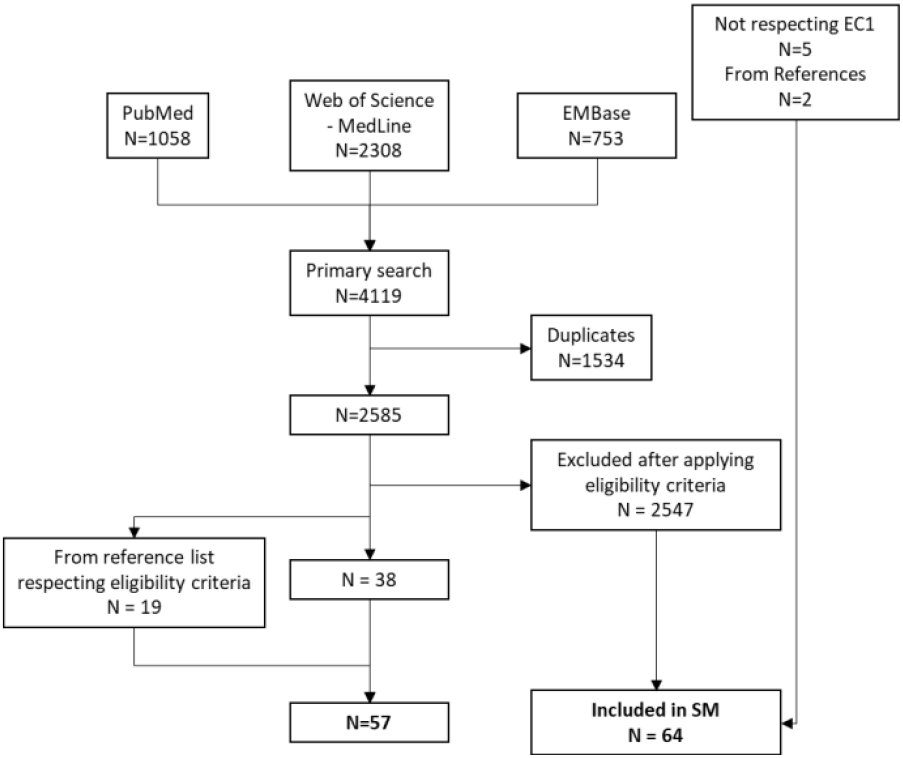
Flowchart of study inclusion according to EC1 to EC6. 38 eligible studies were obtained from the systematic search and 19 from the reference lists of the studies respecting EC1 to EC5. 64 non-eligible studies were listed in Supplementary Material 7 (SM7), 33 of which were assessed for completeness in SM8 for their important contribution to the field of Hill-type modelling. See SM5 for a more detailed version of this diagram.

As reported in Table 4, 35 of the 57 eligible studies primarily aimed to develop the field of Hill-type modelling by proposing innovative muscle models and investigating the sensitivity of model predictions to parameter values, challenging some modelling assumptions, or investigating the strengths and limitations of Hill-type models to predict some physiological mechanisms. Eleven eligible studies primarily aimed to developed new methods to use Hill-type models in neuromusculoskeletal simulations. Eleven eligible studies used Hill-type models to investigate physiological mechanisms including some material stress-strain dynamics, joint mechanisms related to muscle activity, posture and motion control in response to perturbations or the muscle redundancy problem. For clarity and conciseness, the studies are referred to ‘First Author Year of publication’ in the figures of this review and in Table 4. Also, the two eligible complementary studies [34], [117] were grouped together in this review.

The 57 eligible papers were scored for completeness and modelling approach, following the detailed scoring scheme in SM4, by two reviewers (A.C. and L.M.) independently. A high rate of agreement was obtained between scorers for the completeness assessment (binary scoring, 1219/1311 matching scores – 93% agreement rate). A fair rate of agreement was obtained between scorers for the quality assessment (ternary scoring, 382/570 matching scores – 67% agreement rate). Q3 and 4 about model reusability, and Q7 and 8 about the justification of the modelling choices strategy returned the highest rates of disagreement (36-49%) because they are more subjective than the other scoring questions. Only 2% of the quality scores showed strong disagreement (i.e., one scorer scores 0 while the other scorer scores 2). Any disagreement between reviewers was discussed while consulting the original study until a consensus was iteratively reached. The scoring spreadsheet is provided as supplementary material.

### B. Completeness assessment

#### 1) General results

The per-study results of the completeness assessment are displayed in Fig. 4 (additional details in SM8). The 57 eligible models scored in the range [3-16] out of 23 possible neuromuscular properties (mean: 8.1, standard deviation: 3.0) and showed no correlation between model completeness and publication year. As shown in Fig. 5(a), most of the models included between seven and nine neuromuscular properties, six models included five or fewer properties, consistently with the model description by Zajac [23], and eleven models included at least 11 properties [30]-[32], [35], [56], [116], [119], [142], [144], [148], [153], that is half or more of the identified muscle properties reported in Table 2. The specific properties included in each of the 57 eligible studies can be identified in the bar graph in SM8. The rheological arrangements of the eligible models are displayed in SM9.

**Fig. 4.**
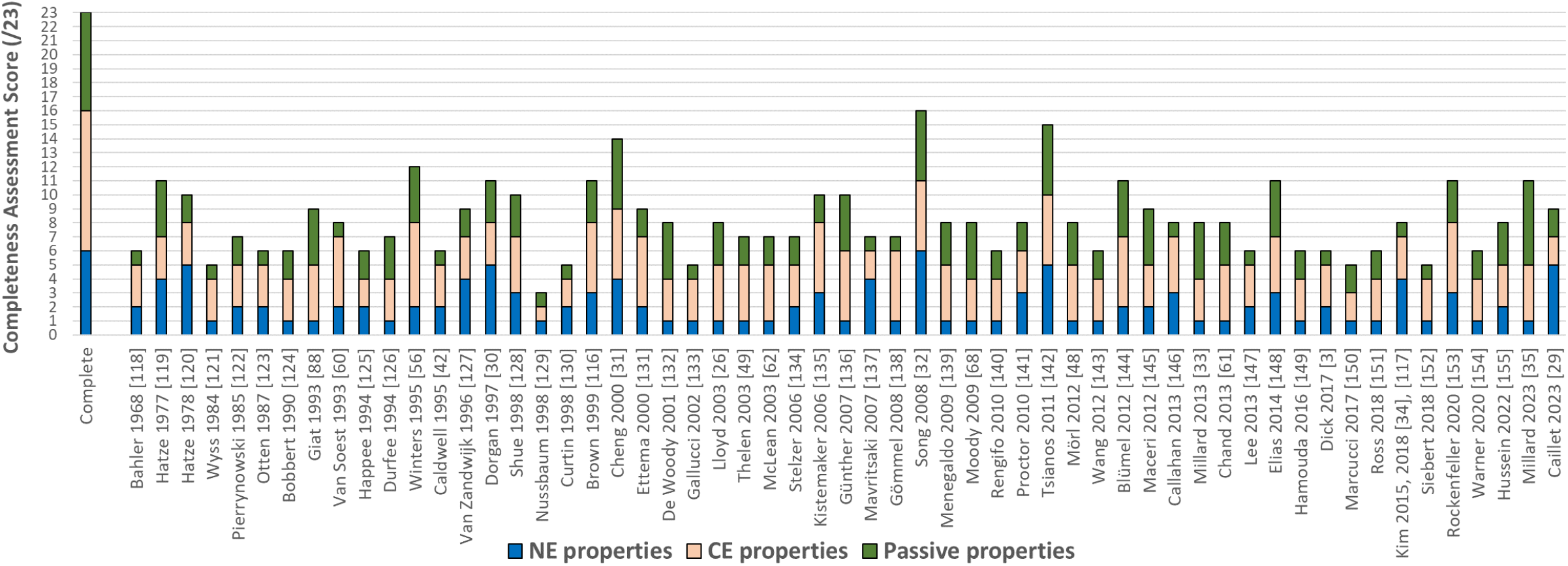
Results of the completeness assessment for the 57 eligible studies. For comparison, a ‘complete’ bar represents a theoretical complete model that would include the 23 assessed properties (see Fig. 2, Table 2). Blue: the 6 properties of the NE. Red: the 10 properties of the CE. Green: the 7 properties of the passive elements. The 57 studies are chronologically ordered. A more detailed version of this graph is provided in SM8.

**Fig. 5.**
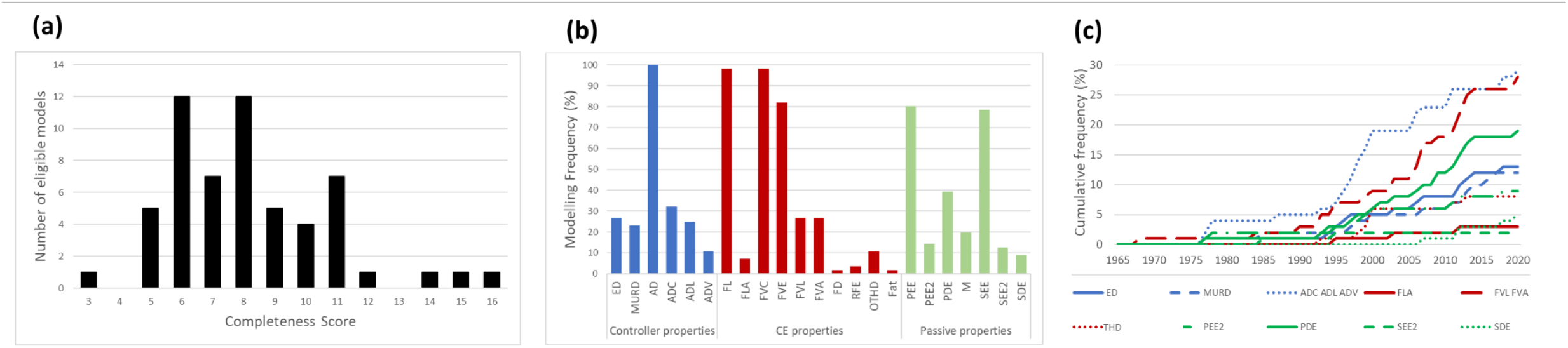
Results of the Completeness Assessment of the 57 eligible studies. (a) Frequency distribution of the eligible models according to their completeness score (out of 23). (b) Modelling frequency of each of the 23 properties (see Fig. 2, Table 2) among the eligible studies, where the modelling occurrence of each property is normalized to a percentage. (c) Time-accumulation of the eligible models that considered the 12 least commonly modelled properties. For clarity, some properties are grouped together: ADC-ADL-ADV, FVL-FVA, and FD-RFE-OTHD-Fat under ‘THD’

#### 2) Common modelling trends

The frequency graph in Fig. 5(b) shows that the six properties AD, FL, FVC, FVE, PEE and SEE are widely considered key in Hill-type modelling as they respectively appear in 98%, 96%, 96%, 81%, 79% and 77% of the eligible studies. These most popular properties were the first to be proposed in Hill-type modelling (see “Concise historical excursus of Hill-type modelling” in SM1) and have been described and extended in the past 50 years with various modelling approaches and mathematical descriptions, the time-evolution of which is detailed in SM10 with inheritance diagrams and a summary of the mathematical equations proposed in the eligible studies.

The dynamics of the NE are most commonly lumped with a phenomenological model of AD consisting of a transient first-order ordinary differential equation (ODE) inspired from [23], [56], [156], [157] in 31 of the 57 eligible studies (31/57), or with a steady-state nonlinear transfer function (7/57), mostly sigmoidal with advanced physiological interpretability [158]. Other eligible studies (19/57) proposed a more physiological modelling of AD and described the calcium dynamics [119], [120], [123] in an ADC approach. Furthermore, a quarter of the eligible models (15/57) drove the muscle’s AD with an excitatory drive derived from a model of ED, including for example feedback controllers (2/57), phenomenological ODEs (9/57) or comprehensive compartmental Hodgkin-Huxley-like models of the motoneuron discharging mechanisms [34], [141]. A quarter of the eligible models (13/57) considered the dynamics of motor unit recruitment (MURD) with, for example, detailed phenomenological models [30], [120], [146], simpler models relying on recruitment thresholds (4/57), or different recruitment characteristics for slow and fast representative units distinguished with EMG wavelet analysis [3], [147]. One eligible study [29], which proposes a muscle model with a motor unit resolution, used the discharge trains of individual motoneurons identified from experimental high-density EMG decomposition to physiologically describe the ED and MURD of the model and drive the AD of individual motor units, that constitute the motor unit pool of the muscle model.

The dynamics of the CE are dominantly modelled (55/57) with a length-dependent FL relationship that linearly scales with the active state, and is typically described with quadratic (11/57) [60] and gaussian (18/57) [119] mathematical descriptions according to SM10 (see [159] for a review). Two studies [35], [149] modelled active lengthening to address the issue of the non-physiological instability of the descending limb of the FL relationship [160]. Most eligible studies (39/57) modelled the FVC relationship with Hill’s two-parameter or normalized hyperbola (see SM10 for details), the 20% force overestimation of which near static conditions was addressed with a double-hyperbolic shape in one study [130]. Around a third (21/57) of the eligible studies modelled the FVE relationship with a two-parameter hyperbolic curve inspired from [161], [162], while less than a quarter (12/57) used an inverted hyperbola providing C2 continuity between concentric and eccentric FV branches.

The muscle and tendon passive properties are typically modelled with a PEE and/or a SEE, described as nonlinear passive springs, the stress-strain relationships of which are mostly defined with quadratic [60], [124], exponential [119], and piecewise linear-exponential [88], [123] expressions (see SM10). Approximately a third of the eligible studies (22/57) used a linear PDE to model the slight viscoelasticity of the muscle’s inner passive structures, or to prevent numerical singularities when inverting the FV relationship.

#### 3) Less commonly modelled properties

Beside the ten aforementioned properties (ED, MURD, AD, ADC, FL, FVC, FVE, PEE, SEE, PDE) that appear in at least 25% of the eligible studies, 20-25% of the eligible studies also consider muscle mass (11/57), accounting for muscle-specific inertial characteristics and preventing numerical instabilities [31], [32], as well as the nonlinear length- or velocity-dependency of the ADL (14/57), FVL (14/57) and FVA (15/57) relationships (see Fig. 5(b)). The nine remaining properties listed in Table 2 (ADV, FLA, FD, RFE, OTHD, Fat, PEE2, SEE2 and SDE) are rarely modelled and appear in less than 15% of the eligible studies (see Fig. 5(b)). As displayed in Fig. 5(c), most of the least commonly modelled properties have not been considered in the eligible models in the past ten years and were mostly developed in the 1990s and 2000s.

#### 4) Model parameters

Most eligible studies maintained low model complexity, as half used between 5 and 14 parameters to define the neuromuscular normalized properties, a third used 15 to 24 parameters, and 6 studies used more than 35 parameters up to around 60. As supported by SM8, on average two parameters were used to define one property, with higher ratios for the studies modelling the MURD [120] or comprehensive descriptions of the NE [34], [141].

### C. Modelling assessment

#### 1) Global results

Fig. 6(a) shows that the eligible studies scored in the range [3-19] out of 20 (mean: 11.5, standard deviation: 4.4), 18 studies scored less than 10/20, and 10 studies scored above 15/20. The per-study results of the modelling assessment are displayed in Fig. 7 (additional details in SM11) and show no correlation between modelling score and publication year. As shown in Fig. 6(b), the four scoring categories (validation, reusability, modelling choices and strategy, and calibration) detailed in Table 3 globally received a similar level of attention from the eligible studies with average scores of 1.0, 1.2, 1.2 and 1.2 respectively. The eligible studies scored on average above 1.0 for all questions, except Q3 for model validation (0.7/2), Q5 for model reproducibility (0.7/2), and Q6 for clear model assumptions (0.9/2). SM12 reports for Q1 to Q10 the most popular modelling approaches proposed in the eligible studies.

**Fig. 6:**
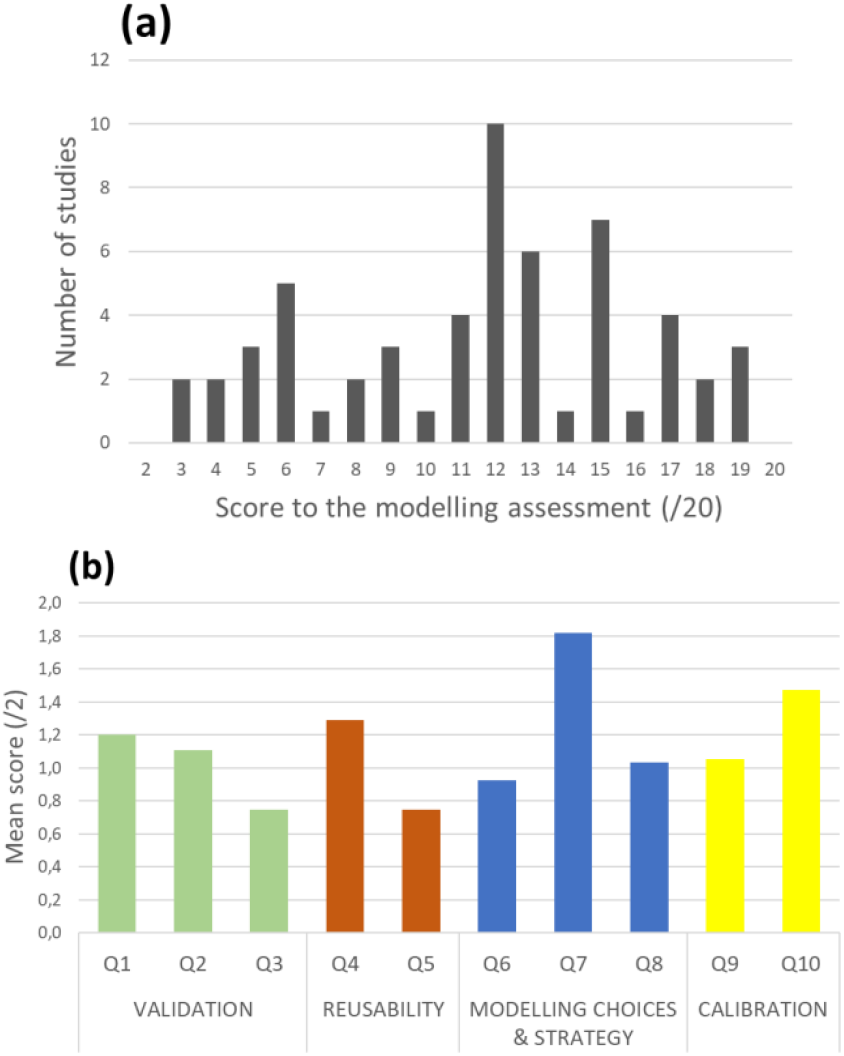
(a) Distribution of the 57 eligible studies according to their total score in modelling assessment (out of 20). (b) Mean scores (out of 2) among the 57 eligible studies for the 10 questions (listed in Table 3) of the modelling assessment.

**Fig. 7:**
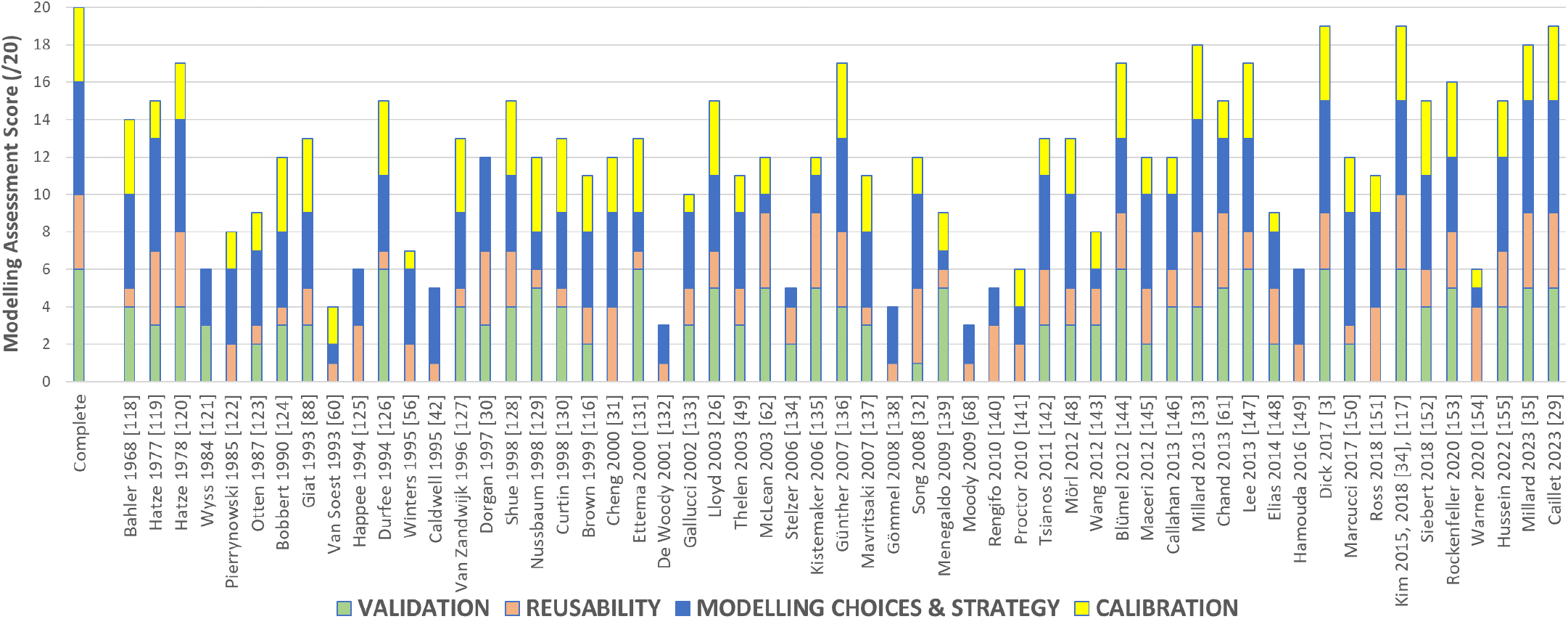
Results of the modelling assessment for the 57 eligible studies. For comparison, a ‘complete’ bar represents a theoretical model that would score 2 to the 10 assessment questions listed in Table 3 and detailed in SM4. Green: model validation. Red: model reusability. Blue: modelling choices and strategy. Yellow: model calibration. A more detailed version of this graph is provided in SM11.

#### 1) Model validation (Q1 to Q3)

Q1&2-Less than half of the eligible studies (25/57) validated the model predictions against ad hoc experimental animal data, mostly mammalian muscle force (10/57) or human (16/57) quantities such as joint torque (12/57). A quarter (17/57) of the studies proposed validations against results from available experimental datasets, the protocol of which were reproduced in simulations, and the remaining eligible studies (15/57) either proposed no model validation or qualitative validations against other results from the literature. Q3-Less than a third (16/57) of the eligible studies used mathematical or statistical metrics for the analysis and interpretation of the model predictions, while the remaining studies (38/57) only proposed qualitative and visual assessments of their analysis. Consequently, approximately a third of the eligible studies proposed sufficient validation for a satisfactory evaluation of the performance of their Hill-type muscle model.

#### 2) Model reusability (Q4 and Q5)

Q4-Approximately half (25/57) of the eligible studies did not provide enough information to make possible a full re-implementation of their Hill-type models and a reproduction of the published results. The non-reproducible models were missing some reference or some description for the chosen rheological arrangement, the mathematical equations describing the modelled properties, and/or the parameter values scaling these properties. Out of the 32 remaining eligible studies, which presented models that could theoretically be re-implemented, only six provided full access to the actual implementation of their computational model [29], [31]-[35], while open-source versions for two more [49], [61] were then made available as the result of other studies. Q5-Most of the eligible models (30/57) did not report any optimization for stability and computational speed, while eight imposed C1 or C2 continuity for all modelled properties, and twelve avoided singularities in the inverted FV relationship (see SM10 and SM12 for additional details). Consequently, most of the eligible models cannot be readily reused in other studies.

#### 3) Modelling choices and strategy (Q6 to Q8)

Q6-More than half of the studies (36/57) commented on some modelling simplifications and assumptions regarding the muscle physiology and geometry, while around a third (23/57) discussed the multiscale modelling approach typical of Hill-type modelling, i.e. the assumptions required to develop a consistent model across sarcomere, fibre and muscle scales. Q7-Around two thirds of the eligible studies (34/57) justified their modelling choices with references and in accordance with the scope of the study. Q8-However, half of them (29/57) did not discuss and compare the strengths and weaknesses of the modelling approach in relation to other studies in the literature. Consequently, most of the eligible studies do not attempt to draw conclusions on the future works necessary to improve their model.

#### 4) Model calibration (Q9 and Q10)

Q9-Half (27/57) of the eligible studies parametrized at least one of the constitutive properties of their generic (normalized) Hill-type model with subject-or muscle-specific values, that were mostly obtained with parameter calibration minimizing a cost function between simulated and experimental quantities, or from ad hoc experimental measurements. Q10-Half of the eligible studies (27/57) used similar methods to scale the generic Hill-type model with subject-specific architectural values for the slack length of the Series Elastic Element (SEE), the optimal length of the CE and the maximum isometric force of the FG. Consequently, most of the eligible models remained either generic, were not scaled to subject- and muscle-specificity, or were parametrized with species-inconsistent values from the literature, e.g., AD relationship derived from measurements in frog fibres, FV relationship scaled with cat gastrocnemius data, and maximum isometric force extracted from literature human cadaveric data.

### D. Global assessment

Fig. 8 provides the global score for the 57 eligible studies. A high score globally suggests that a model includes numerous muscle properties, is carefully described, calibrated, and validated, and is easily reusable. However, the global score is *not* an indicator of the overall quality of the eligible studies, some of which are not focused on Hill-type modelling. The eligible studies scored in the range [25%; 69%] (mean: 45%, standard deviation: 14%). Fourteen eligible studies scored below 33%, 19 in the range [33%; 50%], and fourteen above 60% [3], [29], [31]-[35], [119], [120], [128], [136], [142], [144], [153].

**Fig. 8.**
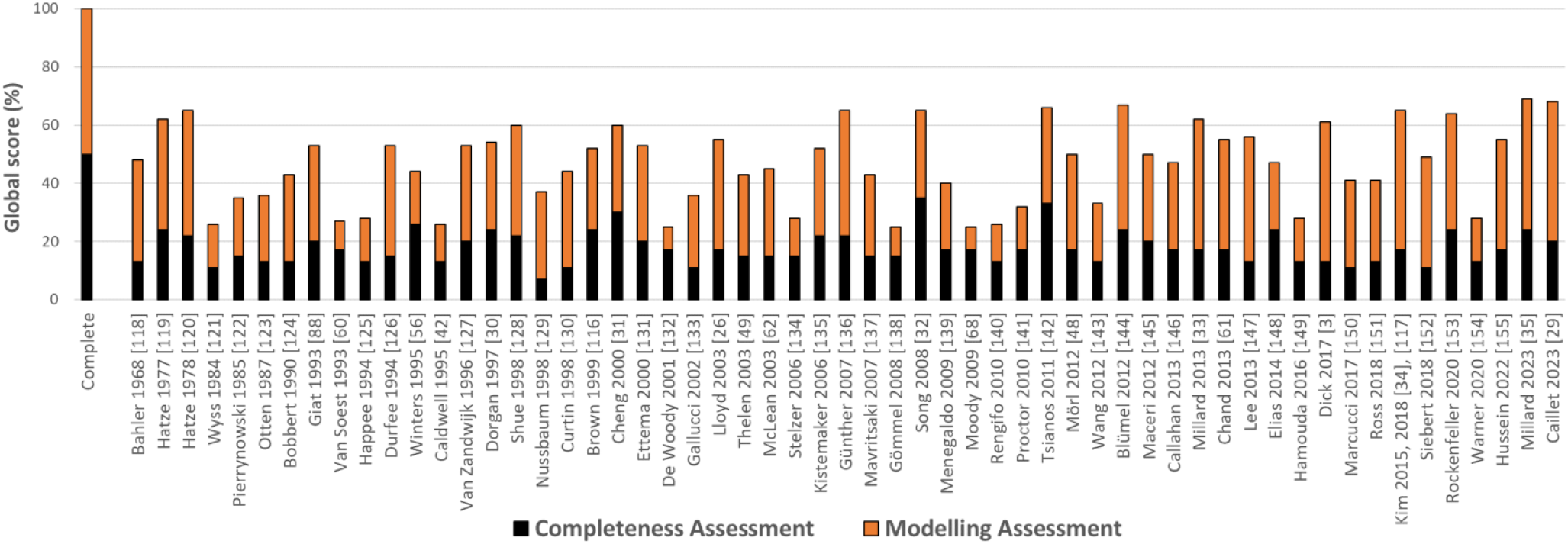
Global scores for the 57 eligible studies. Completeness (black) and modelling (orange) assessments were given the same weights to derive the global score.

## IV. DISCUSSION

### A. Summary and main objectives

This systematic review clarifies the dense and hard-to-explore field of Hill-type modelling, for which recent reviews are lacking, and provides a tool for researchers to identify the current trends in Hill-type modelling and to assess the state-of-the-art and current gaps in property modelling and methodological practices. To meet these objectives, 57 eligible studies (Table 4), which were extracted from the systematic search (Fig. 3) according to six eligibility criteria (EC1 to EC6, Table 1) that impose a minimum level of model complexity (Fig. 1) and innovation (SM6), were investigated according to two scoring schemes. In the following, the results of the completeness and modelling assessments (Fig. 4 to Fig. 8) are used to (1) define the main trends and gold standards in Hill-type modelling (Section IV.C.), (2) challenge these gold standards with advanced modelling methods (Section IV.D.), (3) clarify the state-of-the-art by investigating properties non-commonly modelled in the field and the current gaps in the field (Section IV.E.), (4) discuss the importance of model validation (Section IV.F.), and (5) open the discussion on the necessity of more regular reviews and global recommendations in the field of Hill-type modelling, with an emphasis on model reusability for optimal transmission of knowledge (Section IV.H.).

### B. How to use this study?

Besides clarifying the contents and current challenges of the field of Hill-type modelling for non-specialists, this study provides, with Fig. 4, Fig. 7 and Fig. 8 (with more details available in SM8 and SM11), a simple tool to researchers to guide their choice towards the best available published Hill-type models to address the aims of their study with respect to model completeness, validation, reusability, or calibration, and modelling choices and strategy. Using as support the list of published reviews, sensitivity analyses and comparative studies in SM7, modelers can also take inspiration from the following discussion and the extensive supplementary material provided in this review on the current trends and state-of-the-art in modelling approaches and methodological practices to develop new models to address the current gaps in the field. The set of 90 (57 eligible (Table 4) and 33 non-eligible but innovative (SM7)) Hill-type models reported in this study is representative of all the modelling approaches proposed in published Hill-type structures for modelling the properties listed in Table 2 and displayed in Fig. 2. Detailed and didactic reviews of the rheological structures proposed in the field, and of the main modelling approaches for properties ADC, AD, FVC, FVE, FVL, FVA, FL, FLA, PEE and SEE are proposed respectively in SM9, and in SM10, with lists of mathematical equations and inheritance diagrams. The properties modelled by the 90 studies can be identified with the detailed bar graphs in SM8 resulting from the completeness assessment, while the model complexity can be inferred from the number of model parameters in SM8. The reader interested in open-source solutions is referred to the relevant studies identified in this review [29], [31]-[34], [137], to the GitHub repository https://github.com/modenaxe/awesome-biomechanics, and to the open-source implementation of the models used in the OpenSim platform [37], [38].

### C. Most common trends and gold standards in Hill-type modelling

According to the conclusions drawn in the Results Section from the completeness and modelling assessments, and to Fig. 5(b), Fig. 6(b) for Q6-8, and SM12, Hill-type models are most commonly described with a Voigt-Kelvin structure that includes a NE, a CE, a PEE and a SEE (see examples of rheological structures in SM9) to model the ‘classic’ AD, FL, FVC, FVE, PEE, SEE properties in a multiscale approach where a representative muscle fibre represents the whole muscle. Typically, a single phenomenological ODE produces a time-history-dependent active state *a*(*t*), that describes the lumped activity of the muscle’s neuromechanical interface. Usually, the dynamics of the NE and the CE are simplified to be decoupled (which is physiologically untrue), so that the active state *a*(*t*) linearly scales the instantaneous length-dependent and velocity-dependent force factors *f*_*FL*_ (*l*^*FG*^) and *f*_*FV*_ (*v*^*FG*^) determined by the independent CE’s FL and FV properties. Finally, the total FG force *f*^*FG*^ is calculated as *f*^*FG*^(*t, l*^*FG*^, *v*^*FG*^) = *a*(*t*) · *f*_*FL*_ (*l*^*FG*^) · *f*_*FV*_(*v*^*FG*^). The FG, in-parallel with the PEE, is in equilibrium with the SEE, which passively transmits the active force as *f*^*SEE*^= *f*^*FG*^ + *f*^*PEE*^. These equations are typically rearranged to isolate and solve for unknown state variables, such as the FG length *l*^*FG*^ or force *F*^*FG*^. These usual multiscale rheological arrangements and equations directly result from gold standard modelling approaches proposed between 1938 and 1976 (see SM1 for a historical excursus), that define a common baseline for most of the Hill-type models in the contemporary literature. The most common mathematical descriptions for the AD, FL, FV, PEE and SEE properties are reported in SM10 and their evolution with time is mapped with inheritance diagrams.

### D. Challenging the gold standards in Hill-type modelling

The aforementioned gold standards in Hill-type modelling (Section IV.C.) can be challenged and the most common trends in defining and modelling the multiscale approach, the FL and FV properties, the active state, the interplay between the NE and the CE and the in-series passive elements can be extended.

#### 1) Multiscale approach

As previously mentioned, Hill-type models are flexible enough to describe, with a single actuator and adequate multiscale simplifications, the structure and dynamics of either individual sarcomeres, fibres, motor units or, most commonly, whole muscle-tendon systems. Despite successfully reducing the cost and complexity of Hill-type muscle modelling, multiscale approaches come with numerous important assumptions. When a whole muscle is described as one of its ‘representative’ unidirectional fibre or motor unit, it implies that all muscle fibres or motor units have homogeneous ‘average’ material properties and dynamics, and are in parallel, coplanar, of the same length spanning across the whole muscle length, oriented with the same pennation angle with respect to the tendon and simultaneously excited with the same neural control and electromechanical delay with an homogeneous repartition of activated fibres across the muscle, and work independently without lateral force transmission. Multiscale sarcomere-to-muscle approaches further imply that the muscle’s sarcomeres all have the same length and neuromechanical characteristics and work independently from their in-parallel neighbours. These multiscale simplifications, albeit necessary, may contradict experimental data ([163], [164], for example), prevent modelling of some muscle properties, such as MURD, and affect the physiological accuracy of the muscle description with limitations in reconciliating microscopic and macroscopic scales and mechanisms [165]-[168], with subsequent possible downgrades in prediction accuracy [169].

Despite these important limitations, only a third of the eligible studies (see SM11 and Fig. 6(b), Q6-7), including the eleven studies that model single sarcomeres, fibres, or motor units with a Hill-type model [29], [34], [117], [119], [120], [127], [137], [146], [170], explicitly acknowledged the multiscale approach and discussed its implications on Hill-type modelling. This lack of clarity feeds and is strengthened by a lack of consensus between studies in the notation defining the muscle quantities, with 22 different notations observed in the eligible studies for the length of the contractile unit (*L, L*_*M*_, *L*_*CE*_, *L*_*m*_, *L*_*f*_, *L*_*fibers*_, *L*^*M*^, *L*^*m*^, *L*^*F*^, *l, l*_*CE*_, *l*_*CC*_, *l*_*m*_, *l*_*f*_, *l*_*ce*_, *l*^*M*^, *l*^*CE*^, *X*_*m*_, *x, x*_*eye*_, *x*_*CE*_, *λ*), and often the same notation (e.g., *l* or *l*_*m*_) for different quantities (e.g., muscle and muscle-tendon length). Overlooking the multiscale assumptions, the eligible studies often used experimental data (e.g., measurements on single fibres) to fit the Hill-type properties at a scale inconsistent with the modelled physiology (e.g., muscle scale). This is an important limitation, as the FL relationship is, for example, known to be piecewise linear at the sarcomere scale but to be smoother with a larger plateau [24] and a more slowly decreasing descending limb beyond optimal fibre length at larger scales [171], a multiscale difference sometimes debated [167], [172]. Also, the curvature of the FV relationship decreases with higher scales (compare [173]-[175] for works on rat skinned fibres, bundles of fibres, and whole muscles), while the Young’s modulus in the exponential stress-strain relationship of the PEE nonlinearly increases from the fibre to the muscle scale [176], with dominant contributions to the total passive force of the titin structures and of the extracellular matrix at the fibre and muscle scales respectively [176]-[178]. Consequently, a multiscale approach to Hill-type modelling must be correctly applied and commented to maintain credible descriptions of the muscle physiology and dynamics.

#### 2) FL, FV and FVL properties

Current gold standards in the modelling of the FL and FV properties can be advanced. The non-physiological instability [160] in the classic description of the descending limb of the FL relationship [179] can be corrected with a time-history-dependent approach accounting for the initial length at activation onset [149], a mechanistic model explicitly including the contribution of titin [35], or with a two-mode Hill-type model [180], [181]. Recent advances on our understanding of the active FL curve can be considered in future models, such as the usually overlooked important role of the length-dependent interfilament spacing and electrostatic interaction on the ascending arm of the FL relationship [182]. To avoid an overestimation of instantaneous force at low shortening velocities, the FVC relationship should be modelled as double-hyperbolic [130], [183]-[186]. Also, the standard models of FVE that reach forces higher than maximum isometric force during eccentric contractions are experimentally challenged [187]. As modelled in only a few eligible studies (Fig. 5(b)), the FVC property moreover nonlinearly varies with instantaneous muscle length [188] (FVL): the maximum shortening velocity decreases with shorter sub-optimal muscle lengths [118], [179], [189], despite a lack of consensus [184], [190], and the curvature of the FVC relationship (defined by the *a*_*f*_ parameter in SM10) decreases with shorter sub-optimal muscle lengths [191] and increases for longer over-optimal lengths [184]. The latter suggests an increased probability of cross-bridge attachment during filament sliding when the inter-filamentary spacing is reduced, which can be debated to be redundant with the properties described by the FL relationship.

#### 3) Definition of the muscle active state a

The active state of an FG is commonly defined as the intracellular concentration of calcium ions [49] or the proportion of available troponin receptors bound to calcium ions [192]. These definitions prevent extending the phenomenological description of the NE behaviour to subsequent important transient events that are typically not reflected in the instantaneous CE properties, such as the calcium-troponin binding dynamics or the probabilistic cycling mechanism of cross-bridge formation between detached, weak bound and force-generating states [18], [193], [194], that are typically overlooked in Hill-type modelling. We suggest defining the active state of an FG as the normalized instantaneous ratio of formed cross-bridges in the force-generating state to a representative population of myosin heads, irrespective of the degree of filamentary overlap, relative inter-filamentary sliding speed and other external mechanical mechanisms, which should be modelled in the contraction dynamics of the FG. This definition is compatible with any influence that fibre length or velocity can have on the mechanisms involved in the overall AD discussed below. To the authors’ knowledge, only three analytic [6], [195], [196] and five Hill-type muscle models [114], [123], [137], [197], [198] have proposed phenomenological time-dependent ODEs to explicitly account for the transient mechanisms that chronologically occur between the events of calcium ion release and force generation. These neuromechanical events are otherwise either neglected in Hill-type models or lumped in a steady-state pCa-force sigmoidal relationship [26], [119].

#### 4) Interplay between the NE and the CE: FLA, ADL, FVA and ADV properties

The neuromechanics of real muscles rely on an interplay between the CE’s and NE’s activation, length, and velocity states. However, this interplay is commonly overlooked in Hill-type models as the FLA, ADL, FVA, and ADV properties, where the FG properties are nonlinearly related to the active state *a*, length *l*^*FG*^ and velocity *v*^*FG*^ as described in SM10, are rarely included in Hill-type models (see Fig. 5(b, c) and SM8). Modelling the NE’s and CE’s behaviours to be independent, linearly scaling the FL and FV relationships with the active state (see Section IV.C.), and deriving the FL and FV properties from experimental observations at maximal muscle activity, are important limitations which partly explain the poor performance of Hill-type models in submaximal activation tasks [3], [33], [199].

FLA and ADL - The classic FL relationship is empirically observed [115], [200]-[205] and mechanistically explained [182] to be nonlinearly dependent to active state and to distort for submaximal contractions towards peak forces occurring at longer CE lengths. This nonlinearity is explained, at the CE (mechanistic) level, by changes in the force transmission pathways between the multiscale layer of the muscle [206] to minimize the muscle internal work and optimize force production [207]. The FLA property is phenomenologically captured in four eligible models [26], [29], [56], [144] with an activation-dependency of the FG’s optimal length 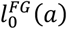, which improves the accuracy of the model predictions [208]. However, it is believed that the FLA approach phenomenologically encapsulates, in these studies, some ADL properties which truly occur at the NE (neuromechanical) level [114] and that are responsible for the nonlinear length-dependencies of the experimental frequency-force [116], [203], EMG-force [200] and pCA-force [209] relationships. In this respect, it is proven that the calcium transients are length-dependent [113], [210]-[212] as the muscle geometry affects the dynamics of calcium release from the sarcoplasmic reticulum and calcium diffusion in the sarcoplasm. The reaction rate for calcium-troponin (CaTn) unbinding decreases with increase sarcomere length [213], causing slower rate of activation decrease. Furthermore, lengths above 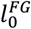 increase the sensitivity of cross-bridge formation to calcium ions [205], [209], [214], [215], and the probability of cross-bridge formation by geometrically reducing the intrafilamentary spacing between myosin and actin filaments [216]. Importantly, the latter phenomenon may be redundantly modelled with the FVL approach described previously. One eligible study modelled the length-dependency of some of these ADL properties (the transient concentrations of free calcium ions and CaTn complex) using experimental mammalian data [29]. Additional work should be achieved to include in Hill-type models those experimental evidence of length-activation nonlinear interdependency as new FLA and ADV properties, the possible redundancy of which should be carefully addressed.

FVA and ADV - The modelling of a nonlinear dependency on active state of the classic FVC relationship (FVCA) proposed in 14 eligible studies is justified by the important decrease in maximum shortening velocity with decreasing activation observed in human joint torques [217], [218], cat dissected muscles [219] and in rabbit skinned fibres [220] and explained by an increase in inactive inertia [221]. Yet, this phenomenon could also be modelled as an ADV property in the NE because of both a lower ratio of fast-contracting motor units at low activation, and an increase in the number of cross-bridges bearing tensile force against shortening dynamics, as cross-bridge detachment rates slow with lower levels of sarcomeric calcium concentration. While denied [217] or neglected with suitable methodology [56], three eligible studies [56], [136], [144] consider another FVCA property, where the curvature of the FVC relationship for a muscle decreases at higher activation [46], [219], which is potentially explained by the lower ratio of slow versus fast recruited motor units, which exhibit different FV contractile properties. Five eligible studies [31], [32], [116], [142] proposed an ADV property to model the yielding phenomenon, which relies on the same physiological mechanisms as short-range stiffness (SRS) [222] during eccentric contractions. Upon stretch deformation, cross-bridges provide rapid tensile stiffness [223], [224] before forcibly detaching leading to the force drop related to yielding [187], the effects being emphasized in slow-type fibres [116], [222] in fast eccentric contractions [222], [225], where the rates of cross-bridge detachment [226] and attachment are respectively higher and slower. Therefore, the SRS dynamics should be modelled as a passive property [227], while yielding is both a passive and a time-history dependent FVA property, for the decrease of weak-bound and force-generating cross-bridges respectively due to mechanical stretches. Although SRS and yielding should not be modelled in the NE according to the definition of AD previously proposed, they influence the time-course of the muscle ED and MURD [187], [228].

#### 5) In-series elastic elements

Few efforts were observed for challenging the gold standard models of the in-series elastic elements in Hill-type actuators. Yet, tendon slack length and stiffness have a key influence on model predictions, according to the sensitivity studies reported in SM7. The tendon behaviour is commonly modelled with a SEE in-series with the FG (see SM9 for rheological structures) and simple nonlinear stress-strain relationships (see equations in SM10). Besides complex multiscale tendon models [145], advanced phenomenological models of the tendon behaviour, that would for example account for its hysteresis [229], have however not yet been proposed in Hill-type models. Besides the tendon, most eligible models describe the additional passive activity of the muscle aponeurosis and other sub-scale passive structures with a SEE2, although their behaviour is non-intuitive [230] and they should be considered in-parallel with the FG and included in the PEE in first approximation [229]. In sarcomere-scale Hill-type structures, a SEE2 can however encapsulate the passive titin and cross-bridge elasticities, which are in-series with the force-generating cross-bridges. One eligible study proposes an advanced rheological architecture of PEEs and SEEs, some of which are activation-dependent, to model the stretching dynamics of titin in a representative sarcomere [35]. Importantly, at the muscle scale, titin and cross-bridge elasticities are mechanically not in-series with the macroscopic tendon due to the in-parallel arrangement of laterally-connected myofibrils and fibres, contradicting some previous sarcomere-to-muscle multiscale modelling approaches [23], [85], [122]. To avoid such inaccurate simplifications in sarcomere-to-muscle multiscale approaches, it is advised to neglect the SEE2 dynamics when the elasticity elastic energy storage is predominantly taken by a long and/or compliant free tendon [23]. The increment in tendon velocity-dependent stiffness arising from the SDE viscoelasticity can be likewise neglected for slow-to-medium contraction speeds [48], [231], [232], although at muscle level they have been reported to improve model stability [136], [233].

### E. Current gaps in Hill-type modelling and lines of improvement

According to the results of the completeness assessment in Fig. 4 and Fig. 5(a), the eligible studies globally score relatively low (mean and median of 8/23) and do not extend the baseline approach (Fig. 1) much beyond the gold standards discussed in Section IV.C.. Individually, the eligible models however tackle numerous gaps in Hill-type modelling and open the field to new lines of improvement, some of which were discussed in the previous section IV.D..

#### 1) Modelling the neuromechanical interface

Most eligible studies limit the modelling of the neuromechanical interface (i.e., the NE’s excitation-contraction coupling properties) to a ‘black box’ first-order ODE and/or a steady-state sigmoid transformation (see SM10 for examples) to transform the excitation state into the active state of the FG. These models of AD are hardly relatable to the real physiological mechanisms and are therefore difficult to interpret, calibrate, validate, and advance [234]. This is a major limitation when, in the actual biophysical system, numerous highly nonlinear mechanisms define the neuromechanical dynamics of the motor unit pool and the neural drive to muscle, and are proved to have a dramatic influence on force generation [235], [236]. To address this limitation, more complex modelling approaches must be proposed to accurately describe the NE dynamics, from the spinal synaptic influx to the muscle activation.

ADC - A first set of possible modelling advances relates to the ADC property, that was proposed in some eligible studies (18/57), which explicitly modelled the time-course of the calcium transients in the sarcoplasm (see SM10 for equations). By describing actual physiological mechanisms, this ADC approach enables a confident modelling of the length-dependency of the AD dynamics, discussed previously, and a direct validation against experimental [237], [238] or simulation [239], [240] studies evaluating calcium transients and pCa-force relationships. When modelling ADC, it is worth noting the preferable choice of the mag-2-fura indicator in experimental studies, as well as the strong dependency of the calcium transients to animal species and temperature [210], [241]-[249], as recently reviewed in the supplementary material of [29]. While some comprehensive models of calcium diffusion [194], [250], [251] can also be included in Hill-type models, additional phenomenological descriptions of the cascading AD dynamics, including the CaTn transients, can also be proposed [29].

MURD - A second set of possible modelling advances relates to the ED and MURD properties. As demonstrated in Fig. 5(b), because of the limitations related to the fibre-to-muscle multiscale approach, most eligible studies disregard the dynamics of MURD and Henneman’s size principle [252]-[254] in their model. In real muscles, the recruitment, discharge and mechanical properties of motor units are inter-related, motor unit-specific, and continuously distributed across the motor unit pool from small low-to large high-force and - threshold motor units, as quantitatively reviewed [255], [256], and the relative importance of MURD and discharge frequency in the determination of the neural drive to muscle is task-dependent [235]. Conversely, in most eligible studies, it is unclear whether the complete motor unit pool is assumed to be recruited at all times and the excitation state is an indicator of the overall motoneuron discharge frequency, or if the excitation state, typically obtained from processed EMG signals in this case, is a lump representation of the motor unit pool recruitment and discharge dynamics. To avoid this confusion, some studies proposed to phenomenologically model the MURD in a single multiscale NE for the whole muscle [24], [30], [32], [120], [142], [146], [170], [257], [258]. A few recent studies proposed a more physiological description of MURD by developing muscle models built as a collection of motor unit actuators, that are either modelled analytically [259], [260], with Hill-Huxley-type structures [261] or with Hill-type actuators [29]. In these motor-unit-scale models, the recruitment dynamics of the motor unit elements are usually modelled after the phenomenological approach from Fuglevand et al. [5], like in [259], [261], while other approaches could be also considered [89], [92], [262]-[266]. Comprehensively modelling the motor unit pool makes it easier to describe the continuous exponential distributions of the motor unit neuromechanical properties across the pool (e.g. force recruitment threshold, maximum isometric force, electrophysiological motoneuron properties) [256]. Another solution is lumping the motor unit pool and its MURD in a two-FG Hill-type model for slow-type and fast-type populations [3], [27], [147].

ED - The growing number of advanced models of MURD motivate advancing the modelling of the ED, i.e., of the electrical-chemical mechanisms that underly the time-course of the motor unit-specific neural message, from the spinal synaptic influx to the depolarization of the t-tubules. In the literature, analytical, phenomenological or Hodgkin-Huxley-like approaches of varying complexity are proposed to model the discharge trains of motor unit action potentials that drive motor unit-scale Hill-type models [34], [117], [119], [120], [127], [137], [141], [170]. This approach, using discrete action potentials as input, enables predicting individual force twitches that fuse into a realistic estimation of motor unit force, and a realistic reconstruction of muscle force by summing these motor unit forces. In contrast, muscle-scale actuators, such as EMG-driven models, rely on a single continuous neural input, yielding a smoothed phenomenological estimate of muscle force. In ED modelling, the robust process of neuron-fibre synaptic transmission that always elicits post-synaptic action potentials [267]-[269] can be neglected.

A recent eligible study [29] described the muscle’s ED and MURD with experimental measurements of the motoneurons’ discharge activity (discharge and recruitment dynamics); the motoneuron spike trains, identified from the decomposition of HD-EMG signals [270] [271], drive the recruitment and firing behaviour of a population of Hill-type models of motor units that collectively produce the whole muscle force in a motoneuron-driven approach with motor unit resolution. Importantly, in such data-driven approach, the limited sample of motoneuron spike trains identified experimentally is usually not representative of the discharge activity of the complete motoneuron pool and usually does not accurately estimate the neural drive to muscle, as previously demonstrated [256]. To address this issue, two solutions are currently proposed. A first experimental solution relies on using dense grids of EMG electrodes covering the whole muscle to identify larger and more representative populations of motor units than usual from experiments [272], [273]. A second computational solution infers, from limited experimental measurements, the discharge behaviour of the complete motoneuron pool [55].

#### 2) Titin-induced time-history-dependent mechanisms

The classic properties of the CE (e.g., FL, FV) are steady-state and instantaneous and should be extended to describe the time-history dependent transient properties of muscle. This was achieved in a few eligible models (Fig. 5(b)), to avoid potential high prediction errors [95], describe certain behaviours like balance control against perturbations [227], [274], or solve for unphysiological modelling instabilities as discussed in Section IV.D.2). While comprehensive models of muscle fatigue exist [194] and can be readily included in Hill-type models, only phenomenological models of fatigue were considered to date in the single eligible model [88] and in the additional non-eligible Hill-type models [93], [258] reported in SM7 and SM8. Likewise, FD and RFE were modelled phenomenologically in most of the eligible [131] and additional Hill-type models [47], [232], [275]-[277] (reported in SM7 and SM8). As proposed in two Hill-type studies [35], [278], which comprehensively related FD and/or (R)FE to the complex passive and activation-dependent dynamics of titin, RD and RFE are explained by the various dynamics of titin [279], [280], rather than the nonuniformity of sarcomere lengths [171]. Titin can shorten with calcium-mediated binding [171], [281] or winding [282] motions and increases its stiffness upon activation [283]-[286], which also stabilizes the aforementioned descending limb of the instantaneous FL relationship [287]. With an unfolding-folding behaviour at activation [288], titin can also provide power strokes to the force-generating cross-bridges and accelerate muscle shortening [289], [290]. Consistent with the previously discussed ADL properties, titin promotes cross-bridge-attachment by passively pre-stretching non-active sarcomeres [291], [292]. Also included in Huxley-type [293] and continuum [14] models, the titin activity could be phenomenologically modelled in a Hill-type model as a hybrid CE-PEE property that alternates between active [294] and passive [176], [288] states, and is in-series or in-parallel with the cross-bridges upon calcium activation [229].

#### 3) Inertial effects

Although considering muscle mass in Hill-type models typically increases the numerical cost of the model by involving 2^nd^-order ODEs, it can increase numerical stability [31] and reduce the muscle maximum shortening velocity and mechanical work [44], [221]. A mass element is necessary to investigate the contraction-related effects of muscle mass flow [295], on which depends the MSK centre of mass location, and wobbling masses [296], [297] on the reduction of impact forces [298], [299], joint contact forces and torques [296], [300], or bone internal stresses [68].

#### 4) Choice of parameter values

The behaviour of the NE, CE and passive elements is highly dependent on the parameter values that describe the properties displayed in Fig. 2 and discussed previously. These parameter values, which are typically obtained from fitting experimental data with parametrized mathematical relationships, vary with multiscale considerations (see Section IV.D.1)), and are mostly also species-, muscle-, temperature-, and subject-specific. For example, the *a*_*f*_ parameter that describes the curvature of the FVC relationship increases from fibre-to muscle-scale (see Section IV.D.1)), and between slow-type and fast-type fibres [301]-[303]. The parameter that defines the half-time of relaxation of the calcium transients (ADC) drastically decreases with increasing temperatures [238], between slow and fast fibres [304] and between mammalian [237] and amphibian [119] species. Potentially because of limitations in knowledge transfer in the field of Hill-type modelling, discussed later in this section, it was observed that most eligible studies overlooked the importance of these numerical values (Fig. 6(b), Q9, 1.1/2), which were not all provided in half of the publications (Fig. 6(b), Q4). Half of the eligible studies proposed CE and passive properties that were described with parameters experimentally obtained from different species and typically copied from milestone but dated Hill-type modelling studies [56], [119], [123]. It was observed that more recent experimental data are usually not exploited; for example, dated frog data [179] is commonly used over more recent human data [4] to describe human muscles.

According to sensitivity studies listed in SM7 [111], [305]-[310], the predictions of muscle forces in MSK simulations are most sensitive to three parameters that scale the generic (normalized) Hill-type models: the tendon slack length, the optimal length of the CE and the maximum isometric force of the FG. It is worth noting that these observations were made mostly with ‘baseline’ models (Fig. 1), and additional sensitivity studies on ‘more complete’ models are required to generalize these conclusions. While the parameters describing the normalized muscle properties can remain generic, these three scaling parameters should be tuned to be subject-muscle-specific [23]. As referenced in SM12, some eligible studies [24], [26], [61], [124], [311] proposed calibration methods, although various configurations of parameter values can explain similar muscle dynamics [308], depending on the role of the muscle in the MSK task [111], [308], [312]. Other eligible studies experimentally measured these properties in animal [60], [136] or human [3] ad hoc experiments, or used available experimental datasets [116], [146]. Some also scaled these scaling parameters using subject-specific anthropometric measurements and generic literature data [3], [31], [139], [143]. In this approach, recent datasets obtained from large populations of young adults and recent experimental techniques should be preferred for estimating the FG’s optimal length [313] and maximum isometric force [314], as performed previously [315], to smaller dated datasets [316]-[318] obtained from specimens of advanced age that present important age-related muscle atrophy and smaller PCSAs [49], [50], [319], [320]. Finally, close to half of the eligible studies used generic (i.e., not subject-muscle-specific) parameters to scale their generic model.

### F. Principles of model validation

The performance of a Hill-type model can be evaluated by comparing its predictions against experimental quantities obtained in situ or from complete experimental datasets made available in the literature [199], [321], [322]. The gold standard quantity for model validation is the time-course of individual fibre or muscle forces, which can be obtained or derived from dissected muscles of anesthetized animals as in some of the eligible studies (see SM12 for details), calculated from non-invasive ultrasound measurements of fascicle or tendon length in humans [1]-[3], [323], [324], or from specific protocols that minimize muscle co-activation during joint torque measurements [311]. Since the latter methods are relatively complex, most models are validated by comparing model-predicted joint torques against those computed from experimental data (also often used for model calibration for the same activity), joint angles, and neural EMG-like signals, as discussed in the Results Section and detailed in SM12. This approach is characterized by limitations in validation accuracy as such simulated and experimental quantities accumulate errors independent from muscle contraction mechanisms. In all cases, different experimental datasets must be used for model calibration and validation (i.e., different training and test sets). Finally, the validation of model performance should be quantified with objective mathematical metrics (see SM12) to prevent qualitative assessments and bias and improve readability. These important considerations were overlooked by many eligible studies (Fig. 6(b), Q3, Fig. 7).

Once validated, a model may be implemented in other studies providing its performance is objectively considered in light of the modelling choices taken and the physiological plausibility of the calibrated parameter values. Most eligible studies overlooked such practices, as reported by the modelling assessment (Fig. 6(b) Q6 to Q8). Model reusability is also promoted when published results can be reproduced, which is possible when the model can be fully reimplemented, either providing an open-source implementation of the model, or from a detailed description of the modelled properties, rheological arrangement of the elements, parameter numerical values and methods for solving the muscle dynamics, most of which the eligible studies failed to provide (SM12 and Fig. 6(b), Q4 and 5)

### G. Limitations of the study

This systematic review, despite following the PRISMA recommendations (see commented checklist in SM2), comes with a few limitations related to the custom design of the eligibility criteria and scoring schemes.

By constraining the eligible models to the six EC in Table 1, milestone studies specialized in research areas interfacing with Hill-type modelling were overlooked, including for example advances in parameter estimation [325], [326], volumetric muscle definitions [327], [328], the efficient implementation of novel optimization algorithms [329], [330], MSK modelling [37], [315] and in-vivo neural inputs for motor control [28], [331], [332]. Besides disregarding other types of muscle models (EC3), such as Huxley-type or hybrid Hill-Huxley-type [11], [13], [261] and continuum [15]-[17] approaches, some key ‘innovative’ Hill-type models that met EC6 but did not include a NE (EC4) were also not considered for global scoring in this study. To complement the extensive results presented in this study for the 57 eligible studies, those 33 additional studies are reported in SM7 and assessed for completeness in SM8 by a single scorer (A.C.). Besides, although it provides a window on the state-of-the-art in Hill-type modelling, the application of EC6 biases the results towards concluding that the field of Hill-type modelling is populated with rather more complete Hill-type models (Fig. 4, mean: 8.1/23) than the majority of models proposed in the literature, which are actually closer to baseline models (Fig. 1) and would score 5/23 or less.

Besides the 23 properties considered in this study, some important neuromuscular properties, such as the titin dynamics, were not considered for completeness assessment, as they were only recently included in Hill-type models [35]. Also, it is important to reiterate that the completeness assessment provides information about the complexity of a model and its capacity to describe specific muscle mechanisms but does not provide any indication on the quality of the modelling approach. Likewise, the outcomes of the modelling assessment do not provide any indication on the accuracy of the eligible model predictions. The choice of model complexity and number of parameters, a key topic in Hill-type modelling [19], [333], could have been considered in this scoring scheme. Although care was taken to achieve a balanced scoring approach, as discussed in the Methods Section for both the scoring schemes, some modelling and methodological approaches may be under-represented, like the choice of rheological structure or the use of the multiscale simplifications that only account for 2.5% of the global score each (Fig. 8). Finally, if the 57 eligible models are representative enough of the field of Hill-type modelling to describe its state-of-the-art, they may not be representative of the methodological approaches globally taken in the literature about model validation, reusability, modelling choices and strategy and calibration (Table 3). This reduced window may weaken the impact of some conclusions drawn from the Modelling Assessment in this Discussion Section, considering for example that some non-eligible studies would have reached high modelling scores [50], [334].

### H. Required recommendations in the field of Hill-type modelling

As shown in the chronological graph in Fig. 5(c), the state-of-the-art in Hill-type modelling seems to have plateaued since the 80s-90s when substantial modelling innovations occurred. It is notably observed that following the turn of the century modelling advances have focused on describing discrete muscle properties and never build upon other contemporary innovations, as they keep relying on the 50-year-old gold standards (Section IV.C.), rather than challenging their important limitations (Sections IV.D. and IV.E.). This tendency is typical of a dense field, for which the most recent narrative reviews (listed in SM7) date back 20 years or more, and where the transmission of knowledge is therefore limited. To permit innovation and incremental breakthroughs, Epstein and Herzog [98] stress the importance of regular literature reviews to periodically clarify and summarize the field, update on its state-of-the-art and avoid the emergence of detrimental inclinations that further aggravate the situation. One of them is the increasing inconsistency that was observed between the eligible studies in the terminology and notations that refer to, for example, the multiscale FG length as optimal length, normalized length, maximum isometric force, tendon slack length, maximum force in eccentric contractions, but also in the definition of isometric contraction as constant muscle or muscle-tendon lengths. Another detrimental tendency is the chain of references between Hill-type studies that is breaking over time; in such case, inheritance diagrams (SM10) are necessary to trace the current modelling techniques back to their source studies. This unclear literature also pushes researchers towards reusing and not adapting the milestone studies from the 80s-90s with more recent available experimental data (Section IV.E.). In this context, the lack of systematic good practices that is stressed by the modelling assessment (Fig. 6(b)) further hinders the transmission of knowledge and the reusability of published models. It is indeed difficult to build upon a model that is not carefully validated and described, and that is not easily reproducible or for which no open-source implementation is available. In these conditions, researchers seem rather to opt for open-source ready-to-use models [33], [49], [61], [329], [335], as 25% of the studies obtained from the systematic search and respecting EC1 to EC3 since 2006 used the muscle models embedded in the simulation platform OpenSim [37], [38] or the open-source CEINMS package [36]. While these models motivate advances in fields related to muscle modelling, the studies using these models however do not contribute to advancing the Hill-type modelling field itself.

Besides regular reviews that clarify the field and summarize its state-of-the-art, the field of Hill-type modelling would strongly benefit from recommendations developed by the research community, as proposed in other related fields, e.g. the recommendations of the International Society of Biomechanics regarding reference system definitions [336], [337], reporting of joint kinematics [338] and intersegmental loading calculation [339], etc. These recommendations could propose (1) standard definitions, notations and terminology for the common variables and dynamics of Hill-type models with respect to the chosen multiscale approach and rheological structure. As done for MSK systems [340], those recommendations could provide a checklist and advise on (2) some best methodological practices, including the choice of mathematical metrics and experimental data for model validation, stating the assumptions and limitations related to the multiscale and rheological approaches, choosing the model complexity and completeness with regards to the study aim, taking the best approach for scaling and calibrating the model, and discussing its strengths and weaknesses in light of other findings in the literature. These recommendations should (3) refer to the sensitivity analyses in the field that have identified the most important model parameters, for which datasets compiling published experimental data for different species, muscles and scales should be produced for an optimized scaling of the models. Finally, they should provide (4a) a checklist of the details to include in the main manuscript or its supplementary material for the full re-implementation of the model including the rheological structure, the equations describing the modelled properties, and a table listing the parameter values describing these properties and scaling the model, and (4b) encourage modellers to release open-source implementations of their models so that the results presented in their study can be reproduced.

## V. Conclusion

This study is the first systematic review of the field of Hill-type muscle-tendon modelling. The results of the completeness assessment (Fig. 4 and Fig. 5) describe the current trends and state-of-the-art in phenomenologically modelling 23 muscle properties in Hill-type systems. The results of the modelling assessment (Fig. 6 and Fig. 7) evaluate the current application of good practices in the field about model validation, reusability, modelling strategy and calibration. Supported by extensive supplementary material, global scores (Fig. 8), and updated terminology (see Discussion Section and Table 2), this work is a convenient tool for scientists and modellers who wish to better understand the literature, choose the most suitable available model for their muscle-related study, or advance the state-of-the-art in Hill-type modelling. This study stresses the importance of contemporary reviews and global modelling recommendations to optimize inter-study consistency, knowledge transfer, and model reusability.

## Supporting information

Scoring Sheet for both assessments

