## Supplementary Material (SM) for "Hill-type models of skeletal muscle and neuromuscular actuators: a systematic review"

### SM 1: Concise historical excursus of Hill-type modelling

The definition of a Hill-type muscle model has emerged from preliminary studies published in the first half of the 20th century. While skeletal muscles were described as rheological arrangements of purely elastic and viscoelastic elements in the late 19th century, Hill proposed in 1922 (Hill, 1922) a new metabolic energy-based force generating element, later called contractile element (CE), that Levin et al. further connected in 1927 (Levin & Wyman, 1927) to an undamped series elastic element. Hill proposed in 1938 (Hill, 1938) in a milestone study the bases of Hill-type modelling with the derivation of the Force-Velocity (FV) hyperbolic relationship (Hill's equation), further developed by others (Katz, 1939), a definition of the active state, and a description of the muscle with three elements including an active CE and passive series and parallel elements in a Maxwell rheological configuration (Maxwell, 1867). The pioneer works from Wilkie, Ritchie, Abbott, Jewell, and other collaborators from their research groups then extended Hill's work in 1950-1960 and derived most of the current gold standards in modelling methods. They validated and calibrated Hill's FV hyperbolic relationship for several muscles, species and fibre types (Wilkie, 1949; Jewell & Wilkie, 1958; Wells, 1965), compared the physiological accuracy of the Maxwell and Kelvin-Voigt rheological structures including a CE (Jewell & Wilkie, 1958), provided a first physiological definition of the active state (Wilkie, 1956), derived a Force-Length (FL) relationship linearly scaled by the active state (Wilkie, 1956; Ritchie & Wilkie, 1958), made Hill's hyperbolic curve active state and length nonlinearly dependent (Abbott & Wilkie, 1953; Wilkie, 1956), and inverted the FV relationship to unveil a first-order differential equation in muscle length relating variations in muscle length with time (Wilkie, 1956; Ritchie & Wilkie, 1958). After applications in cardiac muscle modelling in 1960-1970 (Fry et al., 1964; Sonnenblick, 1965; Brady, 1967), the Hill-type modelling approach was simultaneously extended by the emerging sliding filament theory (Huxley, 1957) and cross-bridge cycle mechanism (Huxley & Hanson, 1954). The current cross-bridge-based definition of the active state was proposed in 1968 (Ebashi & Endo, 1968), while Gordon et al. reconciled in 1966 (Gordon et al., 1966) the phenomenological FL relationship to the degree of sarcomere filament overlap, enabling sarcomere-to-muscle multiscale modelling approaches. Hill-based models gradually became in 1960-1970 a class of muscle models with emerging terminology such as 'Hill's three-element / component model' (Buchthal & Rosenfalck, 1960; Bahler et al., 1967), and 'Hill-type model' (Grood et al., 1974) and the first computational Hill-type model relying on solving time-dependent differential equations of muscle lengths to predict muscle quantities (Bahler, 1968). In 1970-1980 were proposed the first accurate mathematical descriptions of the FV relationship in eccentric contractions (Mashima et al., 1972; FitzHugh, 1977) that the authors are aware of. Subsequently, Hatze derived and validated in 1976 the force of the contractile element as obtained from the linear product of the active state and the instantaneous forces as obtained from the FL and FV relationships, and developed the first Hill-type muscle driven musculoskeletal simulation additionally using processed EMGs as inputs (Hatze, 1976). In 1990-2000, four contributions to the literature assessed the state-of-the-art of Hill-type MT modelling to propose (1) a robust theoretical scheme encouraging Hill-type modelling with normalized properties that can be used to represent, once scaled with architectural parameters, any specific muscle (Zajac, 1989), (2) two didactic introductions to Hill-type modelling (Epstein, 1998; Yamaguchi, 2001), and (3) a detailed review comparing the numerous modelling approaches (Winters, 1990) that had increasingly emerged since the 1980s.

### SM 2: Methods - PRISMA checklist (Moher et al., 2009)

| PRISMA item n° | PRISMA Checklist items | Respected [Y/N] | Comments if relevant |
|---|---|---|---|
| 1 | Identify the report as a systematic review, meta-analysis, or both | Y | |
| 2 | Provide a structured summary including, as applicable: background; objectives; data sources; study eligibility criteria, participants, and interventions; study appraisal and synthesis methods; results; limitations; conclusions and implications of key findings; systematic review registration number. | Y | |
| 3 | Describe the rationale for the review in the context of what is already known | Y | |
| 4 | Provide an explicit statement of questions being addressed with reference to participants, interventions, comparisons, outcomes, and study design (PICOS). | Y | Not completely applicable as defined as not a clinical systematic review, though 3 distinct objectives for the systematic review are defined |
| 5 | Indicate if a review protocol exists, if and where it can be accessed (e.g., Web address), and, if available, provide registration information including registration number. | Y | This is the first systematic review on Hill-type modelling though systematic reviews on muscle modelling-related fields are referred to in the text. No review protocols exist. |
| 6 | Specify study characteristics (e.g., PICOS, length of follow-up) and report characteristics (e.g., years considered, language, publication status) used as criteria for eligibility, giving rationale. | Y | Eligibility criteria: Journal article in English for the 1938-2024 period, specified in the Methods section |
| 7 | Describe all information sources (e.g., databases with dates of coverage, contact with study authors to identify additional studies) in the search and date last searched. | Y | |
| 8 | Present full electronic search strategy for at least one database, including any limits used, such that it could be repeated. | Y | Most details are provided in SM3 along with the key term search string for the PubMed database |
| 9 | State the process for selecting studies (i.e., screening, eligibility, included in systematic review, and, if applicable, included in the meta-analysis) | Y | |
| 10 | Describe method of data extraction from reports (e.g., piloted forms, independently, in duplicate) and any processes for obtaining and confirming data from investigators | N | Not applicable as not a clinical systematic review relying on data |
| 11 | List and define all variables for which data were sought (e.g., PICOS, funding sources) and any assumptions and simplifications made. | Y | 23 properties and 10 questions around methodological practices are defined for which data is sought |
| 12 | Describe methods used for assessing risk of bias of individual studies (including specification of whether this was done at the study or outcome level), and how this information is to be used in any data synthesis. | Y | The study inclusion and the scoring were performed by two independent reviewers. A consensus was reached after discussion in case of different conclusions |

| | | | |
|---|---|---|---|
| 13 | State the principal summary measures (e.g., risk ratio, difference in means) | Y | The review is mainly focused on trends, for which simple ratio measurements are suitable objective criteria |
| 14 | Describe the methods of handling data and combining results of studies, if done, including measures of consistency (e.g., I2 ) for each meta-analysis | N | Not applicable |
| 15 | Specify any assessment of risk of bias that may affect the cumulative evidence (e.g., publication bias, selective reporting within studies) | Y | This is discussed in the limitations of the study in the Discussion section. The chosen eligibility criteria 4 to 6 and the choice of scoring schemes bring mild bias to the conclusions |
| 16 | Describe methods of additional analyses (e.g., sensitivity or subgroup analyses, meta-regression), if done, indicating which were pre-specified. | N | Not applicable |
| 17 | Give numbers of studies screened, assessed for eligibility, and included in the review, with reasons for exclusions at each stage, ideally with a flow diagram. | Y | The tree diagram of study selection with step-by-step details is provided in SM5 |
| 18 | For each study, present characteristics for which data were extracted (e.g., study size, PICOS, follow-up period) and provide the citations. | N | Not applicable |
| 19 | Present data on risk of bias of each study and, if available, any outcome-level assessment (see Item 12). | N | Not applicable |
| 20 | For all outcomes considered (benefits or harms), present, for each study: (a) simple summary data for each intervention group and (b) effect estimates and confidence intervals, ideally with a forest plot | Y/N | The resulting marks for completeness and modelling assessments and global scores were provided for each study |
| 21 | Present results of each meta-analysis done, including confidence intervals and measures of consistency | N | Not applicable |
| 22 | Present results of any assessment of risk of bias across studies (see Item 15). | N | Not applicable |
| 23 | Give results of additional analyses, if done (e.g., sensitivity or subgroup analyses, meta-regression). | Y | Trends in Hill-type modelling approaches, their time-evolution and trends in methodological practices are unveiled |
| 24 | Summarize the main findings including the strength of evidence for each main outcome; consider their relevance to key groups (e.g., health care providers, users, and policy makers). | Y | |
| 25 | Discuss limitations at study and outcome level (e.g., risk of bias), and at review level (e.g., incomplete retrieval of identified research, reporting bias). | Y | |
| 26 | Provide a general interpretation of the results in the context of other evidence, and implications for future research. | Y | |
| 27 | Describe sources of funding for the systematic review and other support (e.g., supply of data); role of funders for the systematic review. | Y | |

### SM 3: Methods – Key terms for the systematic search

The keyterms are grouped into 4 concepts. Within each concept, the keyterms reported in Table 1 are joined with the OR Boolean operator. The systematic search is performed with the string Concept 1 AND Concept 2 AND Concept 3 NOT Concept 4.

PubMed was performed with the following string:

(((hill[tiab] or musculotendon[tiab] or myocybernetic[tiab]) **AND**

(muscl*[tiab] or muscu*[tiab] or tendon[tiab] or myocybernetic[tiab])) **AND**

(model*[tiab] or actuat*[tiab] or system*[tiab] or "contractile element*"[tiab] or complex*[tiab] or unit*[tiab] or simulat*[tiab] or dynamics*[tiab] or optimization[tiab] or "EMG-driven"[tiab] or "Muscle-driven"[tiab] or control*[tiab] or predict*[tiab] or estimat*[tiab])) **NOT**

(smooth[tiab] or cardiac*[tiab] or myocard*[tiab] or papillary[tiab] or heart*[tiab] or myocyt*[tiab] or virus*[tiab])

**Filter:** English, Journal article

And updates were enabled with a weekly frequency to update the review with the most recent published studies.

*Table 1: Keyterms used for the systematic seach, grouped into concepts and free-text and mesh terms*

| | **Concept 1: Hill-type musculotendon models** | **Concept 2: the skeletal muscle and the tendon** | **Concept 3: model – actuator - simulation** | **Concept 4: excluding cardiac and smooth muscles** |
|---|---|---|---|---|
| **Free-text terms** | Hill | Muscl*, muscu* | Model* | Cardiac*<br>Myocard*<br>Papillary<br>Heart*<br>Myocyt* |
| | Musculotendon | Tendon | Actuat*,<br>System*,<br>Contractile element*,<br>complex*,<br>unit* | Smooth |
| | Myocybernetic | myocybernetic | Simulat*<br>Dynamics<br>Optimization<br>EMG-driven,<br>Muscle-driven<br>Control<br>Predict*, estimate* | |
| **Mesh Terms** | | Muscle, Skeletal / physiology | | |

#### SM 4: Methods - Details of the scoring scheme of the Modelling Assessment

In order to yield an objective scoring and a robust comparison between eligible studies for the two reviewers, a detailed scoring scheme was developed and is reported in Table 2. Each paper would score 0, 1, or 2 to the 10 questions following precise criteria. These criteria were derived before the systematic search according to a preview of the literature of a hundred Hill-type studies chosen randomly in the literature.

The 10 questions are provided in the core text of the review. As detailed in the main text of the study:

- (I): Model validation
- (II): Model reusability
- (III): Modelling choices and strategy
- (IV): Model calibration.

*Table 2: Details on the scoring scheme of the Modelling Assessment*

| Benchmarks | Question number | Scoring | Scoring scheme |
|---|---|---|---|
| (I) | Q1 - Is the accuracy of the predictions obtained from the Hill-type model validated against ad hoc experimental data? | 0 | • No validation of the accuracy of the model's predictions, or<br>• Comparison of the model's predictions with user-defined target quantity, or<br>• Comparison of the model's predictions with results from other models, or<br>• Comparison of the model's predictions with inconsistent experimental results from the literature (different muscle, species, protocol of contraction) |
| | | 1 | Predicted results are compared against a test set of experimental data from the literature, the experimental conditions of which are carefully reproduced (e.g., same species and muscles, same stimulation and length-varying protocol) |
| | | 2 | The accuracy of the model's predictions is blindly validated on<br>• a test set of ad hoc experimental data, or<br>• quantities derived from ad hoc experimental data, or<br>• experimental data collected by the same group of authors and reported in another study |
| | Q2 - Are the model's predictions validated against reliable muscle- | 0 | Question 1 scored 0 |
| | | 1 | The model is validated by comparing quantities predicted by the muscle model and quantities that provide an indirect description of muscle activity and/or cannot be precisely measured experimentally:<br>• Joint torques measured by dynamometers and/or with load cells, or<br>• Limb motion (e.g., joint angles), or<br>• Muscle activation profiles (e.g., EMG envelopes), or<br>• Muscle energy consumption |

| | | | |
|---|---|---|---|
| | specific experimental quantities? | 2 | The validation of the model is performed on the actuator's force or length profiles (valid at any scale, e.g., fibre, fibre bundles, muscle, tendon, or muscle-tendon complex) |
| | Q3 - Is the accuracy of the model evaluated using objective, quantitative metrics? | 0 | • No model validation was proposed and/or<br>• the evaluation of the model's accuracy is absent or qualitative (e.g., no discussion on model validation, qualitative comparison of predicted and experimental curves, qualitative assessment of the results) |
| | | 1 | The evaluation of the model's accuracy is quantitative but performed using basic descriptive objective criteria (e.g., percentages, ratios, ranges… when comparing predicted and experimental results) |
| | | 2 | The evaluation of the results of the model validation is addressed with<br>• advanced quantitative objective criteria (e.g., RMS error), or<br>• statistical analysis criteria when applicable (e.g., correlation coefficient) |
| (II) | Q4 - Is enough information provided to reimplement the neuromuscular model? | 0 | The model cannot be fully re-implemented:<br>• Some mathematical expressions describing the material properties (e.g., the differential equation describing the activation dynamics) cannot be retrieved, and/or<br>• Key parameter values are not provided explicitly or by references (e.g., maximum isometric force, or the parameters values of the stress-strain relationship of the tendon) |
| | | 1 | The model is almost fully re-implementable and requires personal effort to be reproduced:<br>• All the mathematical expressions that describe the material properties are reproduceable (explicitly provided or careully referenced), and<br>• Most of the parameter values are provided, and<br>• Yet, any of:<br>o Some mathematical equations describing the material properties are only provided through references, and/or<br>o Some parameter values and/or mathematical relationships between material properties are not provided |
| | | 2 | The model is easily and fully re-implementable<br>• The implementation is provided in supplementary material, or<br>• The code implementing the model is provided as supplementary resource or through open-source repositories, or<br>• All mathematical expressions and parameter values (except potential rare parameter values of lesser importance) defining the material properties are explicitly provided, and<br>• The rheological structure and / or the mathematical relationships between material properties are provided, and<br>• There is help provided in re-implementing the model and / or in reproducing the results of the study |
| | Q5 - Is the model optimized for numerical stability and | 0 | No |
| | | 1 | Only remarks are done on the computational efficiency of Hill-type model in general |
| | | 2 | The model includes at least one voluntary alteration of the Hill-type model that improves the model stability, computational speed, or avoids numerical stiffness, singularities or unnecessary model complexity (e.g., non-zero values to avoid singularities in the force-velocity relationship) |

| | | | |
|---|---|---|---|
| | computational speed? | | |
| | Q6- Are the simplifications adopted in modelling the muscle, both physiological and across relevant scales, clearly stated? | 0 | The taken simplifications with regards to muscle physiology, geometry, and/or multiscale architecture are not explicitly stated. |
| | | 1 | At least one taken modelling simplification is stated with regards to<br>• muscle physiology (e.g., muscle material is assumed massless, frictionless, incompressible, or isotropic) and/or<br>• muscle external geometry (e.g., inclusion/exclusion of the pennation angle, muscle is simplified as a rectilinear pathway), and/or<br>• muscle internal architecture (all fibres are coplanar, parallel straight, or of same length). |
| | | 2 | • Explicit comments about the simplification across physiological scales (sarcomere, fibre, fibril, etc) approach taken in Hill-type modelling (e.g., the muscle is modelled as a scaled representative constitutive fibre, or its neural control is considered functionally equivalent to a single motor unit), and/or<br>• explicit assumption of homogeneous and/or averaged material properties and/or similar dynamics between contractile sub-scale elements |
| (III) | Q7 - Are the modelling choices for modelling the rheological model clearly stated and / or justified with respect to the scope and objectives of the study? | 0 | Incomplete and unjustified modelling choices of the rheological model and its material properties |
| | | 1 | • The arrangement of the rheological elements in the mechanical structure is explicitly stated (e.g., the CE is mathematically modelled in-series with the tendon), and<br>• Each rheological element and its constitutive mathematical expression (e.g., a tendon and its constitutive stress-strain relationship) are explicitly described, and<br>• Yet, no commented reason is given for these modelling choices (e.g., why a typical rheological element of the baseline model (e.g., tendon) was overlooked, or why a more advanced property (e.g., calcium-dependent activation dynamics) was included in the model, or the experimental or physiological reasons justifying the proposed mathematical expressions) |
|  |  | 2 | • The arrangement of the rheological elements in the mechanical structure is explicitly stated, and<br>• Each rheological element and its constitutive mathematical expression are explicitly described, and<br> ○ The modelling choices are justified with references, and/or considering the aim and scope of the study, or<br> ○ Elements and properties are critically commented and linked back to physiological mechanisms, or |
| | Q8 - Is there a discussion about the strengths and limitations of the normalized Hill-type model? | 0 | No concluding remarks on the strengths and limitations of the proposed Hill-type model and its predictions with regards to the scope of the study |
| | | 1 | There are concluding remarks on the strengths and limitations of the model, and<br>• Hill-type modelling is not the main topic of the study, and / or<br>• The analysis is only qualitative and not supported by references and findings from other studies |
| | | 2 | • There are concluding remarks on the strengths and limitations of the model, supported by a quantitative analysis (e.g., sensitivity analysis of modelling choices and/or parameters on metric, or quantitative analysis of the latent predicted state variables of the model against literature data), and / or<br>• Qualitative analysis supported by references and findings from other studies (e.g., commenting and referencing on the weakness of Hill-type models for accurate predictions in submaximal contractions) |

| | | | |
|---|---|---|---|
| (IV) | Q9 - Are the mathematical expressions describing the properties of the normalized Hill-type model parametrized from experimental data consistent with the scope of the study? | 0 | • The mathematical expressions defining the normalized Hill-type model (e.g., tendon stress-strain relationship, muscle force-velocity relationship) are parametrized with arbitrary parameter values, or<br>• Parameter values imported from a previous Hill-type model without critical insight on their origin, or<br>• Parameter values taken from non-consistent literature (e.g., parameters values are taken from different species or muscles, and/or from species and muscles inconsistent with the scope of the study) |
| | | 1 | There is significant effort made for parametrizing the mathematical expression of the normalized Hill-type model with parameters obtained from:<br>• The literature for the same species, muscle-tendon complex and scale (e.g., from experimental studies, or studies that propose a Hill-type model built from ad hoc experiments), and / or<br>• A consensus drawn from a significant review of the literature |
| | | 2 | At least one parameter used in the parametrization of the mathematical expressions of the normalized Hill-type model was obtained from:<br>• The literature for the same species and muscle, and was further scaled for the subject, or<br>• A complete experimental database from a single study for the same species, muscle, and scale (providing input conditions, output force profiles, and experimentally derived parameters), or<br>• An ad hoc sensitivity study, or<br>• Ad hoc muscle-specific experiments, or<br>• Parameter calibration by minimizing an objective function related to model predictions |
| | Q10- Is the normalised Hill-type model defined with species- patient- or muscle-specific architectural scaling parameters? | 0 | The normalised expressions describing the properties of the Hill-type muscle model are not scaled, i.e. denormalised using architectural and contraction parameters to represent actual muscle(s) (e.g., maximum isometric force, optimal fibre length, tendon slack length, maximum shortening velocity, ratios of fibre types, motor unit pool size), or scaled with arbitrary architectural parameter values |
| | | 1 | • The normalized Hill-type model is scaled with at least one architectural scaling parameter to represent actual muscle(s), and<br>• the parameter is obtained from the literature and is at least species-consistent and/or muscle-consistent (e.g., scaling generic Hill-type models of human muscles with muscle-specific maximum isometric force values that are extracted from a databank of cadaveric human measurements available in the literature) |
| | | 2 | The normalized Hill-type model is scaled with at least one architectural scaling parameter, and the parameter value is either obtained from<br>• A muscle-specific or subject-specific scaling of a parameter obtained from the literature and that scored 1 (e.g., a species- and muscle-specific reference value of tendon slack length is linearly scaled according to anthropometric measurements to make it subject-specific), or<br>• Parameter calibration (e.g., calibration of optimal fibre length by minimizing the difference between predicted and measured joint torques), or<br>• Ad hoc experiments (e.g., segmented medical images for the derivation of muscle-specific values of maximum isometric force) |

#### SM 5: Results - Flowchart of paper selection

This flowchart is a detailed version of Fig. 3 of the main manuscript. In Figure 1 are detailed the steps taken to identify the eligible studies, according to the eligibility criteria defined in Table 1 of the main manuscript.

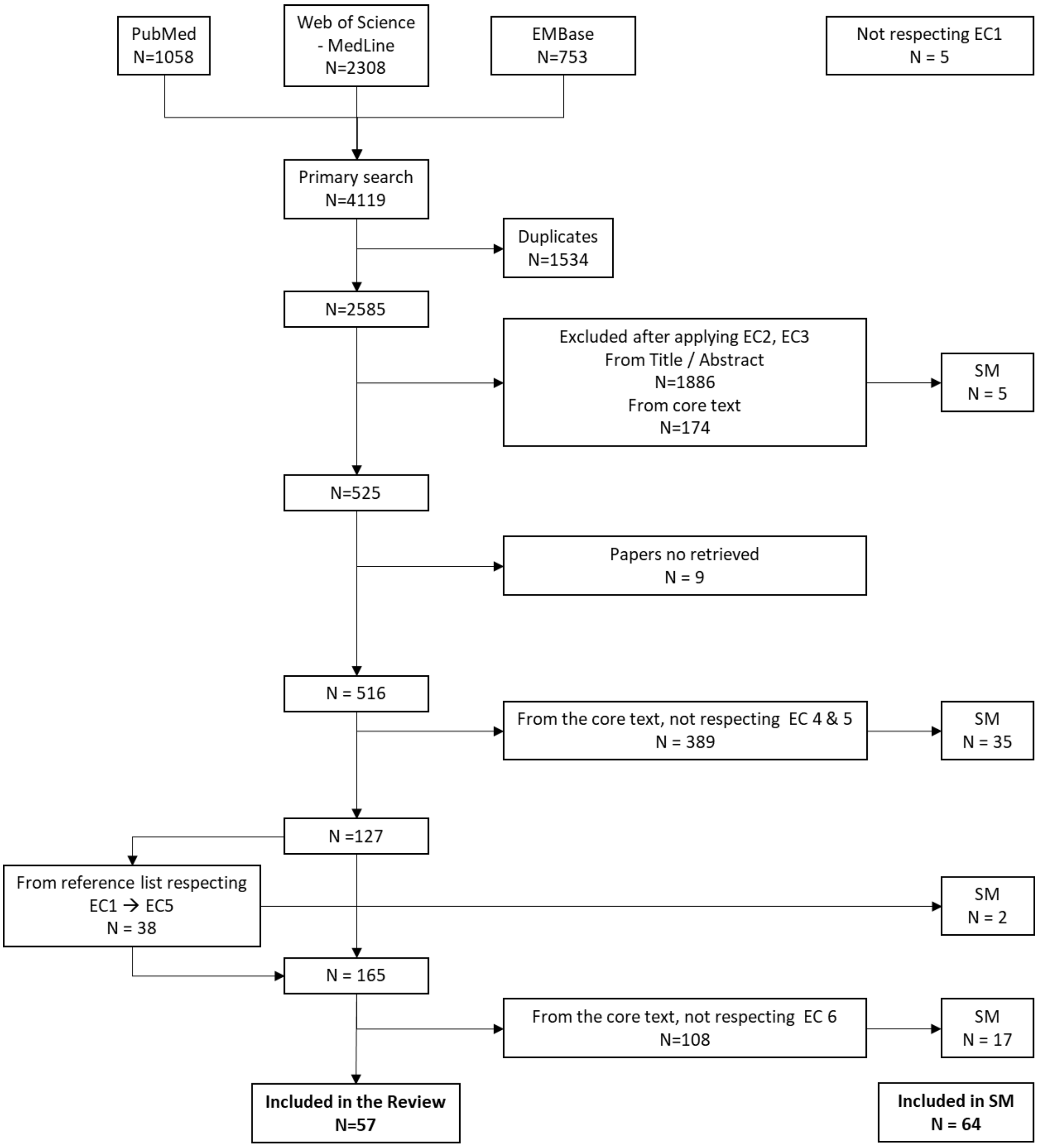

*Figure 1: Flowchart of the selection of the eligible papers after application of EC1 to EC6. N corresponds to the number of retained or discarded studies when and after the relevant tasks are performed. SM (Supplementary Material) depicts the contributory studies that were retained from the search, 34 of which proposed innovative but non-eligible models for which a parallel assessment for completeness was performed in Section SM 8.*

### SM 6: Results – Reasons why the eligible models meet EC6

SM6 details the reasons why the 57 eligible studies fulfilled EC6. Their modelling contributions to the field of Hill-type modelling are gathered in Table 3. It is recalled that these studies fulfilled EC6 if they were considered to be innovative, i.e., they proposed a modelling approach to describe at least one of the properties of the Hill-type model (see Table 2 of the main manuscript) in a way that had not been proposed yet in any of the previous eligible studies. To limit the number of eligible studies, we disregarded for fulfilment of EC6 any innovation related to (1) the arrangement of the elements (NE, CE, passive elements) of the rheological structure, (2) the method for parameter calibration, (3) optimization of computational runnability, (4) improvements in MSK simulation-related topics and (5) the mathematical description of the FL relationship and PEE. Acronyms describing modelling contributions are taken from Fig. 2 of the manuscript.

*Table 3: Eligible studies and their modelling contribution to the field of Hill-type modelling*

| *#* | *Study* | *Ref (in the main manuscript)* | *Ref (in the SM)* | *Modelling contribution to the field* |
|---|---|---|---|---|
| 1 | Bahler 1968 | [118] | (Bahler, 1968) | First computational Hill-type model<br>First Hill study proposing the widely-used $f_l(\Delta l) = \exp\left(-\left(\frac{\Delta l}{W}\right)^2\right)$ gaussian mathematical expression for the FL relationship |
| 2 | Hatze 1977 | [119] | (Hatze, 1977) | Detailed model of ED<br>Detailed model of ADC and ADL, inspiring many subsequent studies<br>First inclusion of an FVE, PDE and PEE2 in a computational Hill-type model<br>First Hill study proposing the widely-used $0;\ k_1\left(e^{k_2 l^M} - 1\right)$ piecewise mathematical expression for the PEE and the SEE |
| 3 | Hatze 1978 | [120] | (Hatze, 1978) | First phenomenological model of MURD |
| 4 | Wyss 1984 | [121] | (Wyss & Pollak, 1984) | Modelling the ED as processed EMGs<br>Models the AD as a twitch event<br>FVL<br>Using torque angle relationship |
| 5 | Pierrynowski 1985 | [122] | (Pierrynowski & Morrison, 1985) | Custom model of MURD<br>Custom AD<br>Inclusion of a SEE2 |
| 6 | Otten 1987 | [123] | (Otten, 1987) | Modelling the ED as processed EMGs<br>Custom ADC |
| 7 | Bobbert 1990 | [124] | (Bobbert & van Ingen Schenau, 1990) | First eligible paper modelling the AD with a 1st-order ODE, inspired from other innovative but non-eligible studies (Clark & Stark, 1974; Winters & Stark, 1988)<br>First Hill study proposing the widely-used $0; k(l - l_s^T)^2$ piecewise mathematical expression for the SEE |
| 8 | Giat 1993 | [88] | (Giat et al., 1993) | First trigonometric form for a C2 FVC-FVE relationship<br>Mass<br>Fatigue |
| 9 | Van Soest 1993 | [60] | (van Soest & Bobbert, 1993) | Custom FVL, FVA<br>C2 FVE-FVE junction<br>First Hill study proposing the widely-used $1 + c - 2cl^M + cl^{M^2}$ mathematical expression for the FL relationship<br>First Hill study proposing the widely-used $0; k(l - l_s^T)^2$ piecewise mathematical expression for the PEE |
| 10 | Happee 1994 | [125] | (Happee, 1994) | Custom AD |
| 11 | Durfee 1994 | [126] | (Durfee & Palmer, 1994) | Custom model of ED |

| | | | | |
|---|---|---|---|---|
| 12 | Winters 1995 | [56] | (Winters, 1995) | Important source in modelling the AD as active state-dependent (de)activation rate<br>FLA<br>Custom FVL, FVA<br>Activation-dependent PEE2 |
| 13 | Caldwell 1995 | [42] | (Caldwell, 1995) | Custom steady-state AD |
| 14 | Van Zandwijk 1996 | [127] | (Van Zandwijk et al., 1996) | Custom steady-state part of ADC |
| 15 | Dorgan 1997 | [30] | (Dorgan & O'Malley, 1997) | Custom model of MURD |
| 16 | Shue 1998 | [128] | (Nussbaum & Chaffin, 1998) | Custom model of steady-state AD |
| 17 | Nussbaum 1998 | [129] | (Nussbaum & Chaffin, 1998) | Simple model of MURD<br>ADV<br>Short-range stiffness |
| 18 | Curtin 1998 | [130] | (Curtin et al., 1998) | ADC<br>Double-hyperbolic FVC |
| 19 | Brown 1999 | [116] | (Brown et al., 1999) | ADL<br>ADV<br>Sag<br>Yield |
| 20 | Cheng 2000 | [31] | (Cheng et al., 2000) | Custom MURD |
| 21 | Ettema 2000 | [131] | (Ettema & Meijer, 2000) | FD<br>RFE |
| 22 | De Woody 2001 | [132] | (DeWoody et al., 2001) | Considers muscle mass but then eliminates it to reduce system dimension |
| 23 | Gallucci 2002 | [133] | (Gallucci & Challis, 2002) | First Hill study introducing the widely-used $1 - \frac{\Delta l^2}{W^2}$ mathematical expression for the FL relationship |
| 24 | Lloyd 2003 | [26] | (Lloyd & Besier, 2003) | Custom model of ED involving EMG processing<br>Custom AD<br>FLA |
| 25 | Thelen 2003 | [49] | (Thelen, 2003) | Custom FVA |
| 26 | McLean 2003 | [62] | (McLean, Scott G. et al., 2003) | Custom FVA |
| 27 | Stelzer 2006 | [134] | (Stelzer & Von Stryk, 2006) | Custom modelling of the PEE |

| | | | | |
|---|---|---|---|---|
| | | | | Wobbling masses (soft tissue mass dynamically shifting relative to the bone) |
| 28 | Kistemaker 2006 | [135] | (Kistemaker et al., 2006) | Custom FVL, FVA |
| 29 | Günther 2007 | [136] | (Günther et al., 2007) | Custom FVL, FVA<br>First SDE among eligible studies |
| 30 | Mavritsaki 2007 | [137] | (Mavritsaki et al., 2007) | Custom ED, MURD, ADC, AD |
| 31 | Gömmel 2008 | [138] | (Gömmel et al., 2007) | Custom FVL, FVA |
| 32 | Song 2008 | [32] | (Song et al., 2008) | Custom ED and MURD |
| 33 | Menegaldo 2009 | [139] | (Menegaldo & Oliveira, 2009) | Custom FVC |
| 34 | Moody 2009 | [68] | (Moody et al., 2009) | Mass |
| 35 | Rengifo 2010 | [140] | (Rengifo et al., 2010) | Custom FV relationship |
| 36 | Proctor 2010 | [141] | (Proctor & Holmes, 2010) | Comprehensive model of ED |
| 37 | Tsianos 2011 | [142] | (Tsianos et al., 2011) | Custom ED and MURD |
| 38 | Mörl 2012 | [48] | (Mörl et al., 2012) | SDE |
| 39 | Wang 2012 | [143] | (Wang et al., 2012) | Custom modelling of the PEE2 |
| 40 | Blümel 2012 | [144] | (Blümel et al., 2012) | Custom AD<br>Custom FVA |
| 41 | Maceri 2012 | [145] | (Maceri et al., 2012) | Custom model of ED<br>Multiscale tendon model |
| 42 | Callahan 2013 | [146] | (Callahan et al., 2013) | Custom FVL, FVA<br>MURD inspired from Fuglevand 1993 |
| 43 | Millard 2013 | [33] | (Millard et al., 2013) | First introduction of quintic Bezier splines |
| 44 | John 2013 | [61] | (John et al., 2013) | Custom FVA |
| 45 | Lee 2013 | [147] | (Lee et al., 2013) | Custom model of ED and MURD from EMG processing and analysis<br>2 CE-approach for fast and slow fibres |
| 46 | Elias 2014 | [148] | (Elias et al., 2014) | Custom steady-state AD |

| | | | | |
|---|---|---|---|---|
| 47 | Hamouda 2016 | [149] | (Hamouda et al., 2016) | Custom MURD<br>Tackles the limitations of the FL relationship (instability of descending limb) |
| 48 | Dick 2017 | [3] | (Dick et al., 2017) | Custom model of ED and MURD from EMG processing and analysis<br>2 CE-approach for fast and slow fibres |
| 49 | Marcucci 2017 | [150] | (Marcucci et al., 2017) | Custom AD<br>FE approach for analysis of the validity of the multiscale approach |
| 50 | Ross 2018 | [151] | (Ross et al., 2018) | SDE<br>Mass |
| 51 | Kim 2015, 2018 | [34], [117] | (Kim et al., 2015; Kim & Kim, 2018) | Custom comprehensive ED<br>ADC / ADL |
| 52 | Siebert 2018 | [152] | (Siebert et al., 2018) | Transversal PEE |
| 53 | Rockenfeller 2020 | [153] | (Rockenfeller et al., 2020) | Custom ADL |
| 54 | Warner 2020 | [154] | (Warner et al., 2020) | Custom AD |
| 55 | Hussein 2022 | [155] | (Hussein et al., 2022) | Custom ADC |
| 56 | Millard 2023 | [35] | (Millard et al., 2023) | Includes titin dynamics |
| 57 | Caillet 2023 | [29] | (Caillet et al., 2023) | Custom ADC, ADL – Includes experimental and calibrated ED and MURD |

### SM 7: Results – Non-eligible studies with significant contributions to the field

Table 4, Table 5 and Table 6 list the non-eligible studies that were identified with the systematic search and were observed to provide important contributions to the field of Hill-type modelling while not meeting all the eligibility criteria EC1 to EC6. Table 4 focuses on milestone Hill-type models and review studies that shaped the field. Table 5 focuses on the studies that investigated the field of Hill-type modelling, including sensitivity studies, studies comparing the behaviour of various muscle and Hill-type models, or studies about the optimization of the numerical cost of Hill-type models. Table 6 gathers all the non-eligible studies that provide an innovative modelling contribution to the field: these studies usually did not meet EC4 but did meet EC6. As was done in Figure 2 below for the eligible studies, the completeness assessment was also performed for the non-eligible studies from Table 6 in Figure 5. Overall, the eligible studies are more complete than this set of non-eligible studies. As was concluded for the eligible studies, the AD, FL, FVC, FVE, PEE and SEE properties are also more frequently modelled in the non-eligible studies than the remaining 17 other properties (Figure 6).

*Table 4: Milestone and review studies in Hill-type modelling, important sources of inspiration for later studies*

| *#* | *Reference* | *Important features* | *Reason why not eligible* |
|---|---|---|---|
| 1 | (Winters & Stark, 1985) | Important source in modelling the AD with a 1st order ODE | Joint angle state variable (EC5)<br>No explicit AD (EC4) |
| 2 | (Zajac, 1989) | Landmark review on Hill-type modelling | Does not include mathematical descriptions for the CE and passive element properties (EC4) |
| 3 | (Winters, 1990) | First narrative review on the properties of Hill-type models | Does not propose a final Hill-type model (EC3) |
| 4 | (Delp et al., 1990) | Hill-type model implemented in simulation platform (SIMM)<br>Source of standard parameter values for human subjects | PhD thesis, not a Journal article (EC1)<br>No AD (EC4) |
| 5 | (Kaufman et al., 1991) | Review on the theoretical formulation of Hill-type models<br>Work on the index of architecture | Does not propose a final Hill-type model (EC3) |
| 6 | (He et al., 1991) | Important source in modelling the AD with a 1st order ODE | No explicit mathematical description of the CE properties (EC4). They are obtainable in He's PhD thesis which is not publicly available |
| 7 | (Schutte & Margaret, 1993) | Important source of inspiration for subsequent models in the field<br>Implemented in OpenSim | PhD thesis, not a Journal article (EC1) |
| 8 | (Van Leeuwen, 1992) | Narrative review | Book chapter, not a journal article (EC1) |
| 9 | (Epstein, 1998) | Narrative review on theoretical models of Skeletal muscle with a focus on Hill-type models | Book, not a journal article (EC1) |
| 10 | (Yamaguchi, 2001) | Narrative review on modelling muscle and tendon with a Hill-type approach | Book chapter, not a journal article (EC1) |
| 11 | (Heinen et al., 2016) | Narrative review on the scaling methods of Hill-type musculoskeletal models | Does not build a final Hill-type model (EC3) |
| 12 | (Günther et al., 2018) | Review / modelling approach on linking microscopic and macroscopic scales including a Hill-type modelling approach | Does not build a final Hill-type model (EC3) |
| 13 | (Schmitt et al., 2019) | Narrative review on the Hill-type modelling of the dynamics of skeletal muscles | Does not build a final Hill-type model (EC3) |

*Table 5: Studies investigating the Hill-type modelling approach*

| # | Reference | Important features | Reason why not eligible |
|---|---|---|---|
| 14 | (Audu & Davy, 1985) | Milestone study about the influence of Hill-type modelling complexity on predicted quantities | No modelling contribution (EC6), mainly inspired from previous studies (Hatze, 1978) |
| 15 | (Brown et al., 1996) | Comparison between modelling approaches for FL, FV, PEE, SEE and PEE2 and final model validation | Does not include AD (EC4) |
| 16 | (Perreault et al., 2003) | Assesses the link between Hill model prediction accuracy and MU firing rate level | No explicit description of the Hill-type model (EC4) |
| 17 | (Scovil & Ronsky, 2006) | Sensitivity study | No modelling contribution (EC6) |
| 18 | (Albracht & Arampatzis, 2006) | Sensitivity study | No modelling contribution (EC6) |
| 19 | (Redl et al., 2007) | Sensitivity study | Does not include AD (EC4) |
| 20 | (Siebert et al., 2008) | Comparison between Maxwell and Voigt approaches | No modelling contribution (EC6) |
| 21 | (Xiao & Higginson, 2010) | Sensitivity study | No explicit description of the Hill-type model (EC4) |
| 22 | (Blümel et al., 2012) | Sensitivity study | Another study from the same year used as eligible model |
| 23 | (Hasson & Caldwell, 2011) | Sensitivity study | No modelling contribution (EC6) |
| 24 | (Romero & Alonso, 2016) | Comparison of three models (van Soest & Bobbert, 1993; Thelen, 2003; Silva & Ambrosio, 2003) | No modelling contribution (EC6) |
| 25 | (Lemaire et al., 2016) | Comparison of the validity of Hill and Huxley-type muscle tendon models | No modelling contribution (EC6) |
| 26 | (Carbone et al., 2016) | Sensitivity study | No explicit description of the Hill-type model (EC4) |
| 27 | (Bayer et al., 2017) | Sensitivity study | No modelling contribution (EC6) |
| 28 | (Hamburger, 2017) | Comparison between Hill- and Huxley-type models | MSc dissertation |
| 29 | (Bujalski et al., 2018) | Sensitivity study | No explicit description of the Hill-type model (EC4) |
| 30 | (Sun et al., 2018) | Assessing the impact of FLA on model predictions | No modelling contribution (EC6) |
| 31 | (Harischandraid et al., 2019) | Comparison of 5 modelling approaches of the ADs | Does not build a full Hill-type model (EC3) |

*Table 6: Non-eligible Hill studies but presenting an innovative modelling contribution to the field*

| # | Reference | Important features | Reason why not eligible |
|---|---|---|---|
| 32 | (Clark & Stark, 1974) | First Hill study proposing a 1$^{st}$ order ODE expression for the AD | Joint angle state variable (EC5) |
| 33 | (Hof & Van den Berg, 1981) | First Hill-type model reported to use processed EMGs as neural inputs | Joint angle state variable (EC5) |
| 34 | (Woittiez et al., 1984) | Index of architecture and 3D volumetric representation of a muscle Hill-type model | No explicit AD (EC4) |
| 35 | (Winters & Stark, 1988) | Proposes a fibre type-dependent expression for the FVC relationship | No explicit mathematical description of the CE properties (EC4) |

| | | | |
|---|---|---|---|
| 36 | (van Ingen Schenau et al., 1988) | First Hill-type study modelling the in-series viscoelastic properties with an SDE | Does not include AD (EC4) |
| 37 | (Van Ruijven & Weijs, 1990) | Interesting approach with EMGs | No modelling contribution (EC6) |
| 38 | (Legreneur et al., 1996) | Connects a phenomenological model of MURD and ED to a Hill-type model | No explicit mathematical description of the controller (EC4) |
| 39 | (Wexler et al., 1997) | Custom model of AD and ADC<br>Includes an SDE | Does not include a CE with CE properties (EC4) |
| 40 | (Meijer et al., 1998) | Includes an SDE<br>First reported Hill model including Force Depression | Does not include AD (EC4) |
| 41 | (Forcinito et al., 1998) | First reported 'Hill model' including Residual Force Enhancement<br>Includes Force Depression | Does not include AD or CE properties (EC4) |
| 42 | (Riener et al., 1996; Riener & Fuhr, 1998) | Fatigue<br>First Hill-time model involving splines<br>MURD from FES | Complementary studies, that individually have no explicit mathematical description of the CE or passive properties (EC4) |
| 43 | (Rosen et al., 1999) | First reported Hill-type model connected to a model of neural network<br>FVA | Does not include AD (EC4) |
| 44 | (Stroeve, 1999) | FVL and FVA<br>Early study modelling muscle impedance with a Hill-type model | No explicit mathematical description of the NE properties (AD) (EC4) |
| 45 | (Wang & Buchanan, 2002) | Development of the neural network – Hill-type model approach | No explicit mathematical description of the CE properties (EC4) |
| 46 | (Till et al., 2008) | Models RFE | Does not include AD (EC4) |
| 47 | (Rode et al., 2009) | Models FD and RFE | Does not include AD (EC4) |
| 48 | (Geyer & Herr, 2010) | Open-source implementation | No modelling contribution (EC6) |
| 49 | (Van Den Bogert et al., 2011) | Implicit formulation of musculoskeletal dynamics leading to new numerical methods and optimal control<br>Formulation of a new lump state variable | No modelling contribution (EC6) |
| 50 | (Kosterina et al., 2012) | Phenomenological models of FD and RFE | No explicit mathematical description of the Hill model (EC4) |
| 51 | (Gerus et al., 2012) | Subject-specific SEE relationship obtained from in vivo ultrasound measurements – first reported study to connect this with a Hill-type model | No explicit mathematical description of the CE properties (EC4) |
| 52 | (Wakeling et al., 2012) | Derivation of a Hill rheological structure with 2 CEs to account for fibre type recruitment pattern | No complete description of the model (EC4) |
| 53 | (Nowshiravan et al., 2012) | Use of a fuzzy-genetic Hill-type model | Does not include AD (EC4) |
| 54 | (McGowan et al., 2009; McGowan et al., 2012) | Models FD and RFE | Does not include AD, and does not provide explicit description of the remaining properties (EC4) |
| 55 | (Haeufle et al., 2012) | Open-source implementation<br>Advanced model of the SDE<br>Explicit FVE relation for the eligible model from Günther et al. (2007) | No explicit mathematical description of the AD (EC4) |
| 56 | (Smith & Hunter, 2013; Smith & Hunter, 2014) | Proposes an Ogden material PEE<br>FVL, FVA | Does not include AD (EC4) |

| | | | |
|---|---|---|---|
| 57 | (Siebert et al., 2014) | Introductory work to (Siebert et al., 2018) on the inclusion of transversal loads in Hill-type models | Improved in (Siebert et al., 2018) by the same author (EC6) |
| 58 | (Ovesy et al., 2016) | Proposes an equivalent linear damping characterization in linear and nonlinear force-stiffness models | No modelling contribution (EC6) |
| 59 | (De Groote et al., 2017) | Considers short-range stiffness | No explicit mathematical description of the Hill model (EC4) |
| 60 | (Sartori et al., 2017) | First Hill-type model receiving processed HDEMGs as neural inputs | No modelling contribution (EC6) |
| 61 | (Lai et al., 2018) | Mass<br>Investigation of the improved accuracy brought by 2 CEs for slow and fast fibres | No explicit mathematical description of the Hill model (EC4)<br>No modelling contribution (EC6) |
| 62 | (Penasso & Thaller, 2018) | Proposes a new modelling approach in making model parameters fatigue-dependent | No modelling contribution (EC6) |
| 63 | (Heinen et al., 2019) | Fibre type-dependent FV relationship | Does not include AD (EC4) |
| 64 | (Guo et al., 2020) | Introduction to mass-flowing Hill-type muscle models | No modelling contribution (EC6) |

### SM 8: Results – Detailed results for the completeness assessment

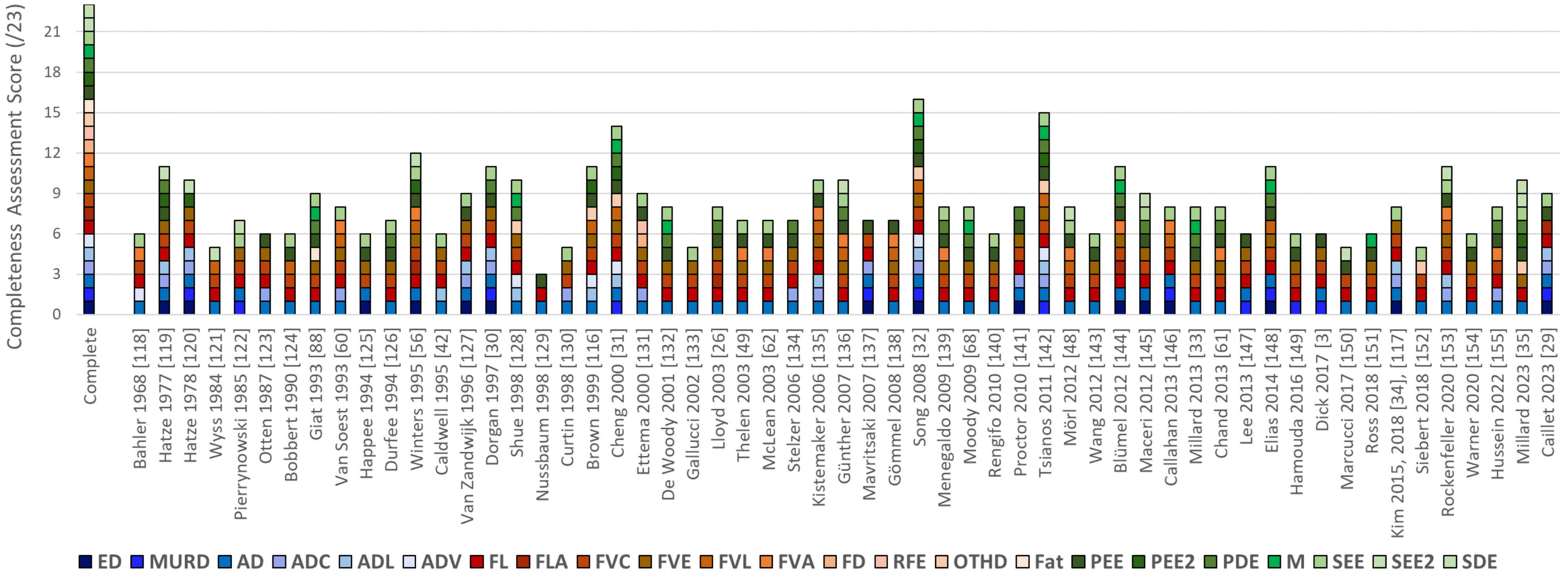

*Figure 2: Bar graph of the **completeness** assessment of the **eligible** studies. This is a detailed version of the bar graph reported in Fig. 4 of the main manuscript. Studies are reported in a chronological order. Shades of blue: NE, of red: CE, of green: passive elements. For consistency, the references in brackets [X] are those from the main manuscript.*

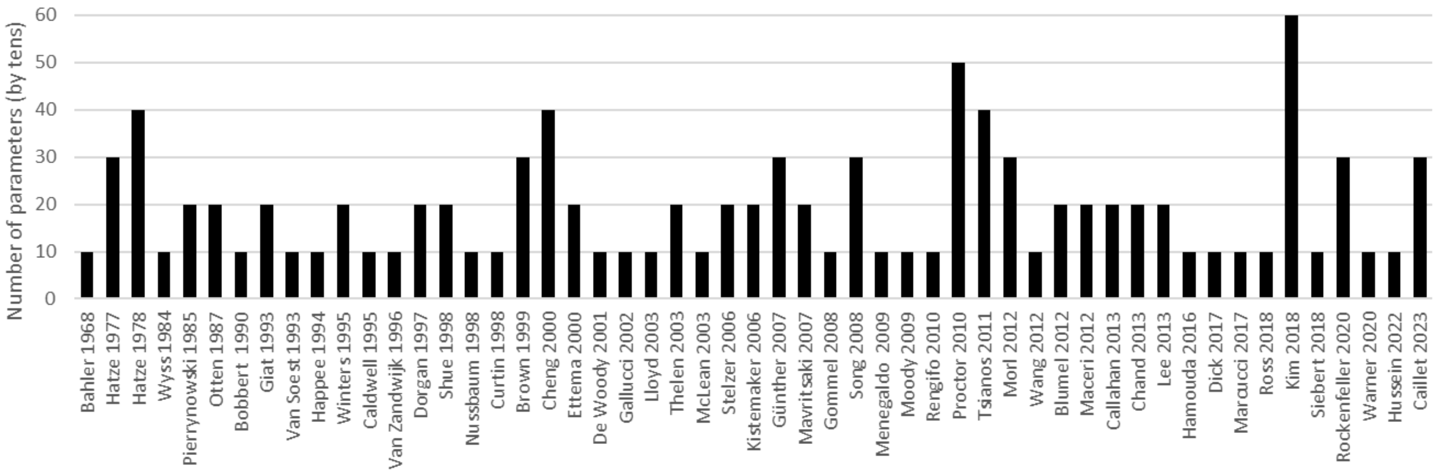

*Figure 3: Order of magnitude of the number of parameters used to define the Hill-type model in each **eligible** study. A paper including N parameters as read in the graph will include between N-4 and N+4 parameters. Are considered the parameters defining the properties of the model, and the three scaling parameters counted once even in case of a multi-muscle MSK model*

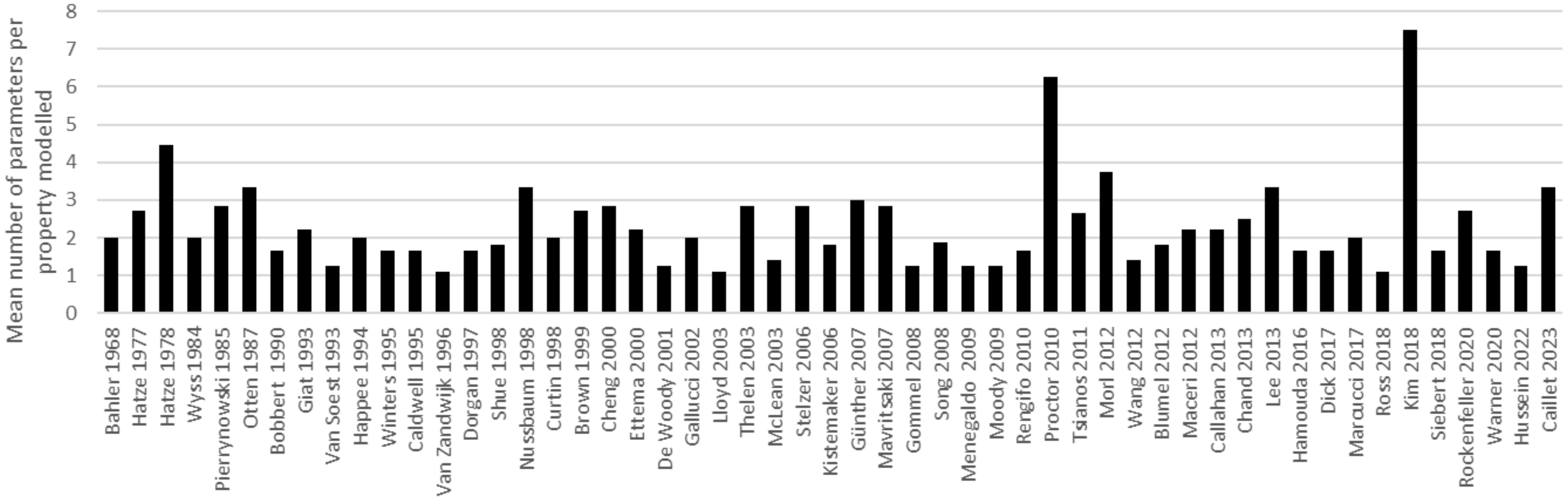

*Figure 4: Average number of parameters used to describe one property of the **eligible** models.*

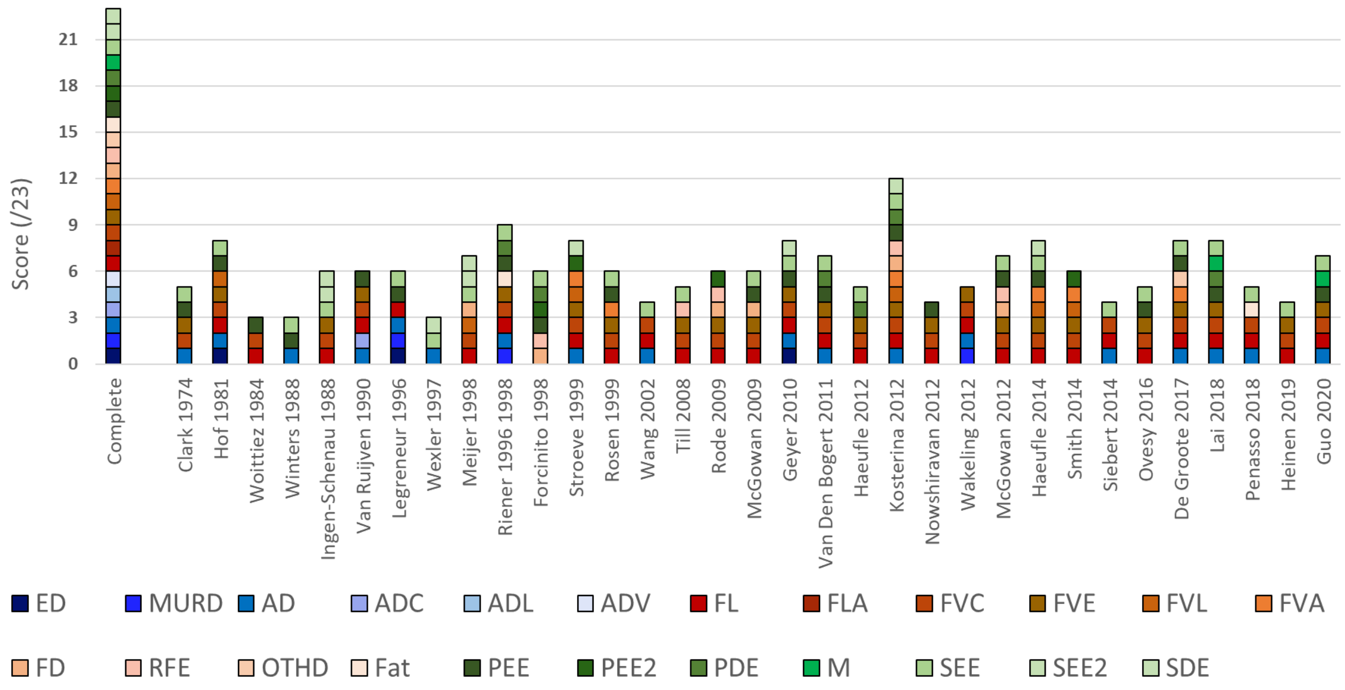

*Figure 5: Bar graph of the completeness assessment of the 33* ***non-eligible but innovative*** *Hill-type models reported in Table 6.*

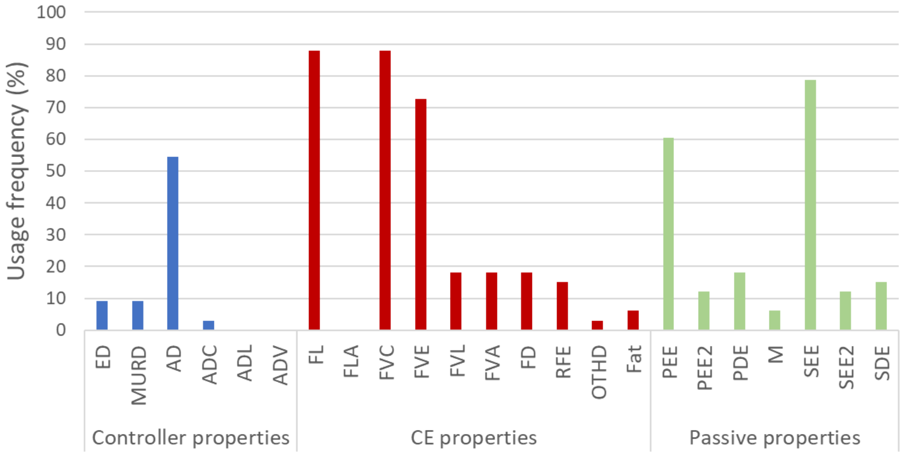

*Figure 6: Frequency of property modelling among the 33* ***non-eligible but innovative*** *studies in Table 6. As for the eligible studies, the five standard properties (AD, FL, FV, PEE, SEE) are the most widely modelled.*

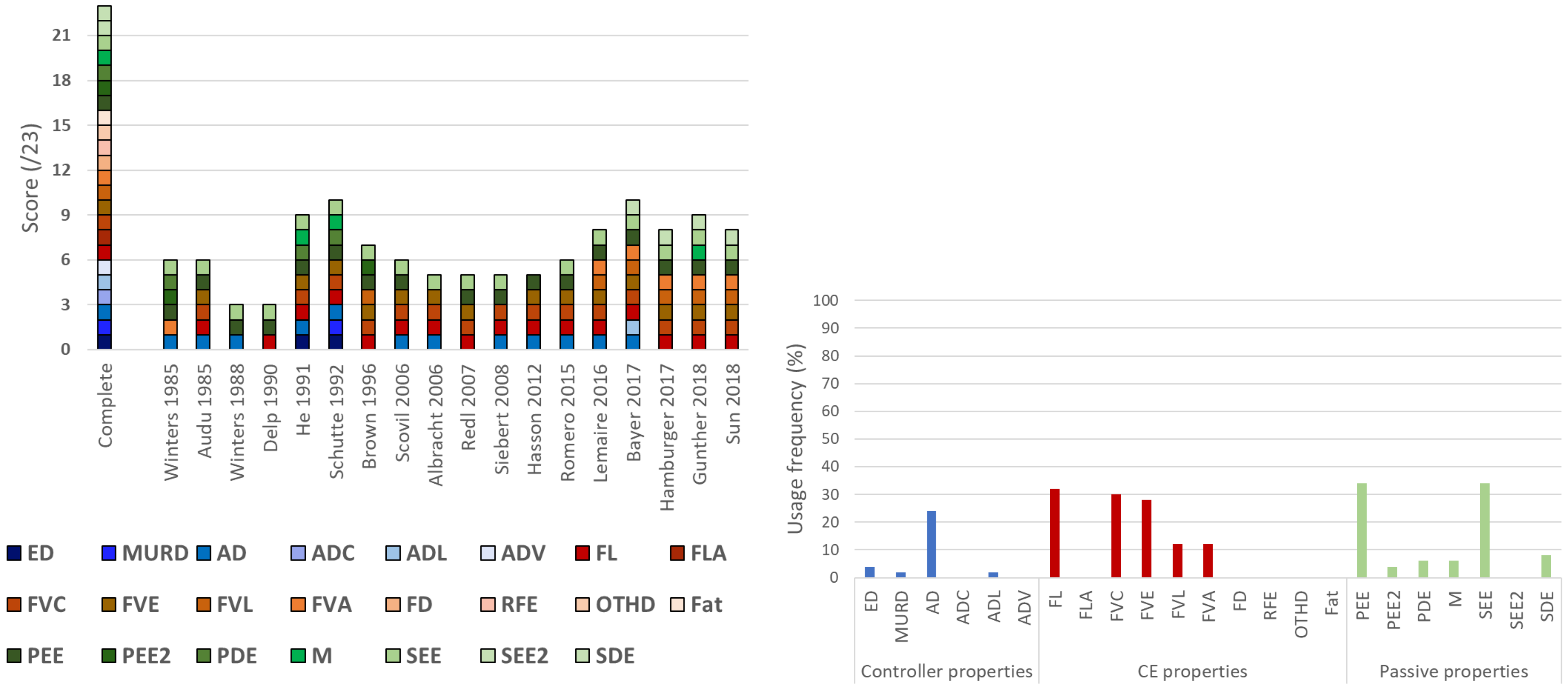

*Figure 7: Bar graph (left) and frequency of property modelling (right) resulting from the completeness assessment of some 18 __**non-eligible**__ Hill-type models reported in Tables 4 and 5.*

### SM 9: Results - Rheological structures proposed in the eligible studies

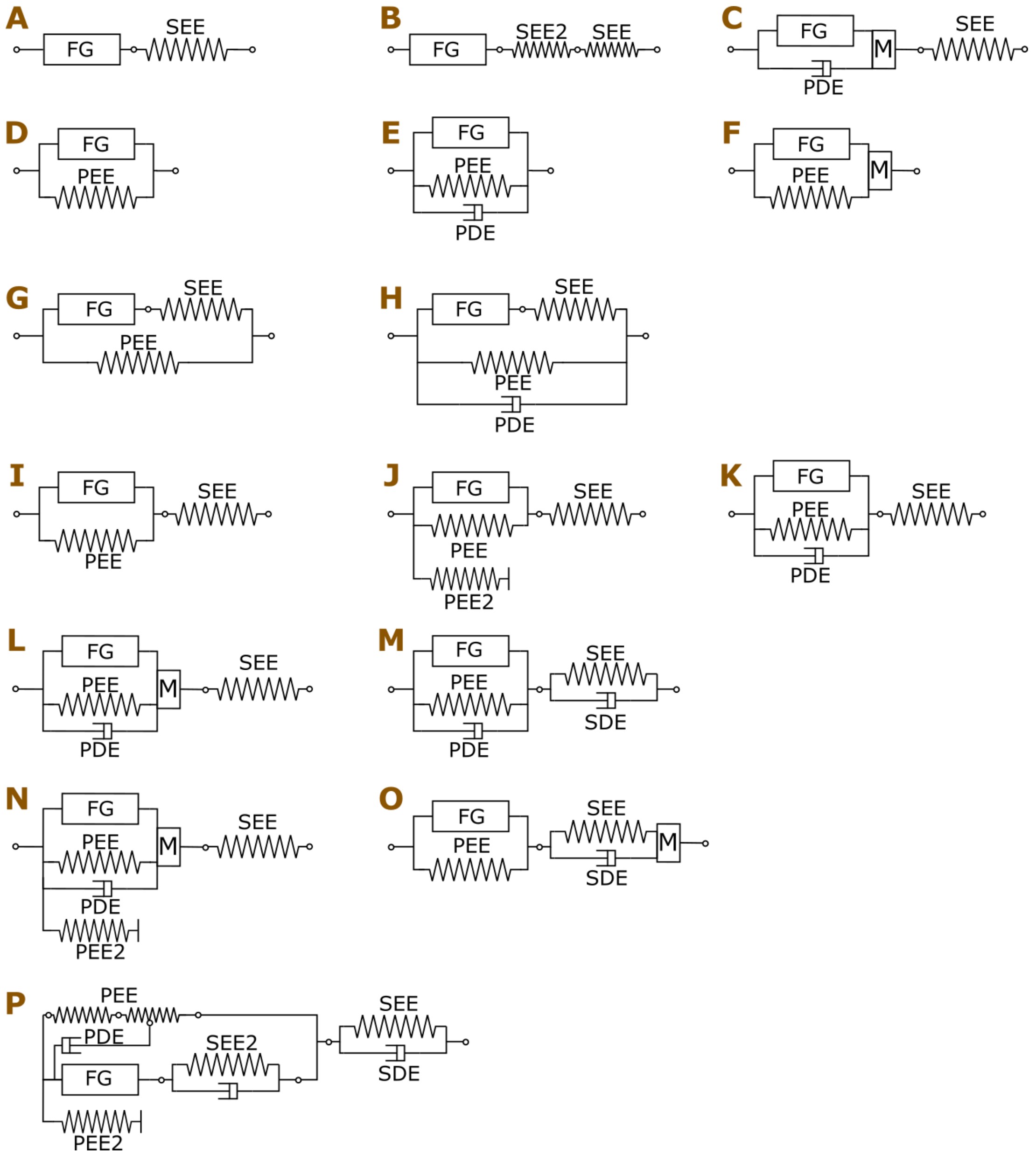

*Figure 8: Rheological structures proposed in the eligible studies and identified in Table 7. The structures in the first row (A, B, C) do not include a PEE, the second row (D, E, F) a SEE, the third row (G, H) reports Maxwell-Hill-type structures (a SEE and a PEE in parallel), while the following rows (I-O) report Kelvin-Voigt-Hill-type structures (a SEE and a PEE in-series) with increasing numbers of rheological elements. For clarity, pennation angles are not represented.*

.

*Table 7: the rheological structures of the eligible models are identified with the letters A-O from Figure 8.*

| # | Study | Ref (in the main manuscript) | Ref (in the SM) | Rheological structure in Figure 8 |
|---|---|---|---|---|
| 1 | Bahler 1968 | [118] | (Bahler, 1968) | **I** |
| 2 | Hatze 1977 | [119] | (Hatze, 1977) | **H** |
| 3 | Hatze 1978 | [120] | (Hatze, 1978) | **H** |
| 4 | Wyss 1984 | [121] | (Wyss & Pollak, 1984) | **A** |
| 5 | Pierrynowski 1985 | [122] | (Pierrynowski & Morrison, 1985) | **B** |
| 6 | Otten 1987 | [123] | (Otten, 1987) | **D** |
| 7 | Bobbert 1990 | [124] | (Bobbert & van Ingen Schenau, 1990) | **I** |
| 8 | Giat 1993 | [88] | (Giat et al., 1993) | **L** |
| 9 | Van Soest 1993 | [60] | (van Soest & Bobbert, 1993) | **A** |
| 10 | Happee 1994 | [125] | (Happee, 1994) | **A** |

| | | | | |
|---|---|---|---|---|
| 11 | Durfee 1994 | [126] | (Durfee & Palmer, 1994) | **H** |
| 12 | Winters 1995 | [56] | (Winters, 1995) | **I** |
| 13 | Caldwell 1995 | [42] | (Caldwell, 1995) | **A** |
| 14 | Van Zandwijk 1996 | [127] | (Van Zandwijk et al., 1996) | **I** |
| 15 | Dorgan 1997 | [30] | (Dorgan & O'Malley, 1997) | **H** |
| 16 | Shue 1998 | [128] | (Nussbaum & Chaffin, 1998) | **C** |
| 17 | Nussbaum 1998 | [129] | (Nussbaum & Chaffin, 1998) | **D** |
| 18 | Curtin 1998 | [130] | (Curtin et al., 1998) | **A** |
| 19 | Brown 1999 | [116] | (Brown et al., 1999) | **J** |
| 20 | Cheng 2000 | [31] | (Cheng et al., 2000) | **N** |
| 21 | Ettema 2000 | [131] | (Ettema & Meijer, 2000) | **G** |
| 22 | De Woody 2001 | [132] | (DeWoody et al., 2001) | **L** |
| 23 | Gallucci 2002 | [133] | (Gallucci & Challis, 2002) | **A** |
| 24 | Lloyd 2003 | [26] | (Lloyd & Besier, 2003) | **K** |
| 25 | Thelen 2003 | [49] | (Thelen, 2003) | **I** |
| 26 | McLean 2003 | [62] | (McLean, Scott G. et al., 2003) | **I** |
| 27 | Stelzer 2006 | [134] | (Stelzer & Von Stryk, 2006) | **E** |
| 28 | Kistemaker 2006 | [135] | (Kistemaker et al., 2006) | **I** |
| 29 | Günther 2007 | [136] | (Günther et al., 2007) | **M** |
| 30 | Mavritsaki 2007 | [137] | (Mavritsaki et al., 2007) | **D** |
| 31 | Gömmel 2008 | [138] | (Gömmel et al., 2007) | **F** |
| 32 | Song 2008 | [32] | (Song et al., 2008) | **N** |
| 33 | Menegaldo 2009 | [139] | (Menegaldo & Oliveira, 2009) | **K** |
| 34 | Moody 2009 | [68] | (Moody et al., 2009) | **L** |
| 35 | Rengifo 2010 | [140] | (Rengifo et al., 2010) | **I** |
| 36 | Proctor 2010 | [141] | (Proctor & Holmes, 2010) | **E** |
| 37 | Tsianos 2011 | [142] | (Tsianos et al., 2011) | **N** |
| 38 | Mörl 2012 | [48] | (Mörl et al., 2012) | **I, M** |
| 39 | Wang 2012 | [143] | (Wang et al., 2012) | **J** |
| 40 | Blümel 2012 | [144] | (Blümel et al., 2012) | **L** |
| 41 | Maceri 2012 | [145] | (Maceri et al., 2012) | **M** |
| 42 | Callahan 2013 | [146] | (Callahan et al., 2013) | **A** |
| 43 | Millard 2013 | [33] | (Millard et al., 2013) | **K** |
| 44 | John 2013 | [61] | (John et al., 2013) | **K** |
| 45 | Lee 2013 | [147] | (Lee et al., 2013) | **D** |
| 46 | Elias 2014 | [148] | (Elias et al., 2014) | **L** |
| 47 | Hamouda 2016 | [149] | (Hamouda et al., 2016) | **G** |
| 48 | Dick 2017 | [3] | (Dick et al., 2017) | **D** |
| 49 | Marcucci 2017 | [150] | (Marcucci et al., 2017) | **G** |
| 50 | Ross 2018 | [151] | (Ross et al., 2018) | **O** |
| 51 | Kim 2015, 2018 | [34], [117] | (Kim et al., 2015; Kim & Kim, 2018) | **A** |
| 52 | Siebert 2018 | [152] | (Siebert et al., 2018) | **A** |
| 53 | Rockenfeller 2020 | [153] | (Rockenfeller et al., 2020) | **M** |
| 54 | Warner 2020 | [154] | (Warner et al., 2020) | **I** |
| 55 | Hussein 2022 | [155] | (Hussein et al., 2022) | **K** |
| 56 | Millard 2023 | [35] | (Millard et al., 2023) | **P** |
| 57 | Caillet 2023 | [29] | (Caillet et al., 2023) | **I** |

### SM 10: Results - Inheritance diagrams and mathematical descriptions of the properties of the eligible models

In this section, inheritance diagrams are proposed for the AD, FV, FL, PEE, and SEE relationships. Only the eligible models are considered in this section, as well as historical models that inspired the eligible models. These diagrams are useful for tracking the modelling approaches taken in a study back to their source study. In the diagrams, the green colour is used for the source studies. The yellow colour is used for studies that take inspiration from the source studies but slightly adapted the mathematical equations. White is for the studies that used the modelling approach described in the source study. For each property, when a main trend is observable, the corresponding mathematical expressions are reproduced in the main text and commented.

In the following, the excitation, calcium concentration and active states are respectively referred to as $u(t), \gamma(t)$ and $a(t)$. **All other parameters are constants consistent with the notations of the corresponding studies. Please refer to the original studies for the values of the specific parameters used in the equations.**

#### a. Activation dynamics

##### i. Calcium-dependent activation dynamics - ADC

In the following, the equations for calcium transient depict the time evolution of the representative calcium concentration $\gamma(t)$ in the sarcoplasm of the muscle fibres. The steady-state transfer function embodies the sigmoid steady-state experimental relationship $F$-pCa between isometric tetanic force $F$ and calcium concentration in the muscle fibre. The transient attachment dynamics represent the transient phenomenon of cross-bridge attachment.

In this section:

- **Key state variables, function of time ($\boldsymbol{t}$)**:
    - $u(t)$: some neural signal / stimulation frequency
    - $\gamma(t)$ : representative calcium concentration in the sarcoplasm of the muscle fibres
    - $a(t)$: activation state
    - $l^{FG}(t)$: length of the Force Generator (FG)
- All other parameters are **constant parameters** related to time constants of calcium diffusion, shape factors of sigmoid functions, or (maximum) levels of calcium concentration.

Hill-type studies in the literature dominantly modelled the ADC after two studies (see Figure 9):

- (Hatze, 1977) - Model 2 (M2 in Figure 9):
    - Calcium transient due to a normalized stimulation frequency $\nu$:

$$\frac{d\gamma}{dt} = m(c\nu - \gamma) \tag{1}$$

    - Steady-state transfer function $\rho_n$ with dependency on muscle length $l^M$:

$$a = \frac{a_0 + [\rho_n(l^{FG})\gamma]^n}{1 + [\rho_n(l^{FG})\gamma]^n}$$

$$\rho_n(l^{FG}) = k_n \frac{k-1}{\frac{k}{l^{FG}} - 1} \tag{2}$$

- (Otten, 1987):
    - Calcium transient due to a neural signal $u$:

$$\frac{d\gamma}{dt} = c(\gamma_{max} u - \gamma) \tag{3}$$

    - Steady-state transfer function:

$$a_{ss} = \frac{\gamma^{2.6}}{\gamma_{0.5}^{2.6} + \gamma^{2.6}} \tag{4}$$

- Transient attachment dynamics:

$$\frac{da}{dt} = c_2(1 + \frac{a - a_{ss}}{2})(a_{ss} - a) \quad (5)$$

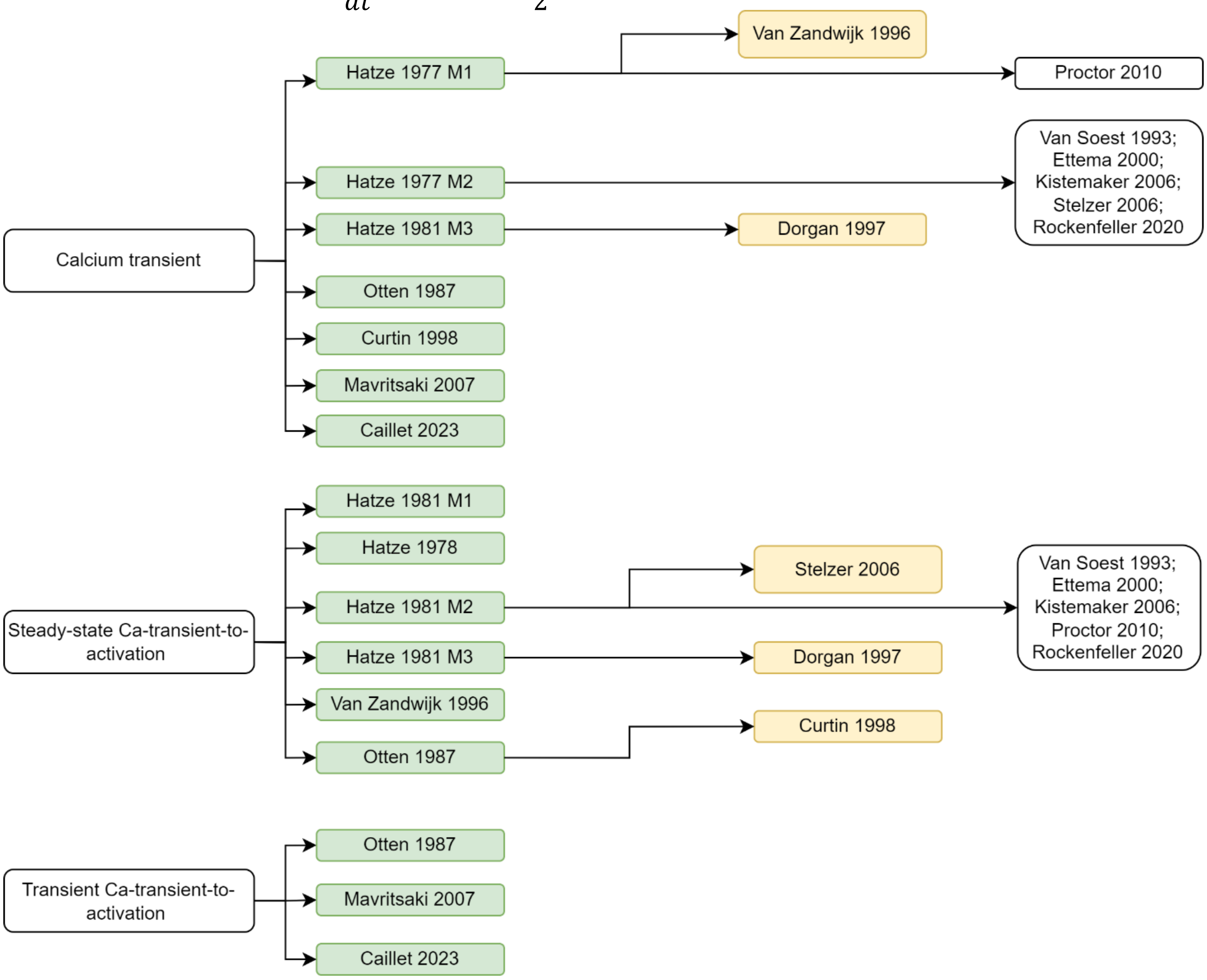

*Figure 9: Inheritance tree diagram for the Calcium-dependent models of Activation Dynamics (ADC) among the eligible studies.*

##### ii. Activation Dynamics as Steady-state transfer functions

In case of EMG-driven simulation, the most common trend is to estimate the muscle activation $a(t)$ providing the excitation state $u(t)$, that is usually obtained from processed and filtered EMGs as input, with a steady-state experimental transfer function accounting for the linear or non-linear EMG-to-force relationship. The most observed steady-state transfer function in the field is from Potvin et al. (1996) and is reproduced in equation (6).

- **State variables, function of time** $(t)$:
  - $u(t)$: excitation state
  - $a(t)$: activation state
- **Main constant parameter**: $A$ – shape factor defining the steepness of the sigmoid

$$a(t) = \frac{e^{Au} - 1}{e^{A} - 1} \quad (6)$$

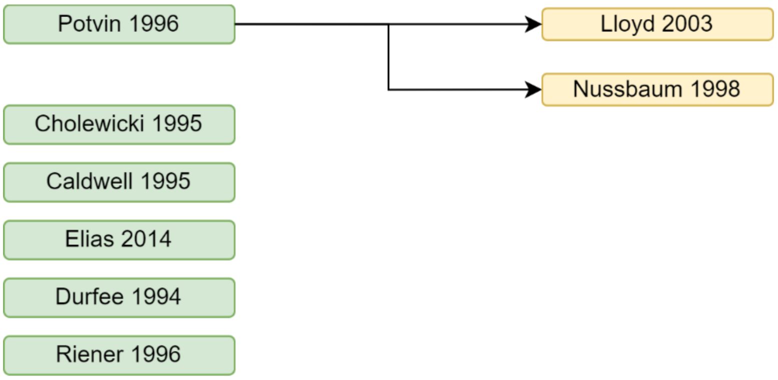

*Figure 10: Inheritance tree diagram for the Activation Dynamics modelled as a steady-state transfer function among the eligible studies*

##### iii. Activation Dynamics as 1st-order ODEs

Most Hill-type studies have been lumping the mechanisms occurring during the ADs (calcium release from the sarcoplasmic reticulum, calcium diffusion in the sarcoplasm, Calcium-troponin binding, cross-bridge attachment and detachment, cross-bridge power stroke…) with a single phenomenological first-order ODE. Various mathematical expressions have been proposed (see Figure 11) for representing the dependency of muscle activation $a(t)$ to the neural command $u(t)$. In the following are replicated the most common mathematical expressions along with their source study. The widely-used 1st-order ODE from Zajac (1989) reproduced below was compared in a sensitivity study (Rockenfeller et al., 2015) to the classic ADC formulation described previously in Hatze (1977).

> In this section:
> - **State variables, function of time ($t$)**:
>   - $u(t)$: excitation state
>   - $a(t)$: activation state
> - **Main constant parameters**: $\tau_X$ and $c_X$ represent parameters related to time constants. In some relationships, the $\tau_X, c_X$ values at activation and deactivation are not equal.

Most common mathematical expressions (see Figure 11):

- (Winters & Stark, 1988):

$$\begin{cases} \frac{da}{dt} = \frac{1}{\tau_a}(u-a) \quad ; \quad u > a \\ \frac{da}{dt} = \frac{1}{\tau_{dea}}(u-a) \quad ; \quad u \leq a \end{cases} \tag{7}$$

- (Zajac, 1989):

$$\frac{da}{dt} = \frac{u}{\tau_a} - a\left[\frac{1}{\tau_a}(\frac{\tau_a}{\tau_{dea}} + \left(1 - \frac{\tau_a}{\tau_{dea}}\right)u)\right] \tag{8}$$

- (He et al., 1991):

$$\frac{da}{dt} = (u-a)(c_1 u + c_2) \tag{9}$$

With similar notations (i.e., some $c, c_1, c_2$ parameters related to time constants and gains), these three modelling approaches can be re-written as:

- (Winters & Stark, 1988): $\frac{da}{dt} = c(u-a)$
- (Zajac, 1989): $\frac{da}{dt} = c\big(u - a(c_1 u + c_2)\big)$
- (He et al., 1991): $\frac{da}{dt} = (u-a)(c_1 u + c_2)$

Some studies also use active state dependent (de)activation time constants (represented by an activation-dependent function $f(a)$) based on Winters (1995):

$$\begin{cases} \dfrac{da}{dt} = \dfrac{1}{f(a)}(u-a) \\ \tau(a) = \tau_a(k+a) \quad ; \quad u > a \\ \tau(a) = \dfrac{\tau_{dea}}{(k+a)} \quad ; \quad u \leq a \end{cases} \tag{10}$$

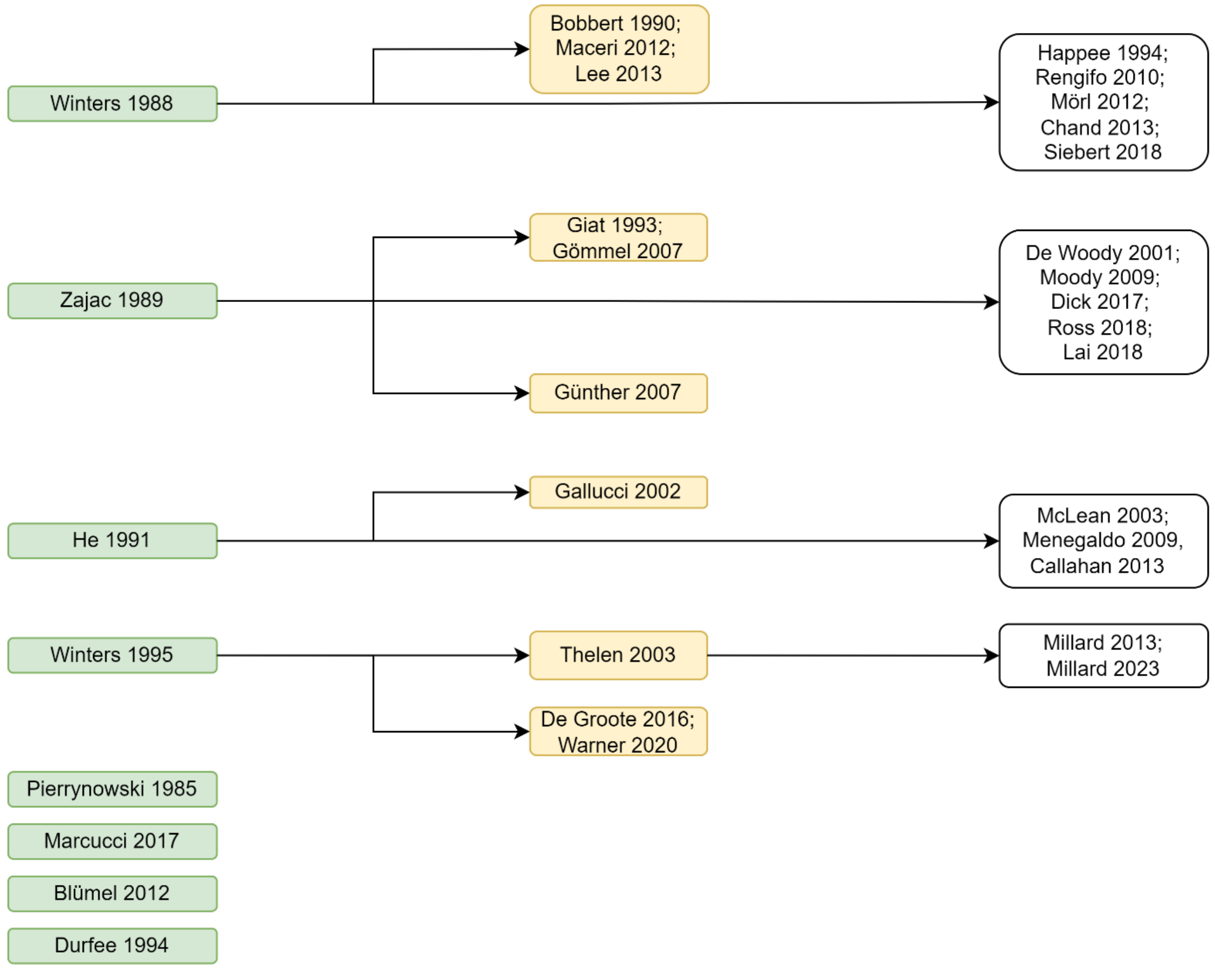

*Figure 11: Inheritance tree diagram for the Activation Dynamics modelled as a 1st-order differential equation among the eligible studies*

#### b. Force-Length relationships - FL

All the eligible studies considered an instantaneous FL relationship between the length of the Force Generator (FG), which is often taken as the fibre length in the eligible studies, and isometric force, except the work from Hamouda et al. (2016), where the issue of the instability of the descending limb of the relationship is accounted for by considering for the fibre length recorded at the onset of the contraction. Numerous different mathematical expressions have been proposed to describe the FL relationship as shown Figure 12 and reviewed in Rockenfeller & Günther (2017).

In this section:

- **State variables, function of time $(\boldsymbol{t})$:**
  - $l(t)$: normalized length of the FG – calculated as $l = \frac{l^{FG}}{l_0^{FG}}$, where $l^{FG}$ is the FG length (often named muscle or fibre length) and $l_0^{FG}$ the optimal FG length (often named optimal muscle or fibre length)
  - $\Delta l(t)$: normalized length variation of the FG – calculated as $\Delta l = \frac{l^{FG}-l_0^{FG}}{l_0^{FG}}$
  - $f_l(l), f_l(\Delta l)$: force-length factor
  - $a(t)$: activation state
- **Main constant parameters:**
  - $l_0^M$: optimal length of the Force Generator (FG)
  - $c, k_X, W, w, c$: shape parameters defining the width of the bell-shaped FL relationship and/or the steepness/curviness of its branches

As reported in Figure 12, the most popular formulations of the FL relationship are:

- The quadratic expressions from
  - (van Soest & Bobbert, 1993):
$$f_l(\Delta l) = 1 + c - 2c \cdot \Delta l + c \cdot (\Delta l)^2 \quad (11)$$
  - (Caldwell, 1995):
$$f_l(l) = k_1 + k(l - k_1)^2 \quad (12)$$
  - (Gallucci & Challis, 2002):
$$f_l(\Delta l) = 1 - \left(\frac{\Delta l}{W}\right)^2 \quad (13)$$
- The gaussian-like expressions from:
  - (Bahler, 1968):
$$f_l(\Delta l) = \exp\left(-\left(\frac{\Delta l}{W}\right)^2\right) \quad (14)$$
  - (Otten, 1987):
$$f_l(l) = \exp\left(-\left|\frac{l^b - 1}{W}\right|^c\right) \quad (15)$$

Three studies include an active-state dependency of the optimal length $l_0^{FG}$ to their FL relationship (FLA):

- (Winters, 1995):
$$f_l(l, a) = \exp\left(-\left(\frac{l - l_0(a)}{W}\right)^2\right)$$
$$l_0(a) = 1.05 + \frac{l_1}{l_0^{FG}}(1 - a) \quad (16)$$
- (Lloyd & Besier, 2003):

Cubic spline interpolation
$$l_0(a) = 1 + 0.15(1 - a) \quad (17)$$

- (Blümel et al., 2012):

$$f_l(l,a) = A(a) \cdot \left[ \frac{1 + \sin\left(wl - \left(\frac{\pi}{2} + 2.7w\right)\right)}{2} \right]$$

$$A(a) = 15e^{-1.6w(a)} \tag{18}$$

$$w(a) = 2.5 + \frac{1}{(k(a+0.5)^2\ )^2}$$

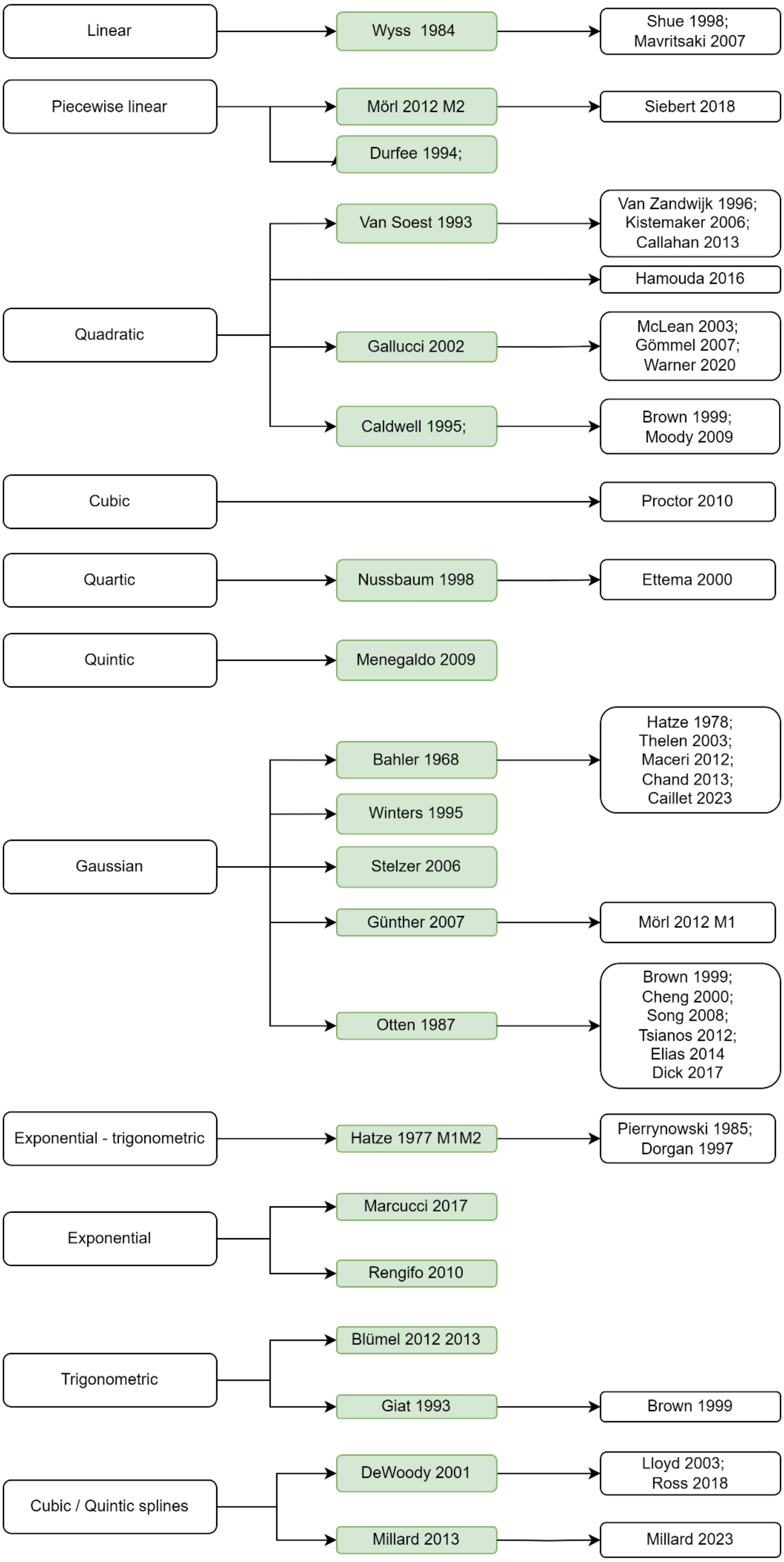

*Figure 12: Inheritance tree diagram for the FL relationship among the eligible studies.*

#### c. Force-Velocity relationships - FV

##### i. Continuous Force-Velocity relationships

A few studies modelled both the concentric (FVC) and eccentric contraction (FVE) branches of the FV relationship with a single continuous mathematical expression, either with a trigonometric function such as arctan, hyperbolic function such as Arcsinh, or with Bezier splines.

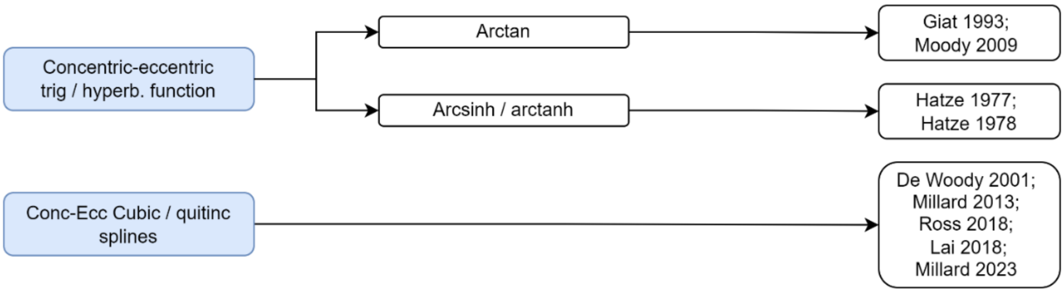

*Figure 13: Inheritance tree diagram for the complete FV relationship modelled as a continuous function among the eligible studies*

##### ii. Concentric Force-Velocity relationship - FVC

In this section, the constant parameter $a$ (in N), which partly defines the curvature of the FVC relationship, is not to be mistaken with the activation state $\boldsymbol{a}$. To avoid confusion, the activation state $\boldsymbol{a}$ is systematically reported, in this section c.ii., in bold '$\boldsymbol{a}$'.

In this Supplementary Material about FV relationships, the contraction velocity $v$ (in $m \cdot s^{-1}$) is defined negative for shortening contractions and positive for lengthening contraction, i.e.,

- $v < 0$: concentric contraction (FVC is defined for negative values of $v$)
- $v > 0$: eccentric contraction (FVE is defined for positive values of $v$)

**This choice of formalism is most commonly found in the Hill-type modelling literature**. The opposite formalism (i.e., velocity of shortening is positive) is also common in the literature, like in the pioneer studies from Hill (1938) and Mashima (1972).

To go from one formalism to another, a minus sign must be put to the $v$ variable: $v \rightarrow -v$.

- **State variables, function of time ($\boldsymbol{t}$):**
  - $v(t)$: instantaneous contraction velocity of the active tissue (muscle, fibre, etc) – defined negative for concentric contractions
  - $\bar{v}(t) = \frac{v(t)}{v_{max}}$: normalized contraction velocity ($\in [0; 1]$ as $v_{max}$ is also defined negative)
  - $f(t) = \frac{F(t)}{F_0^M}$: normalized force.
  - $f_v(v), f_v(\bar{v}), \bar{f}_v(\bar{v})$: FVC factors when the FVC expression is not normalized (Type 1 FCV), normalized by the $F_0^M$ parameters (Type 2 FVC), and by both $F_0^M$ and $v_{max}$ parameters (Type 2 FVC), respectively. These factors also represent the force $F(v), f(v)$, and $f(\bar{v})$ respectively developed by the FVC relationships.
  - $l(t)$: normalized length of the FG – calculated as $l = \frac{l^{FG}}{l_0^{FG}}$, where $l^{FG}$ is the FG length (often named muscle or fibre length) and $l_0^{FG}$ the optimal FG length (often named optimal muscle or fibre length)
  - $f_l(l)$: force-length factor – described in the previous section
  - $\boldsymbol{a}(t)$: activation state
- **Main constant parameters:**
  - $v_{max}$: maximum shortening velocity – defined negative in this formalism
  - $F_0^M$: maximum isometric force of the active tissue – defined as $P_0$ in Hill (1938)
  - $a, b, a_f$: shape factors of the FVC relationship

The FV relationship for concentric contractions (FVC) is dominantly modelled in the field of Hill-type modelling after Hill's hyperbolic formula (Hill, 1938) as shown in Figure 15. In the first equation of Hill (1938), the FVC relationship is reported as '$(F+a)(v+b) = (F_0^M + a)b = const$', where shortening contraction velocities are defined positive.

Equation (19) rewrites the latter FVC relationship from Hill (1938) when shortening contraction velocities $v$ are negative (i.e., after the $v \rightarrow -v$ transformation is achieved):

$$\begin{aligned} &(F+a)(-\boldsymbol{v}+b) = b(F_0^M + a) \\ \Leftrightarrow\ &-v(F+a) = b(F_0^M + a) - b(F+a) \\ \Leftrightarrow\ &\boldsymbol{(F+a)v = b\left(F - F_0^M\right)} \end{aligned} \tag{19}$$

Equation (19) can also be reshaped in the Type 1 FVC form by isolating the force variable $F$ (see next sub-section for details about this form).

$$\begin{aligned} &(F+a)(-\boldsymbol{v}+b) = b(F_0^M + a) \\ \Leftrightarrow\ &F(-v+b) = b(F_0^M + a) - a(-v+b) \\ \Leftrightarrow\ &\boldsymbol{F(v) = \frac{b\left(F_0^M + a\right)}{b-v} - a = \frac{bF_0^M + av}{b-v}} \end{aligned} \tag{20}$$

At the maximum concentric contraction velocity $v = v_{max}$ (negative value in the chosen formalism), the muscle cannot generate any force ($F = 0$) and we can write the following adimensional ratios, that are strictly positive according to experimental measurements:

$$\boldsymbol{\frac{b}{-v_{max}} = \frac{a}{F_0^M} \equiv a_f > 0} \tag{21}$$

**<u>Type 1 FVC relationship – unnormalized FVC relationship</u>**

Equations (22) and (23), that reproduce the result of Equation (20), describe the unnormalized FVC scaling factor $f_v(v)$ in forms that are commonly reported in the literature, and which are named 'Type 1' FVC relationship in the following.

$$F(v) = \frac{b(F_0^M + a)}{b-v} - a = f_v(v) \tag{22}$$

$$\boldsymbol{F(v) = \frac{bF_0^M + av}{b-v} = f_v(v)} \tag{23}$$

**<u>Type 2 FVC relationships – normalized by the maximum isometric force $F_0^M$ and/or by the maximum shortening velocity $v_{max}$</u>**

In the literature, Hill's FVC relationship is commonly normalised by the architectural $F_0^M$ and $v_{max}$ parameters. Those normalized relationships are named 'Type 2 FVC' relationships in the following. Let $f = \frac{F}{F_0^M} \in [0;1]$ be the normalized force as obtained by the FV relationship, $\bar{v} = \frac{v}{v_{max}} \in [0;1]$ the normalized contraction velocity, and $a_f = \frac{a}{F_0^M} > 0$ the coefficient that defines the curvature of the hyperbola according to the species and the fibre type under consideration at fixed length.

Equation (23) is normalized by the maximum isometric force $F_0^M$ as:

$$F(v) = \frac{bF_0^M + av}{b-v} = f_v(v) \tag{24}$$

$$\Leftrightarrow f(v) = \frac{F(v)}{F_0^M} = \frac{b + \dfrac{a}{F_0^M} v}{b - v} = \overline{f_v}(v)$$

$$\Leftrightarrow \boldsymbol{f(v)} = \frac{\boldsymbol{1 + a_f \dfrac{v}{b}}}{\boldsymbol{1 - \dfrac{v}{b}}} = \boldsymbol{\overline{f_v}(v)}$$

Considering in Equation (21) that $\frac{a_f}{b} = -\frac{1}{v_{max}}$, Equation (24) is further normalized by the maximum contraction velocity $v_{max}$ as:

$$f(v) = \frac{1 + a_f \dfrac{v}{b}}{1 - \dfrac{v}{b}} = \overline{f_v}(v)$$

$$\Leftrightarrow f(v) = \frac{1 - \dfrac{v}{v_{max}}}{1 + \dfrac{v}{a_f v_{max}}} = \overline{f_v}(v) \tag{25}$$

$$\Leftrightarrow \boldsymbol{f(\bar{v})} = \frac{\boldsymbol{1 - \bar{v}}}{\boldsymbol{1 + \dfrac{\bar{v}}{a_f}}} = \overline{f_v}(\bar{v})$$

This Type 2 normalized form of the FVC therefore relies on the parameter $a_f$, that varies in the range $[0.02; 0.84]$ in the literature, and the maximum shortening velocity $v_{max}$ that is often taken constant across muscles in the range $[3; 10] m \cdot s^{-1}$.

In some studies, the FVC relationship is inverted, and the velocity variable is isolated to display a first-order ordinary differential equation of the form $v = \frac{dl^{FG}}{dt} = f_v^{-1}(l^{FG})$. Inverted expressions are reported in the summary Table 8.

*Table 8: Summary of the expressions commonly taken by the FVC relationships in the literature, when they are unnormalized (Type 1) or normalized by the $F_0^M$ and $v_{max}$ parameters (Type 2), and when they are inverted. Representative curves are given for $F_0^M = 1000N, v_{max} = -10m \cdot s^{-1}, a_f = 0.2, a = 200N$, and $b = 2m \cdot s^{-1}$. Remark: $v$ and $v_{max}$ are defined negative for shortening contractions; therefore, $\bar{v} = \frac{v}{v_{max}} \in [0; 1]$.*

| SUMMARY - FVC RELATIONSHIP | | | |
|---|---|---|---|
| | Type 1 - unnormalized | Type 2 - $F_0^M$-normalized | Type 2 - $F_0^M - v_{max}$-normalized |
| Curve | 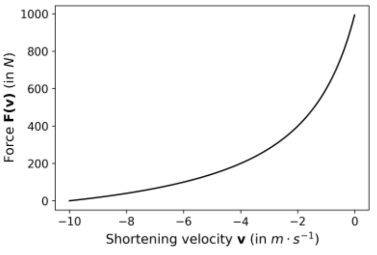 | 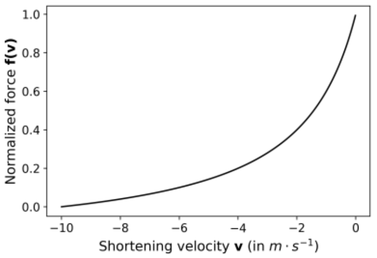 | 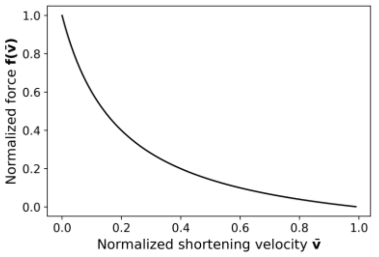 |
| Expression | $F(v) = \frac{bF_0^M + av}{b - v} = f_v(v)$ | $f(v) = \frac{1 + a_f \frac{v}{b}}{1 - \frac{v}{b}} = \overline{f_v}(v)$ | $f(\bar{v}) = \frac{1 - \bar{v}}{1 + \frac{\bar{v}}{a_f}} = \overline{f_v}(\bar{v})$ |
| Inverted expression | $v(F) = \frac{b}{a} \cdot \frac{F - F_0^M}{\frac{F}{a} + 1} = f_v^{-1}(F)$ | $v(f) = \frac{b}{a_f} \cdot \frac{f - 1}{\frac{f}{a_f} + 1} = f_v^{-1}(f)$ | $\bar{v}(f) = \frac{1 - f}{1 + \frac{f}{a_f}} = \overline{f_v^{-1}}(f)$ |

Some studies propose a nonlinear **activation-dependency and/or length-dependency of Hill's hyperbolic relationship**. In this case, the reader is invited not to make a common confusion between

- an inverted FVC relationship that is nonlinearly on activation and/or length, and writes $v = f_v^{-1}(F, l^M, a)$, and
- an inverted FVC that is simply dependent on velocity and classically scales linearly with activation and FL, but that would include (in the way it is presented) the description of the whole CE dynamics, which would write $v = f_v^{-1}(F^{CE}) = f_v^{-1}(a \cdot f_l \cdot F)$ with $f_l$ the isometric force obtained from the FL relationship as in section b..

In the following, $l = \frac{l^{FG}}{l_0^{FG}}$ is the normalized length of the FG. Most studies further develop the expression of Hill's FV relationship in imposing a zero or low constant force boundaries for shortening speed exceeding the maximum shortening velocity. These regions of high shortening speed are not investigated in the following.

**Length-dependent FVC:**

The following expressions report FVC relationships that nonlinearly scale with the normalized length $l$ of the FG, by sometimes including the FL relationship $f_l(l)$ directly in the FVC relationship.

- (Brown et al., 1999; Cheng et al., 2000; Song et al., 2008; Tsianos et al., 2011; Elias et al., 2014):

$$f(\bar{v}, l) = \frac{1 - \bar{v}}{1 + f_c(l)\bar{v}} = \overline{f_v}(\bar{v}, l) \qquad (26)$$

$$f_c(l) = (c_1 + c_2 l)$$

- (Wyss & Pollak, 1984; Hamouda et al., 2016):

In this approach, the Type 2 relationship in Equation (24) is modified to include $f_l(l)$ instead of the scalar '1' in numerator. This approach is only partially valid as stated previously (Epstein, 1998); the issue is that the maximum shortening velocity becomes length-dependent: $-v_{max}(l) = \frac{f_l(l) \cdot b}{a}$ which is contradictory with other experimental findings of a constant maximum shortening velocity at all lengths (Edman, 1979).

$$f(v, l) = \frac{f_l(l) + \frac{a_f}{b} v}{1 - \frac{v}{b}} = f_v(\bar{v}, l)$$

Or

$$f(\bar{v}, l) = \frac{f_l(l) - \bar{v}}{1 + \frac{\bar{v}}{a_f}} = \overline{f_v}(\bar{v}, l)$$

Or

$$v(f) = \frac{b}{a_f} \frac{f - f_l(l)}{\frac{f}{a_f} + 1} = f_v^{-1}(f, l) \qquad (27)$$

Or

$$\bar{v}(f) = \frac{f_l(l) - f}{1 + \frac{f}{a_f}} = \overline{f_v^{-1}}(f, l)$$

**Activation-dependent FVC:**

Here, the maximum shortening velocity is made activation-dependent to reflect the nonlinear effects of the gradual recruitment of slow to fast fibre types on the maximum shortening velocity.

(Otten, 1987; McLean, Scott G. et al., 2003; Thelen, 2003; John et al., 2013; De Groote et al., 2017): these studies make

$$f(\bar{v},\boldsymbol{a}) = \frac{1-\frac{\bar{v}}{g(\boldsymbol{a})}}{1+\frac{\bar{v}}{a_f \cdot g(\boldsymbol{a})}} = \bar{f_v}(\bar{v},\boldsymbol{a})$$

With definition of g($a$) changing across studies: (28)

- (Otten, 1987): $g(\boldsymbol{a}) = (0.35\, f^M(\boldsymbol{a}) + 0.65)$
- (Thelen, 2003): $g(\boldsymbol{a}) = 0.25 + 0.75\boldsymbol{a}$
- (McLean et al., 2003): $g(\boldsymbol{a}) = 1 - e^{-3.82a} + \boldsymbol{a}e^{-3.82}$
- (John et al., 2013): $g(\boldsymbol{a}) = 0.5(1+\boldsymbol{a})$

**Activation-length-dependent FVC:**

Some other studies make the FVC relationship nonlinearly dependent on both length $l$ and active state $a$, in which case one or both of the maximum shortening velocity and/or the curvature of the hyperbola nonlinearly vary with $l$ and/or $\boldsymbol{a}$.

- (Winters, 1995):

$$f(\bar{v},\boldsymbol{a},l) = \frac{1-\frac{\bar{v}}{g(\boldsymbol{a},l)}}{1+\frac{\bar{v}}{a_f \cdot g(\boldsymbol{a},l)}} = \bar{f_v}(\bar{v},\boldsymbol{a},l) \quad (29)$$

With:

$$g(\boldsymbol{a},l) = 0.5\big(1+\boldsymbol{a}\cdot f_l(l)\big)$$

- (van Soest & Bobbert, 1993; Kistemaker et al., 2006; Callahan et al., 2013):

$$v(f,\boldsymbol{a},l) = h(\boldsymbol{a})\frac{b}{a_f}\cdot\frac{f-1}{\frac{f}{a_f}+\frac{1}{g(l)}} = f_v^{-1}(f,\boldsymbol{a},l)$$

Or

$$f(\bar{v},\boldsymbol{a},l) = \frac{1-\frac{\bar{v}}{h(a)\cdot g(l)}}{1+\frac{\bar{v}}{a_f\cdot h(a)}} = \bar{f_v}(\bar{v},\boldsymbol{a},l)$$

With definition of $g(l)$ and h($a$) changing across studies:

- (Callahan et al., 2013): $g(l) = f_l(l) \quad ; \quad h(\boldsymbol{a}) = \boldsymbol{a}^{0.3}$ (30)

- (Van Soest & Bobbert, 1993): $g(l) = \begin{cases} f_l(l) & ; \quad l < 1 \\ 1 & ; \quad else \end{cases}$ ;
  $h(a) = \min(1, 3.33\boldsymbol{a})$

- (Kistemaker et al., 2006): $g(l) = \begin{cases} f_l(l) & ; \quad l < 1 \\ 1 & ; \quad else \end{cases}$ ;
  $h(\boldsymbol{a}) = b(1-0.9Q(\boldsymbol{a}))^2$ ;
  and
  $$\begin{cases} Q(\boldsymbol{a}) = \frac{\boldsymbol{a}-a_{crit}}{a_0 - a_{crit}} & ; \quad a < a_{crit} \\ 1 & ; \quad else \end{cases}$$

- (Günther et al., 2007; Kosterina et al., 2012; Mörl et al., 2012; Rockenfeller, R. et al., 2020):

$$v(f,\boldsymbol{a},l) = f_B(\boldsymbol{a})\frac{b}{a_f}\cdot\frac{f-1}{\frac{f}{a_f}+\frac{f_A(\boldsymbol{a})}{g(l)}} = f_v^{-1}(f,\boldsymbol{a},l)$$

Or

$$f(\bar{v},\boldsymbol{a},l) = \frac{1-\frac{f_A(\boldsymbol{a})\cdot\bar{v}}{f_B(\boldsymbol{a})\cdot g(l)}}{1+\frac{\bar{v}}{a_f\cdot f_B(\boldsymbol{a})}} = \bar{f_v}(\bar{v},\boldsymbol{a},l)$$

(31)

With

$$f_A(l) = \begin{cases} f_l(l) & ; \quad l<1 \\ 1 & ; \quad else \end{cases}$$

$$f_A(\boldsymbol{a}) = \frac{0.25}{\boldsymbol{a}}(1+3\boldsymbol{a})$$

$$f_B(\boldsymbol{a}) = \frac{1}{7}(3+4\boldsymbol{a})$$

One study (Curtin et al., 1998) proposed a **double-hyperbolic FV relationship**:

$$f(\bar{v}) = \begin{cases} \frac{1}{m}\cdot\frac{1-\bar{v}}{1+\frac{\bar{v}}{a_f}} & ; \quad 0<f<f_1 \\ 1-k_1\ln\left(1+\frac{\bar{v}}{k_2}\right) & ; \quad f_1<f<1 \end{cases} \tag{32}$$

Two studies fitted the curvature of Hill's relation with mathematical expressions according to the fraction of fast and slow fibres in the muscle into consideration, or to the relative fraction of recruited MUs. In the following, $\%_{fast}$ represents the relative fraction of fast fibres composing the muscle under consideration, and $n_{MU}(t)$ the relative number of recruited MUs at time instant t.

(Winters & Stark, 1988):

$$a_f = 0.1 + 0.4 * \%_{fast} \tag{33}$$

(Callahan et al., 2013):

$$a_f = 0.2 - 0.3n_{MU}(t) \tag{34}$$

**Example of a FVC that is nonlinearly dependent on fibre type, activation state, and normalized length**

In this exemple, the maximum shortening velocity is made nonlinearly dependent on:

- Fibre type – with the $\boldsymbol{k_{MU}(type)}$ coefficient, it is assumed that fast-type MUs ($k_{MU}=1$) reach maximum shortening velocities twice those of slow-type MUs ($k_{MU}=\frac{1}{2}$). This is adequate as muscles with respectively dominantly slow and fast fibres are usually modelled to reach $v_{max}$ values of $5m\cdot s^{-1}$ and $10m\cdot s^{-1}$, respectively.
- Activation state $\boldsymbol{a}$ – with the $\boldsymbol{h(a)=0.8+0.2\cdot a}$ coefficient, to account for activation-related mechanisms independent from the type of recruited MUs (in the same approach as other studies (Otten, 1987; Thelen, 2003; Günther et al., 2007; John et al., 2013), see above)

- Length $l$ – the maximum shortening velocity decreases with sub-optimal shorter MU lengths by $\boldsymbol{g(\bar{l})} = \begin{cases} \boldsymbol{f_{FL}(\bar{l})} & ; \quad \boldsymbol{\bar{l} < 1} \\ \boldsymbol{1} & ; \quad \boldsymbol{else} \end{cases}$ (van Soest & Bobbert, 1993; Kistemaker et al., 2006; Günther et al., 2007).

Then, the curvature of the FVC relationship is fibre type dependent, with $a_f$ values typically 1.5-to-2.5 times higher in rodent and monkey fast-type fibres and rodent and cat muscles of dominantly fast-type fibres (Spector et al., 1980; Otten, 1987; Bottinelli et al., 1991; Fitts et al., 1998). $a_f$ also linearly increases by a factor two between fibre- and muscle-scales (compare (Ranatunga, 1982; McDonald et al., 1994; Roots et al., 2007) for works on rat skinned fibres, bundles of fibres, and whole muscles). Using these observations in combination with experimental $a_f$ values obtained in human skinned fibres and whole lower limb muscles (Andersen et al., 2005; de Brito Fontana et al., 2014; Hauraix et al., 2015; Dada et al., 2018), it was chosen $a_f(slow) = 0.20$ for slow-type MUs and $a_f(fast) = 0.40$ for fast-type MUs in the human TA muscle, which is consistent with previous reviews (Wakeling et al., 2012; Dick et al., 2017). It yields Equation (35), that is plotted in Figure 14.

$$f(\bar{v}, type, \boldsymbol{a}, l) = \frac{1 - \dfrac{\bar{v}}{K(type, \boldsymbol{a}, l)}}{1 + \dfrac{\bar{v}}{a_f(type) \cdot K(type, \boldsymbol{a}, l)}} = \bar{f}_v(\bar{v}, type, \boldsymbol{a}, l) \tag{35}$$

with

$$\begin{cases} K(type, \boldsymbol{a}, l) = k_{MU}(type) \cdot h(\boldsymbol{a}) \cdot g(l) \\ k_{MU}(type) = \begin{cases} 0.5 & ; \quad slow \\ 1 & ; \quad fast \end{cases} \\ h(a) = 0.8 + 0.2 \cdot a \\ g(l) = \begin{cases} f_{FL}(l) & ; \quad l < 1 \\ 1 & ; \quad else \end{cases} \\ a_f(type) = \begin{cases} 0.20 & ; \quad slow \\ 0.40 & ; \quad fast \end{cases} \end{cases}$$

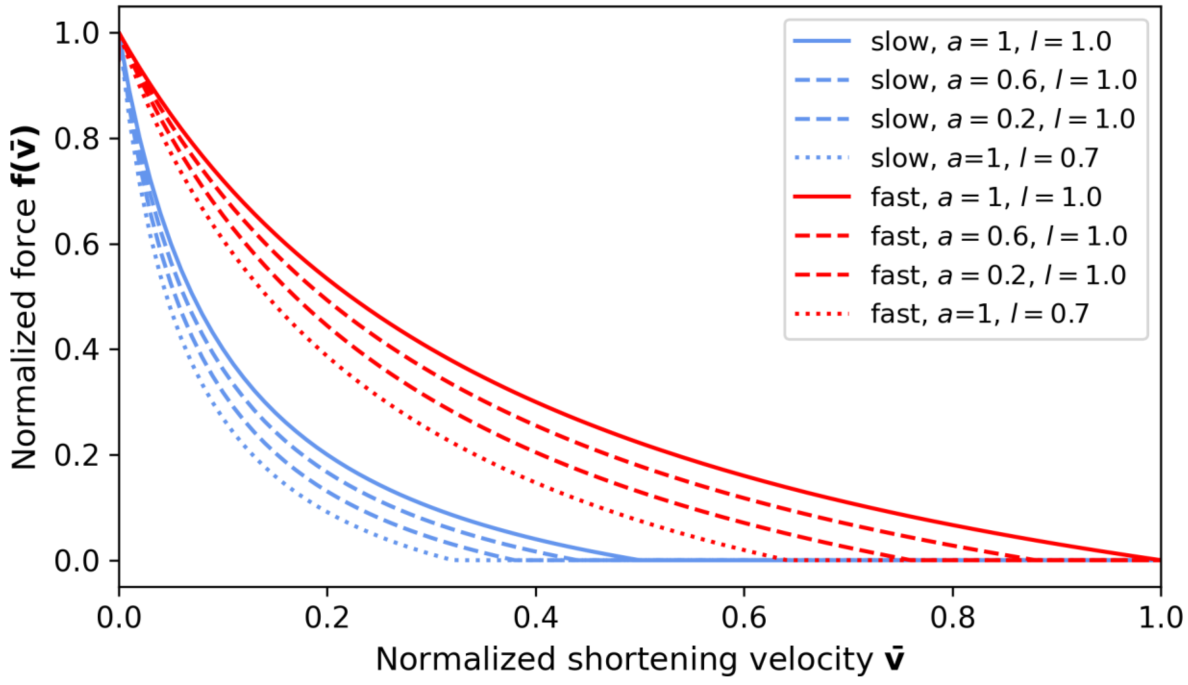

*Figure 14: Type 2 FVC relationship that nonlinearly depends on activation state* $\boldsymbol{a}$*, length* $l$*, and fibre type (slow or fast).*

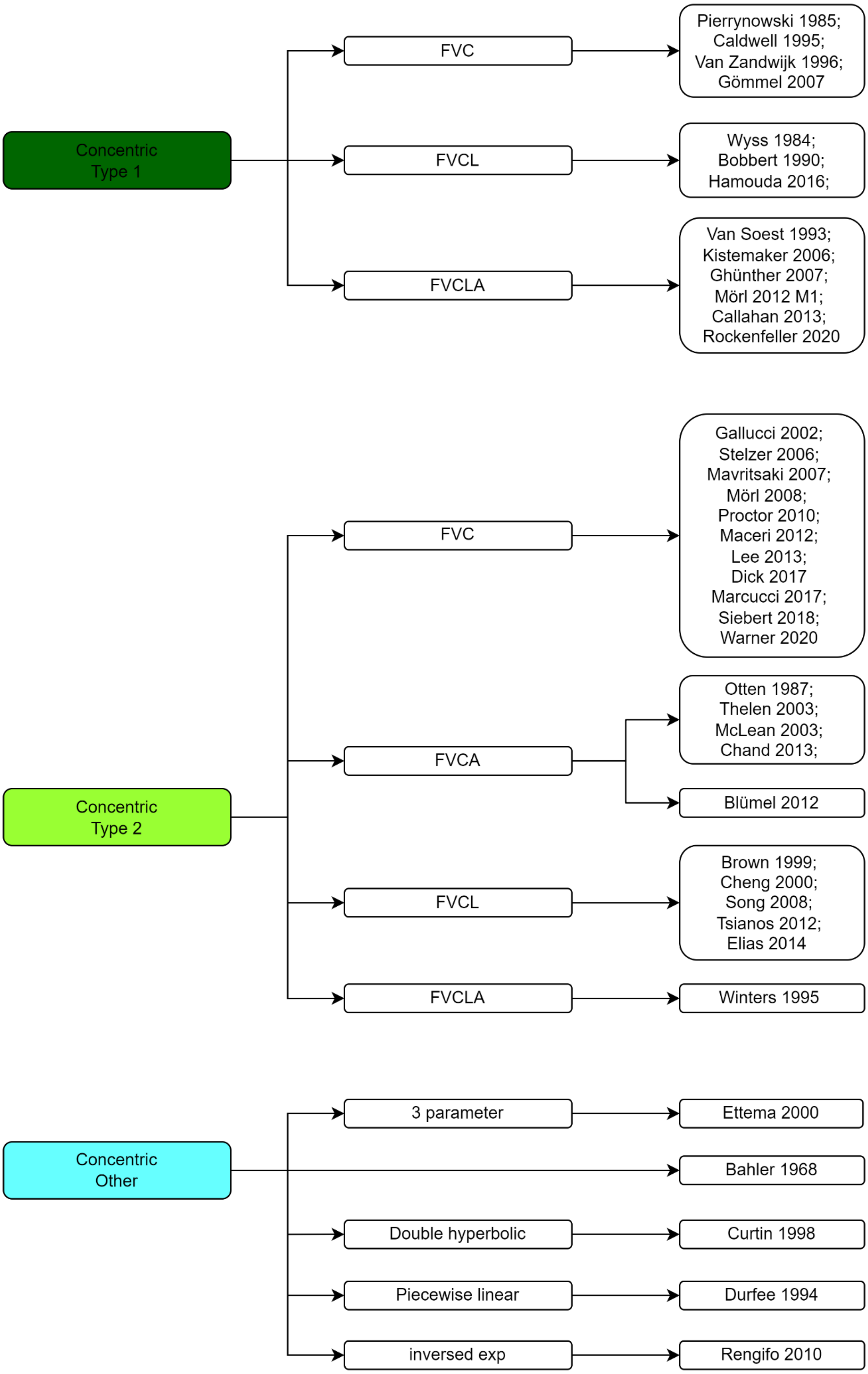

*Figure 15: Inheritance tree diagram for the complete FVC relationship among the eligible studies. Please refer to the text for the definition of Type 1 (unnormalized) and Type 2 (normalized) relationships.*

##### iii. Eccentric Force-velocity relationship - FVE

It is recalled that, in this supplementary material, the lengthening velocities $v$ of FVE's eccentric contractions are defined positive.

| |
|---|
| ➢ **State variables, function of time ($\boldsymbol{t}$):**<br>o $v(t)$: instantaneous contraction velocity of the active tissue (muscle, fibre, etc) – defined positive for eccentric contractions<br>o $\bar{v}(t) = \frac{v(t)}{v_{max}}$: normalized contraction velocity (negative for eccentric contractions, as $v_{max}$ the concentric maximum shortening velocity and is therefore defined negative)<br>o $f(t) = \frac{F(t)}{F_0^M}$: normalized force.<br>o $\overline{f_v}(\bar{v}) = f(\bar{v})$: FVE force factor when the FVE expression is normalized by both $F_0^M$ and $v_{max}$ parameters (Type 2 FVC), respectively.<br>o $l(t)$: normalized length of the FG – calculated as $l = \frac{l^{FG}}{l_0^{FG}}$, where $l^{FG}$ is the FG length (often named muscle or fibre length) and $l_0^{FG}$ the optimal FG length (often named optimal muscle or fibre length)<br>o $f_l(l)$: force-length factor – described in the previous section<br>o $\boldsymbol{a}(t)$: activation state<br>➢ **Main constant parameters:**<br>o $v_{max}$: maximum shortening velocity – defined negative in this formalism<br>o $F_0^M$: maximum isometric force of the active tissue – defined as $P_0$ in Mashima (1972)<br>o<br>o $\overline{F^{max}}$ : normalized maximum force achievable in eccentric contractions, ranging in $[1.1; 1.25]$ for humans and up to 1.8 for other species<br>o $a', b', c'$: shape factors of the FVE relationship |

Most FVE relationships are described in the eligible studies as a nonlinear curve up to a certain lengthening velocity at which the relationship becomes mainly linear or constant. The following only focuses on the nonlinear region of the relationship. As for the FVC relationship, a mathematical form is dominantly used to describe the FVE relationship as summarised in Figure 16. FVE is derived from two source studies (Mashima et al., 1972; FitzHugh, 1977) and is named 'Type 3' relationship in the following.

This inversed hyperbolic Type 3 form was initially reported (Mashima et al., 1972) as $'(2F_0^M - F + a')(-\bar{v} + b') = b'(F_0^M + a')$ with the normalized lengthening speeds $\bar{v}$ taking negative values, as in the formalism chosen for this supplementary material. The transformation $\bar{v} \rightarrow -\bar{v}$ was therefore not necessary here. As in Equations (19) to (25) for FVC, normalization by $F_0^M$ is applied below.

$$(2F_0^M - F + a')(b' - \bar{v}) = b'(F_0^M + a') \qquad (36)$$

$$\Leftrightarrow -F\left(b' - \bar{v}\right) = b'\left(F_0^M + a'\right) - \left(2F_0^M + a'\right)\left(b' - \bar{v}\right)$$

$$\Leftrightarrow -F\left(b' - \bar{v}\right) = -b'F_0^M + \bar{v}\left(2F_0^M + a'\right)$$

$$\Leftrightarrow F(\bar{v}) = \frac{b'F_0^M - \bar{v}\left(2F_0^M + a'\right)}{b' - \bar{v}}$$

$$\Leftrightarrow f(\bar{v}) = \frac{F(\bar{v})}{F_0^M} = \frac{1 - \frac{\left(2 + \frac{a'}{F_0^M}\right)}{b'}\bar{v}}{1 - \frac{\bar{v}}{b'}}$$

Setting for this study $\overline{F^{max}} = 2 + \frac{a'}{F_0^M}$, it yields the classic Type 3 form of the FVE relationship reported in Equation (37), that is very similar to the Type 2 relationship of the FVC:

$$f(\overline{\boldsymbol{v}}) = \frac{\mathbf{1} - \overline{\boldsymbol{F^{max}}}\frac{\overline{\boldsymbol{v}}}{\boldsymbol{b'}}}{\mathbf{1} - \frac{\overline{\boldsymbol{v}}}{\boldsymbol{b'}}} = \overline{\boldsymbol{f}_{\boldsymbol{v}}}(\overline{\boldsymbol{v}}) \tag{37}$$

From experiments, it can be obtained $b' = a_f' \frac{\overline{F^{max}}-1}{a_f'+1}$.

Equation (37) is also found reported with other expressions, reported in Equation (38).

$$f(\overline{\boldsymbol{v}}) = \overline{\boldsymbol{F^{max}}} - (\overline{\boldsymbol{F^{max}}} - \mathbf{1})\frac{\mathbf{1} + \overline{\boldsymbol{v}}}{\mathbf{1} - \frac{\overline{\boldsymbol{v}}}{\boldsymbol{b'}}} = \overline{\boldsymbol{f}_{\boldsymbol{v}}}(\overline{\boldsymbol{v}})$$

or

$$\boldsymbol{f}(\overline{\boldsymbol{v}}) = \frac{\mathbf{1} - \boldsymbol{c}'\overline{\boldsymbol{v}}}{\mathbf{1} - \frac{\overline{\boldsymbol{v}}}{\boldsymbol{b'}}} = \overline{\boldsymbol{f}_{\boldsymbol{v}}}(\overline{\boldsymbol{v}}) \tag{38}$$

with

$$c' = \overline{F^{max}} - 1 + \frac{\overline{F^{max}}}{b'}$$

Equation (37) can be inverted as reported in Equation (39).

$$\overline{\boldsymbol{v}} = \boldsymbol{b}'\frac{\mathbf{1} - \boldsymbol{f}}{\boldsymbol{f} - \overline{\boldsymbol{F^{max}}}} = \boldsymbol{f}_{\boldsymbol{v}}^{-\mathbf{1}}(\boldsymbol{f}) \tag{39}$$

Some studies use a length-dependent and/or activation-dependent Type 3 relationship.

**Length-dependent FVE:**

- (Brown et al., 1999; Cheng et al., 2000; Song et al., 2008; Tsianos et al., 2011; Elias et al., 2014):

$$f(\bar{v}, l) = \frac{1 - f_e(l)\frac{\bar{v}}{b}}{1 - \frac{\bar{v}}{b}} \tag{40}$$

$$f_e(l) = (a_1 + a_2 l + a_3 l^2)$$

**Activation-dependent FVE:**

- (Thelen, 2003; John et al., 2013):

$$f(\bar{v}, \boldsymbol{a}) = \frac{1 - \overline{F^{max}}\frac{\bar{v}}{h(\boldsymbol{a})b'}}{1 - \frac{\bar{v}}{h(\boldsymbol{a})b'}}$$

$$b' = \frac{\overline{F^{max}} - 1}{2 + \frac{2}{A}} \tag{41}$$

  - (Thelen, 2003): $h(\boldsymbol{a}) = 0.25 + 0.75\boldsymbol{a}$
  - (John et al., 2013): $h(\boldsymbol{a}) = 0.5(1 + \boldsymbol{a})$

**Activation-length-dependent FVE:**

- (Winters, 1995):

$$f(\bar{v}, \boldsymbol{a}, l) = \frac{1 - \frac{\bar{v}}{g(\boldsymbol{a}, l)}}{1 - \frac{\bar{v}}{g(\boldsymbol{a}, l)(\overline{F^{max}} - 1)c}} \quad (42)$$

with

$$g(\boldsymbol{a}, l) = 0.5\big(1 + \boldsymbol{a} \cdot f_l(l)\big)$$

##### iv. $C^2$ continuity between concentric and eccentric branches

Numerous studies use a hyperbolic shape to describe the FVE relationship and enforce a $C^2$ condition at the point of isometric contraction that joins the concentric and eccentric branches, following the work from Van Soest & Bobbert (1993).

The eccentric branch is written as a 3-parameter $c_1, c_2, c_3$ expression:

$$\bar{v} = -\frac{c_1}{f + c_2} - c_3 = \frac{-c_1 - c_2 c_3 - c_3 f}{c_2 + f} \Leftrightarrow f(\bar{v}) = \frac{-(c_1 + c_2 c_3) - c_2 \bar{v}}{\bar{v} + c_3} \quad (43)$$

The 3 parameters are obtained to respect the following conditions:

1. $f_v^{conc}$ and $f_v^{ecc}$ are continuous for $\bar{v} = 0$
2. At $\bar{v} = 0$, the ratio between the eccentric and concentric derivatives is specified by the $df$ coefficient: $\frac{df_{ecc}}{d\bar{v}} = sf * \frac{df_{conc}}{d\bar{v}}$
3. $\lim_{\bar{v} \to \infty} f_v^{ecc} = \overline{F^{max}}$

From condition 3, $c_2 = -\overline{F^{max}} * f_l(l)$.

From condition 2, $c_1 = \frac{b * (f_l(l) + c_2)^2}{\big(a + f_l(l)\big) * sf}$

From condition 1, $c_3 = \frac{c_1}{f_l(l) + c_2}$

Reinserting the initial expression, $\bar{v} = c_1 \left[\frac{1}{1 - \overline{F^{max}}} - \frac{1}{f - \overline{F^{max}}}\right] = \frac{b\big(\overline{F^{max}} - 1\big)}{sf(1+a)} \frac{f - 1}{\overline{F^{max}} - f} = d_1 \frac{f - 1}{\overline{F^{max}} - f}$

Inverting the last expression, a Type 3 expression can be observed:

$$f(\bar{v}) = \frac{1 + \overline{F^{max}} \, \overline{\frac{v}{d_1}}}{1 + \frac{\bar{v}}{d_1}}$$

$$d_1 = \frac{b(\overline{F^{max}} - 1)}{sf(1 + a)} \quad (44)$$

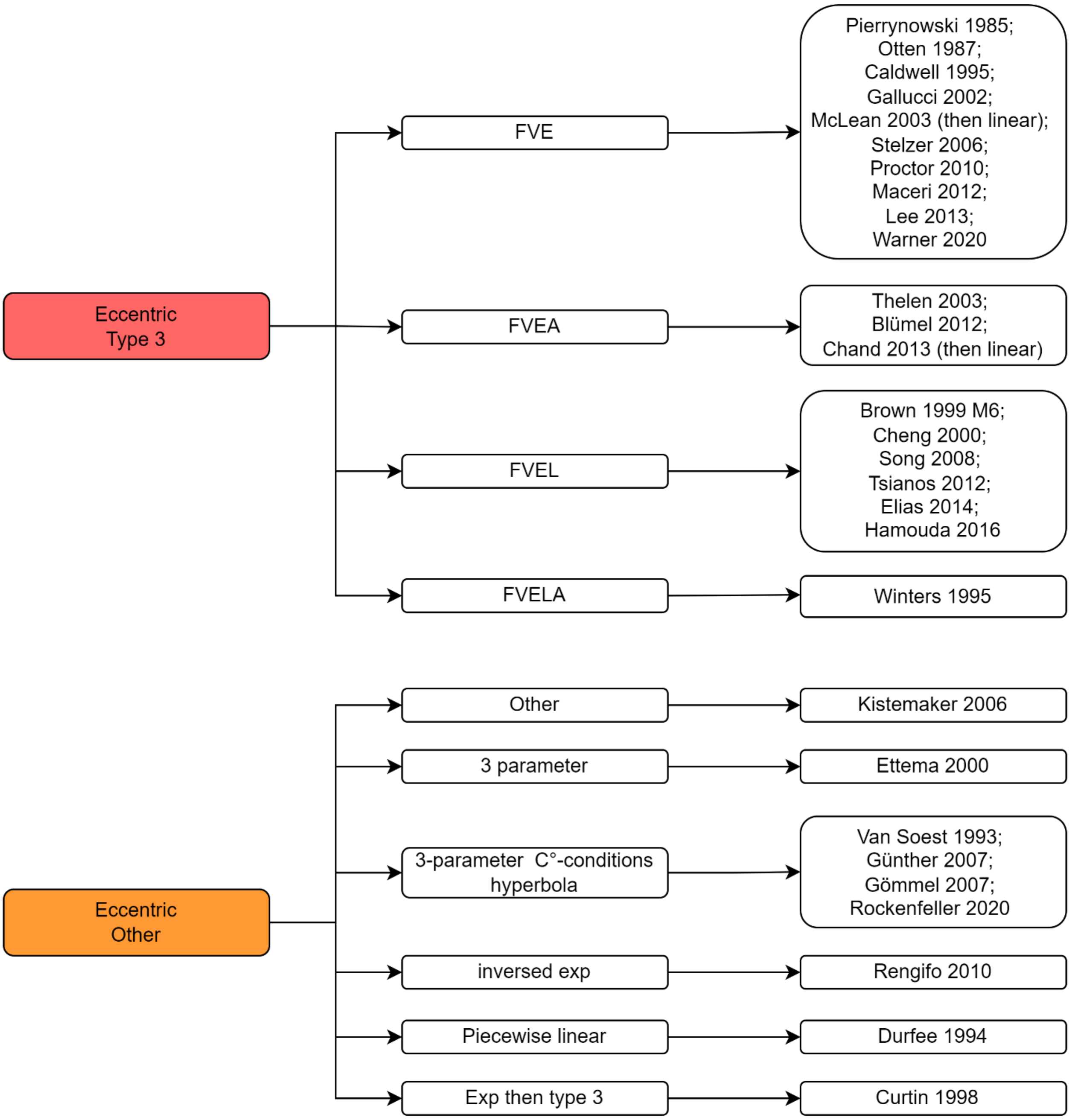

*Figure 16: Inheritance tree diagram for the complete FVE relationship among the eligible studies. Please refer to the text for the definition of the Type 3 relationship.*

#### d. Parallel Elastic Element - PEE

In all the eligible studies, the behaviour of the PEE is represented as an instantaneous force-displacement relationship applied to a tissue of constant cross-section. In the following, it is assumed that

- The length of the PEE $l^{PEE}$ equals the length of the Force Generator (FG), i.e., $l^{PEE} = l^{FG}$,
- the slack length of the PEE occurs at the FG's optimal length $l_0^{FG}$

In this section:

- **State variables, function of time ($t$):**
  - $\Delta l(t)$: normalized length variation of the FG around the slack length– calculated as $\Delta l = \frac{l^{FG} - l_0^{FG}}{l_0^{FG}}$
  - $f^{PEE}(\Delta l)$: PEE factor – normalized force of the PEE
- **Main constant parameters:**
  - $k_X$: shape parameters defining the PEE's stiffness, i.e., the steepness and/or curviness of the exponential shape of the force-displacement relationship

Four main mathematical descriptions of the PEE are encountered in the literature, as shown Figure 17 and reproduced below. In the literature, the PEE factor $f^{PEE}$ is set to 0 or to negligible values for PEE lengths below $l_0^{FG}$. Therefore, in the following, the equations are only reported for $\Delta l > 0$.

- (van Soest & Bobbert, 1993):

$$f^{PEE}(\Delta l) = k(\Delta l)^2 \quad (45)$$

- (Van Ruijven & Weijs, 1990):

$$f^{PEE}(\Delta l) = k_1 \exp(k_2 \Delta l) \quad (46)$$

- (Hatze, 1977):

$$f^{PEE}(\Delta l) = k(e^{k_1 \Delta l} - 1)$$

Or

$$f^{PEE}(\Delta l) = \frac{e^{\frac{k}{k_1}\Delta l} - 1}{e^k - 1} \quad (47)$$

- (Brown et al., 1999):

$$f^{PEE}(\Delta l) = c_1 \cdot k \cdot \ln\left[e^{\frac{l^M - l_1}{k}} + 1\right] \quad (48)$$

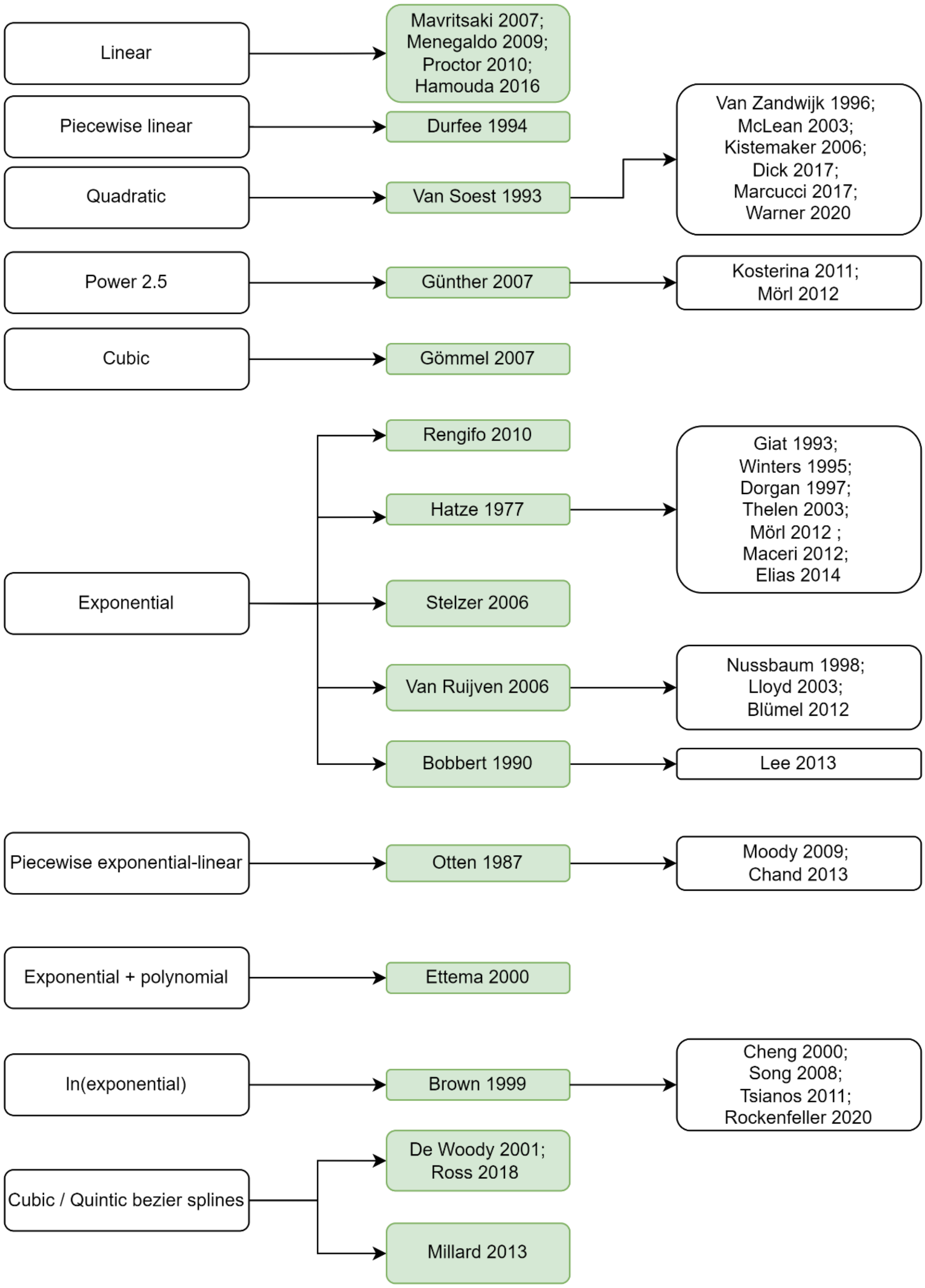

*Figure 17: Inheritance tree diagram for the modelling of the properties of the PEE among the eligible studies.*

#### e. Series Elastic Element - SEE

In the vast majority of the eligible studies, the SEE describes the physiological tendon. In those studies, the behaviour of the SEE is described with an instantaneous stress-strain relationship applied to a tendon of constant cross-section, with mathematical expressions similar to the ones used for describing the PEE, but different values for the coefficients. One study proposed a multiscale model of the SEE (Maceri et al., 2012), and one study considered the cross-sectional area of the tendon to derive its stiffness (Pierrynowski & Morrison, 1985).

In this section:

- **State variables, function of time $(t)$:**
  - $l^{SEE}(t)$: length of the SEE
  - $\varepsilon(t)$: strain of the SEE as defined by the normalized variation of SEE length around the SEE's slack length $l_s^{SEE}$, often called tendon slack length ($l_s^T$) in the studies – calculated as $\varepsilon(t) = \frac{l^{SEE}(t)-l_s^{SEE}}{l_s^{SEE}}$
  - $f^{SEE}(\varepsilon)$: SEE factor – stress in the SEE
- **Main constant parameters:**
  - $k_X$: shape parameters defining the SEE's stiffness, i.e., the steepness and/or curviness of the exponential shape of the stress-strain relationship
  - $\varepsilon_0$: strain of the SEE at peak isometric force

Four main mathematical descriptions of the PEE are encountered in the literature, as shown Figure 18 and reproduced below. In the literature, the SEE factor $f^{SEE}$ is set to 0 or to negligible values for SEE lengths below the SEE's slack length $l_s^{SEE}$. Therefore, in the following, the equations are only reported for $\varepsilon > 0$.

- (Bobbert & van Ingen Schenau, 1990):

$$f^{SEE}(\varepsilon) = k\varepsilon^2 \tag{49}$$

- (Hatze, 1977; Winters & Stark, 1988):

$$f^{SEE}(\varepsilon) = \frac{e^{\frac{k}{k_1}\varepsilon} - 1}{e^k - 1} \tag{50}$$

- (Giat et al., 1993):

$$f^{SEE}(\varepsilon) = \begin{cases} k\left(e^{k_1\varepsilon} - 1\right) & ; \quad \varepsilon < \varepsilon_0 \\ k_2(\varepsilon - \varepsilon_0) + f^{SEE}(\varepsilon_0) & ; \quad \varepsilon > \varepsilon_0 \end{cases} \tag{51}$$

- (Brown et al., 1999):

$$f^{SEE}(\varepsilon) = c_1 \cdot k \cdot \ln\left[e^{\frac{\varepsilon}{k}} + 1\right] \tag{52}$$

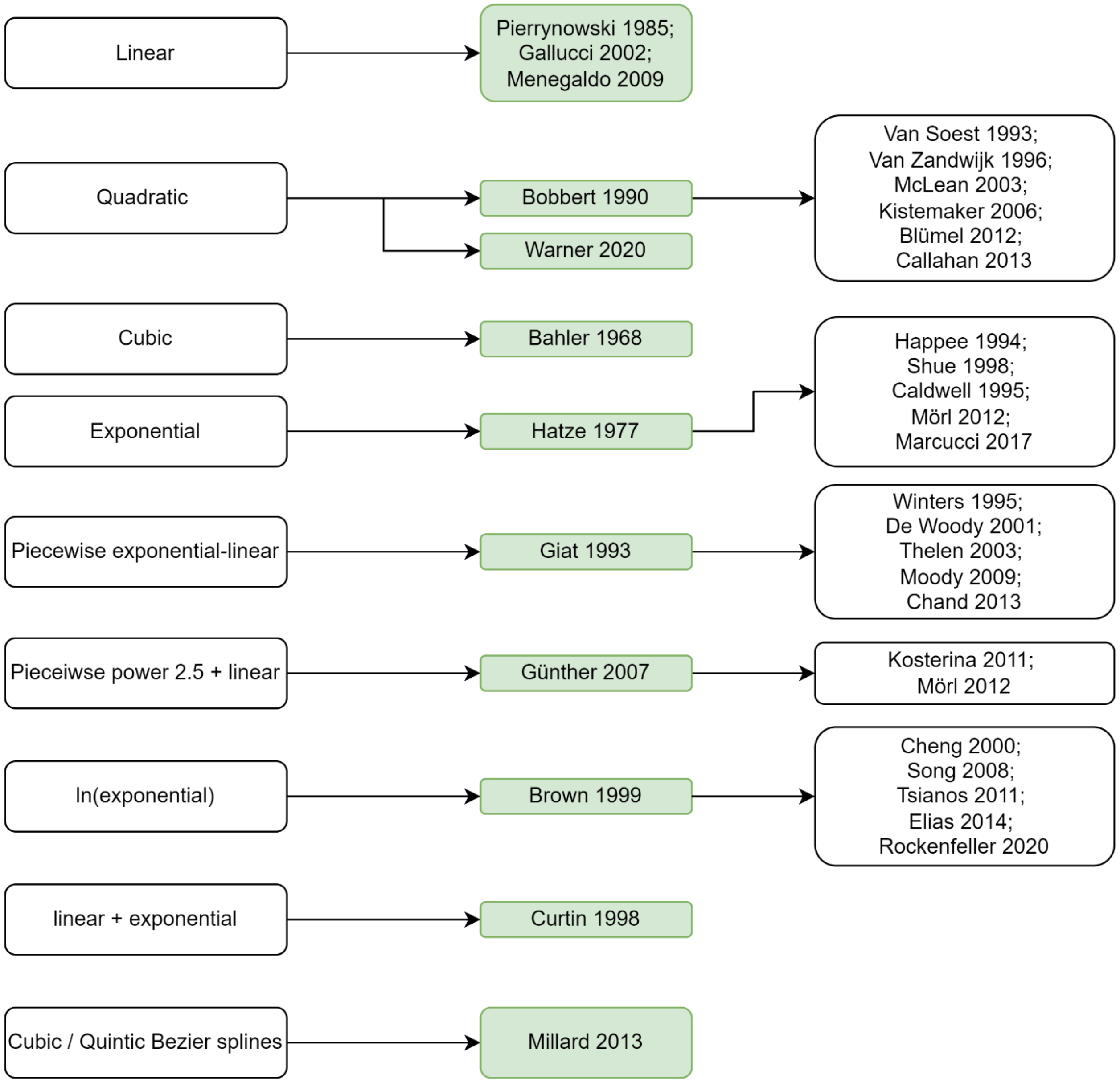

*Figure 18: Inheritance tree diagram for the modelling of the properties of the SEE among the eligible studies.*

#### SM 11: Results - Detailed results on the modelling assessment of each eligible model

The results of the scoring of the modelling assessment are provided in the bar graph Figure 19. The excel spreadsheet gathering the detail of the scoring for each question for each model is also provided as supplementary material online. In Figure 19, studies are chronologically listed. A first bar is provided for comparison, representing the scoring of a theoretical Hill-type model that fulfils all requirements for model validation, reusability, strategy, and calibration, respectively represented by shades of green, red, blue and yellow, respectively.

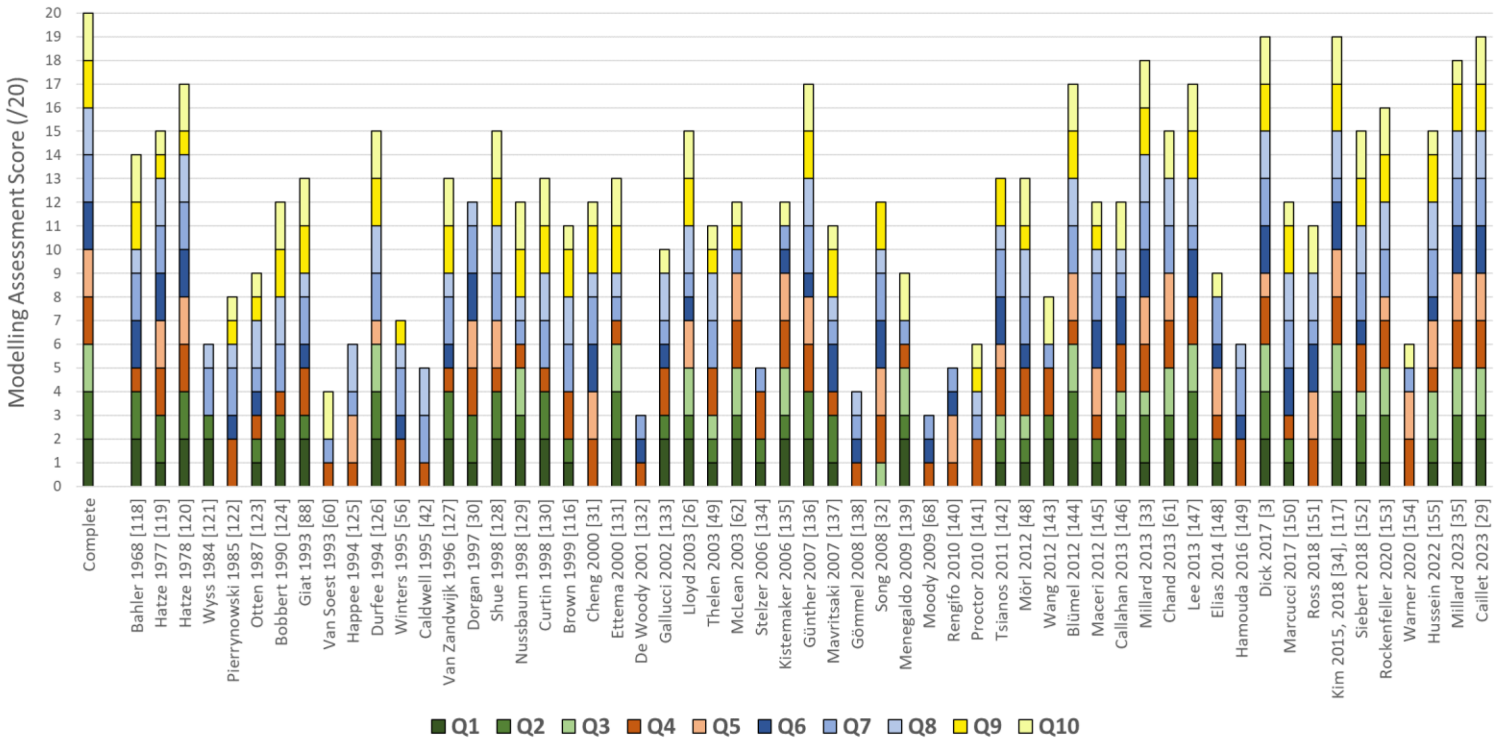

*Figure 19: Bar graph presenting the detailed results of the **modelling assessment** of the **eligible** models. This is a detailed version of the bar graph reported in Fig. 7 of the main manuscript. Shades of green: model validation, of red: model reusability, of blue: modelling choices and strategy, of yellow: parameter calibration. Each question could score 0, 1, or 2. For consistency, the references in brackets [X] are those from the main manuscript.*

### SM 12: Results – Trends in the methodological practices presented in the eligible models

*Table 9: Details on the methodological practices among the 54 eligible studies. For each of the ten questions, the most popular and/or the most advanced methodological approach is retrieved, and the corresponding references are provided.*

| Q n° | Feature | Additional precisions | Percentage of eligible studies | References |
|---|---|---|---|---|
| 1 | Ad hoc experimental data for model validation | | 44% | |
| | | Ad hoc animal experiments | 18% | (Bahler, 1968; Durfee & Palmer, 1994; Van Zandwijk et al., 1996; Curtin et al., 1998; Shue & Crago, 1998; Ettema & Meijer, 2000; Günther et al., 2007; Blümel et al., 2012; Lee et al., 2013; Kim & Kim, 2018) |
| | | Ad hoc human experiments | 26% | (Hatze, 1978; Wyss & Pollak, 1984; Bobbert & van Ingen Schenau, 1990; Giat et al., 1993; Nussbaum & Chaffin, 1998; Gallucci & Challis, 2002; McLean et al., 2003; Lloyd & Besier, 2003; Kistemaker et al., 2006; Menegaldo & Oliveira, 2009; Wang et al., 2012; Callahan et al., 2013; John et al., 2013; Dick et al., 2017; Caillet et al., 2023) |
| | Validation using a complete experimental dataset and simulating the experimental protocol available in the literature | | 30% | (Hatze, 1977; Otten, 1987; Brown et al., 1999; Thelen, 2003; Stelzer, M. & von Stryk, 2006; Mavritsaki et al., 2007; Tsianos, G. A. et al., 2012; Mörl et al., 2012; Maceri et al., 2012; Millard et al., 2013; Elias et al., 2014; Marcucci et al., 2017; Siebert et al., 2018; Rockenfeller, R. et al., 2020; Hussein et al., 2021; Millard et al., 2023) |
| | No validation / Validation against data from the literature | | 26% | All remaining studies |
| 2 | Validation against ad hoc animal individual muscle force data | | 18% | (Bahler, 1968; Durfee & Palmer, 1994; Van Zandwijk et al., 1996; Curtin et al., 1998; Shue & Crago, 1998; Ettema & Meijer, 2000; Günther et al., 2007; Blümel et al., 2012; Lee et al., 2013; Kim & Kim, 2018) |
| | Validation against ad hoc in vivo human muscle force either via (1) ultrasound-measurements of tendon length or (2) a specific experimental protocol that enables interpretation of muscle-induced joint moment | | 5% | (Hatze, 1978; Dick et al., 2017; Caillet et al., 2023) |
| | Validation against ad hoc human joint torque | | 21% | |
| | | Measured experimentally with a dynamometer / force transducer | 18% | (Hatze, 1978; Wyss & Pollak, 1984; Bobbert & van Ingen Schenau, 1990; Giat et al., 1993; Nussbaum & Chaffin, 1998; Gallucci & Challis, 2002; McLean, Scott G. et al., 2003; Lloyd & Besier, 2003; |

| | | | | |
|---|---|---|---|---|
| | | | | Menegaldo & Oliveira, 2009; Callahan et al., 2013) |
| | | Obtained from inverse dynamics analysis | 5% | (Lloyd & Besier, 2003; Wang et al., 2012; John et al., 2013) |
| | Validation against human joint angle | | 6% | (McLean et al., 2003; Kistemaker et al., 2006; Wang et al., 2012) |
| **3** | Objective criterion used for the analysis of the results | | 28% | |
| | | Root-mean square (RMS) error, or equivalent, between experimental and simulated data | 25% | (Durfee & Palmer, 1994; Nussbaum & Chaffin, 1998; McLean et al., 2003; Lloyd & Besier, 2003; Kistemaker et al., 2006; Menegaldo & Oliveira, 2009; Blümel et al., 2012; Lee et al., 2013; John et al., 2013; Kim & Kim, 2018; Rockenfeller, R. et al., 2020; Hussein et al., 2021; Millard et al., 2023; Caillet et al., 2023) |
| | | Determination / Cross correlation coefficients between experimental and simulated profiles | 9% | (Nussbaum & Chaffin, 1998; Lloyd & Besier, 2003; Menegaldo & Oliveira, 2009; Lee et al., 2013; Caillet et al., 2023) |
| | | Validation the predicted values if these fall within one or two standard deviations of the experimental data | 9% | (McLean et al., 2003; Thelen, 2003; Wang et al., 2012; Callahan et al., 2013; Siebert et al., 2018) |
| | | Mean standard deviations of the RMS and correlation coefficients for inter-trial or inter-subject procedures | 9% | (Durfee & Palmer, 1994; Nussbaum & Chaffin, 1998; Lloyd & Besier, 2003; Menegaldo & Oliveira, 2009; Blümel et al., 2012) |
| | | Analyses of Variance (ANOVA) or analyses of covariance (ANCOVA) followed by post-hoc tests for different testing sessions, gait conditions, experimental protocols or data, types of muscle models, Hill model scaling procedures, fibre type proportions and the use of mean against muscle-specific parameter values. | 12% | (Ettema & Meijer, 2000; Lloyd & Besier, 2003; Kistemaker et al., 2006; Menegaldo & Oliveira, 2009; Blümel et al., 2012; Lee et al., 2013; Dick et al., 2017) |
| | | Paired student's t- or Wilcoxon signed-rank tests for assessing the effects of model physiological correctness, parameter optimization and type of muscle model | 5% | (Lloyd & Besier, 2003; Kistemaker et al., 2006; Blümel et al., 2012) |
| | Qualitative analysis of the results | | 60% | |
| **4** | Provided enough information for the full re-implementation of the Hill-type model | | 56% | |
| | | Open-source implementation is provided | 11% | (Cheng et al., 2000; Song, D. et al., 2008; Millard et al., 2013; Kim & Kim, 2018; Millard et al., 2023; Caillet et al., 2023) |
| | | Sufficient material is provided for the full reproduction of the reported results | 5% | (Millard et al., 2013; Millard et al., 2023; Caillet et al., 2023) |
| | Missing equations, rheological arrangement of the elements, parameter values | | 44% | |
| **5** | Optimized numerical stability and computational speed, notably in avoiding singularities and numerical stiffness | | 40% | |
| | | C1 or C2 continuity for all properties (C1-C2 functions or Bezier interpolation splines) | 14% | (Van Soest & Bobbert, 1993 ; Happee, 1994; Gömmel et al., 2007; Günther et al., 2007; Rengifo et al., 2010; Millard et al., 2013; Ross et al., 2018; Rockenfeller, R. et al., 2020) |

| | | | | |
|---|---|---|---|---|
| | | Avoiding singularities and improving stability by:<br>• enforcing non-zero minimal values for the active state, the isometric muscle force and the tendon force,<br>• preventing a 90° pennation angle or non-physiological large stretching or shortening,<br>• adding a damper or a muscle mass element to the rheological structure | 32% | (Happee, 1994; Shue & Crago, 1998; Cheng et al., 2000; McLean et al., 2003; Lloyd & Besier, 2003; Günther et al., 2007; Song, D. et al., 2008; Menegaldo & Oliveira, 2009; Rengifo et al., 2010; Tsianos, G. A. et al., 2012; Blümel et al., 2012; Millard et al., 2013; John et al., 2013; Elias et al., 2014; Warner et al., 2020; Hussein et al., 2021; Millard et al., 2023; Caillet et al., 2023) |
| | | Adding a SDE to prevent unstable non-physiological mass oscillations. | 9% | (Günther et al., 2007; Mörl et al., 2012; Maceri et al., 2012; Ross et al., 2018; Rockenfeller, R. et al., 2020) |
| | Models not optimized for stability and computational speed | | 60% | |
| 6 | Explicit assumptions on muscle physiology or muscle internal/external geometry/ architecture | Massless, isotropic straight-line and fusiform model (pennation angle disregarded); fibres in parallel and not interacting with each other | 55% | |
| | Explicit comments about the simplification across physiological scales and/or explicit assumption of homogeneous and/or averaged material properties between contractile sub-scale elements | | 32% | |
| | | Building a sarcomere- or fibre-scale model | 11% | (Hatze, 1977; Cheng et al., 2000; Song, D. et al., 2008; Tsianos, G. A. et al., 2012; Marcucci et al., 2017) |
| | | Building a MU-scale model | 1% | (Hatze, 1977; Hatze, 1978; Dorgan & O'Malley, 1997; Mavritsaki et al., 2007; Callahan et al., 2013; Caillet et al., 2023) |
| | | Lumping a whole muscle as two fast and slow representative MUs | 4% | (Lee et al., 2013; Dick et al., 2017) |
| | | Assuming homogeneity in material properties, and lengths between sarcomeres, fibres or MUs further, and describing the whole muscle as a representative scaled sarcomere, fibre or MU capable of generating a whole muscle force | 23% | (Bahler, 1968; Hatze, 1977; Hatze, 1978; Dorgan & O'Malley, 1997; Cheng et al., 2000; Mavritsaki et al., 2007; Song, D. et al., 2008; Tsianos, G. A. et al., 2012; Lee et al., 2013; Millard et al., 2013; Dick et al., 2017; Ross et al., 2018; Kim & Kim, 2018) |
| | Simplifications and assumptions are not presented in the study | | 37% | |
| 7 | Modelled the force of the CE as product between active state and forces from the FL and the FV properties | | 91% | |
| | Activation and contraction dynamics are decoupled | | 51% | |
| | Using a Voigt-Kelvin-like rheological structure | | 54% | |
| | Using a Maxwell-like rheological structure | | 12% | |
| | Not using both PEE and SEE | | 40% | |
| 8 | Conclusions on the strengths and limitations of their Hill model regarding their results, the results from other studies and potential future modelling advances | | 49% | |
| 9 | At least one parameter used in the parametrization of the mathematical expressions of the normalized Hill-type model was obtained with: | | 47% | |
| | | Ad hoc muscle-specific experiments | 25% | (Bahler, 1968; Bobbert & van Ingen Schenau, 1990; Giat et al., 1993; Durfee & Palmer, 1994; Curtin et al., 1998; Brown et al., 1999; Ettema & Meijer, 2000; Günther et al., 2007; Blümel et al., 2012; Lee et al., 2013; Dick et al., 2017; Siebert et al., 2018; Kim & Kim, 2018; Caillet et al., 2023) |
| | | Parameter fitting, or calibration by minimization of a cost | 16% | (Hatze, 1978; Van Zandwijk et al., 1996; Shue & Crago, 1998; Nussbaum & Chaffin, 1998; Lloyd |

| | | | | |
|---|---|---|---|---|
| | | function between simulated and experimental data | | & Besier, 2003; Mavritsaki et al., 2007; Millard et al., 2013; Rockenfeller, R. et al., 2020; Hussein et al., 2021) |
| | | Animal-to-human parameter scaling | 2% | (Cheng et al., 2000) |
| **10** | The normalized Hill-type model is scaled with at least one architectural scaling parameter, and the parameter value is either obtained from: | | 47% | |
| | | Ad hoc experiments of individual muscles | 26% | (Bahler, 1968; Bobbert & van Ingen Schenau, 1990; Giat et al., 1993; van Soest & Bobbert, 1993; Zandwijk et al., 1996; Curtin et al., 1998; Ettema & Meijer, 2000; Günther et al., 2007; Blümel et al., 2012; Mörl et al., 2012; Lee et al., 2013; Dick et al., 2017; Siebert et al., 2018; Kim & Kim, 2018; Caillet et al., 2023) |
| | | Parameter calibration by minimization of a cost function between simulated and experimental force or torque profiles | 12% | (Durfee & Palmer, 1994; Shue & Crago, 1998; Nussbaum & Chaffin, 1998; Lloyd & Besier, 2003; Callahan et al., 2013; Millard et al., 2013; Rockenfeller, R. et al., 2020) |
| | | Scaling parameters combining subject-specific anthropometric measurements and literature data | 16% | (Hatze, 1978; Bobbert & van Ingen Schenau, 1990; van Soest & Bobbert, 1993; Van Zandwijk et al., 1996; Menegaldo & Oliveira, 2009; Wang et al., 2012; John et al., 2013; Dick et al., 2017; Ross et al., 2018; Caillet et al., 2023) |
| | Obtaining parameters from the literature for the same species | | 30% | |
