## Supplementary material for "Hill-type models of skeletal muscle and neuromuscular actuators: a systematic review": Scoring Sheet for both assessments

| Author | NE properties | | | | | | CE properties | | | | | | | | | | Passive properties | | | | | | |
|---|---|---|---|---|---|---|---|---|---|---|---|---|---|---|---|---|---|---|---|---|---|---|---|
| | ED | MURD | AD | ADC | ADL | ADV | FL | FLA | FVC | FVE | FVL | FVA | FD | RFE | OTHD | Fat | PEE | PEE2 | PDE | M | SEE | SEE2 | SDE |
| Complete | 1 | 1 | 1 | 1 | 1 | 1 | 1 | 1 | 1 | 1 | 1 | 1 | 1 | 1 | 1 | 1 | 1 | 1 | 1 | 1 | 1 | 1 | 1 |
| Bahler 1968 | 0 | 0 | 1 | 0 | 0 | 1 | 1 | 0 | 1 | 0 | 0 | 1 | 0 | 0 | 0 | 0 | 0 | 0 | 0 | 0 | 1 | 0 | 0 |
| Hatze 1977 | 1 | 0 | 1 | 1 | 1 | 0 | 1 | 0 | 1 | 1 | 0 | 0 | 0 | 0 | 0 | 0 | 1 | 1 | 1 | 0 | 0 | 1 | 0 |
| Hatze 1978 | 1 | 1 | 1 | 1 | 1 | 0 | 1 | 0 | 1 | 1 | 0 | 0 | 0 | 0 | 0 | 0 | 0 | 1 | 0 | 0 | 0 | 1 | 0 |
| Wyss 1984 | 0 | 0 | 1 | 0 | 0 | 0 | 1 | 0 | 1 | 0 | 1 | 0 | 0 | 0 | 0 | 0 | 0 | 0 | 0 | 0 | 0 | 1 | 0 |
| Pierrynowski 1985 | 0 | 1 | 1 | 0 | 0 | 0 | 1 | 0 | 1 | 1 | 0 | 0 | 0 | 0 | 0 | 0 | 0 | 0 | 0 | 0 | 1 | 1 | 0 |
| Otten 1987 | 0 | 0 | 1 | 1 | 0 | 0 | 1 | 0 | 1 | 1 | 0 | 0 | 0 | 0 | 0 | 0 | 1 | 0 | 0 | 0 | 0 | 0 | 0 |
| Bobbert 1990 | 0 | 0 | 1 | 0 | 0 | 0 | 1 | 0 | 1 | 0 | 1 | 0 | 0 | 0 | 0 | 0 | 1 | 0 | 0 | 0 | 1 | 0 | 0 |
| Giat 1993 | 0 | 0 | 1 | 0 | 0 | 0 | 1 | 0 | 1 | 1 | 0 | 0 | 0 | 0 | 0 | 1 | 1 | 0 | 1 | 1 | 1 | 0 | 0 |
| Van Soest 1993 | 0 | 0 | 1 | 1 | 0 | 0 | 1 | 0 | 1 | 1 | 1 | 1 | 0 | 0 | 0 | 0 | 0 | 0 | 0 | 0 | 1 | 0 | 0 |
| Happee 1994 | 1 | 0 | 1 | 0 | 0 | 0 | 0 | 0 | 1 | 1 | 0 | 0 | 0 | 0 | 0 | 0 | 1 | 0 | 0 | 0 | 1 | 0 | 0 |
| Durfee 1994 | 0 | 0 | 1 | 0 | 0 | 0 | 1 | 0 | 1 | 1 | 0 | 0 | 0 | 0 | 0 | 0 | 1 | 0 | 1 | 0 | 1 | 0 | 0 |
| Winters 1995 | 1 | 0 | 1 | 0 | 0 | 0 | 1 | 1 | 1 | 1 | 1 | 1 | 0 | 0 | 0 | 0 | 1 | 1 | 0 | 0 | 1 | 1 | 0 |
| Caldwell 1995 | 0 | 0 | 1 | 0 | 1 | 0 | 1 | 0 | 1 | 1 | 0 | 0 | 0 | 0 | 0 | 0 | 0 | 0 | 0 | 0 | 1 | 0 | 0 |
| Van Zandwijk 1996 | 1 | 0 | 1 | 1 | 1 | 0 | 1 | 0 | 1 | 1 | 0 | 0 | 0 | 0 | 0 | 0 | 1 | 0 | 0 | 0 | 1 | 0 | 0 |
| Dorgan 1997 | 1 | 1 | 1 | 1 | 1 | 0 | 1 | 0 | 1 | 1 | 0 | 0 | 0 | 0 | 0 | 0 | 1 | 0 | 1 | 0 | 1 | 0 | 0 |
| Shue 1998 | 0 | 0 | 1 | 0 | 1 | 1 | 1 | 0 | 1 | 1 | 0 | 0 | 0 | 0 | 1 | 0 | 0 | 0 | 1 | 1 | 1 | 0 | 0 |
| Nussbaum 1998 | 0 | 0 | 1 | 0 | 0 | 0 | 1 | 0 | 0 | 0 | 0 | 0 | 0 | 0 | 0 | 0 | 1 | 0 | 0 | 0 | 0 | 0 | 0 |
| Curtin 1998 | 0 | 0 | 1 | 1 | 0 | 0 | 0 | 0 | 1 | 1 | 0 | 0 | 0 | 0 | 0 | 0 | 0 | 0 | 0 | 0 | 1 | 0 | 0 |
| Brown 1999 | 0 | 0 | 1 | 0 | 1 | 1 | 1 | 0 | 1 | 1 | 1 | 0 | 0 | 0 | 1 | 0 | 1 | 1 | 0 | 0 | 1 | 0 | 0 |
| Cheng 2000 | 0 | 1 | 1 | 0 | 1 | 1 | 1 | 0 | 1 | 1 | 1 | 0 | 0 | 0 | 1 | 0 | 1 | 1 | 1 | 1 | 1 | 0 | 0 |
| Ettema 2000 | 0 | 0 | 1 | 1 | 0 | 0 | 1 | 0 | 1 | 1 | 0 | 0 | 1 | 1 | 0 | 0 | 1 | 0 | 0 | 0 | 1 | 0 | 0 |
| De Woody 2001 | 0 | 0 | 1 | 0 | 0 | 0 | 1 | 0 | 1 | 1 | 0 | 0 | 0 | 0 | 0 | 0 | 1 | 0 | 1 | 1 | 1 | 0 | 0 |
| Gallucci 2002 | 0 | 0 | 1 | 0 | 0 | 0 | 1 | 0 | 1 | 1 | 0 | 0 | 0 | 0 | 0 | 0 | 0 | 0 | 0 | 0 | 1 | 0 | 0 |
| Lloyd 2003 | 0 | 0 | 1 | 0 | 0 | 0 | 1 | 1 | 1 | 1 | 0 | 0 | 0 | 0 | 0 | 0 | 1 | 0 | 1 | 0 | 1 | 0 | 0 |
| Thelen 2003 | 0 | 0 | 1 | 0 | 0 | 0 | 1 | 0 | 1 | 1 | 0 | 1 | 0 | 0 | 0 | 0 | 1 | 0 | 0 | 0 | 1 | 0 | 0 |
| McLean 2003 | 0 | 0 | 1 | 0 | 0 | 0 | 1 | 0 | 1 | 1 | 0 | 1 | 0 | 0 | 0 | 0 | 1 | 0 | 0 | 0 | 1 | 0 | 0 |
| Stelzer 2006 | 0 | 0 | 1 | 1 | 0 | 0 | 1 | 0 | 1 | 1 | 0 | 0 | 0 | 0 | 0 | 0 | 1 | 0 | 1 | 0 | 0 | 0 | 0 |
| Kistemaker 2006 | 0 | 0 | 1 | 1 | 1 | 0 | 1 | 0 | 1 | 1 | 1 | 1 | 0 | 0 | 0 | 0 | 1 | 0 | 0 | 0 | 1 | 0 | 0 |
| Günther 2007 | 0 | 0 | 1 | 0 | 0 | 0 | 1 | 0 | 1 | 1 | 1 | 1 | 0 | 0 | 0 | 0 | 1 | 0 | 1 | 0 | 1 | 0 | 1 |
| Mavritsaki 2007 | 1 | 1 | 1 | 1 | 0 | 0 | 1 | 0 | 1 | 0 | 0 | 0 | 0 | 0 | 0 | 0 | 1 | 0 | 0 | 0 | 0 | 0 | 0 |
| Gömmel 2008 | 0 | 0 | 1 | 0 | 0 | 0 | 1 | 0 | 1 | 1 | 1 | 1 | 0 | 0 | 0 | 0 | 1 | 0 | 0 | 0 | 0 | 0 | 0 |
| Song 2008 | 1 | 1 | 1 | 1 | 1 | 1 | 1 | 0 | 1 | 1 | 1 | 0 | 0 | 0 | 1 | 0 | 1 | 1 | 1 | 1 | 1 | 0 | 0 |
| Menegaldo 2009 | 0 | 0 | 1 | 0 | 0 | 0 | 1 | 0 | 1 | 1 | 0 | 1 | 0 | 0 | 0 | 0 | 1 | 0 | 1 | 0 | 1 | 0 | 0 |
| Moody 2009 | 0 | 0 | 1 | 0 | 0 | 0 | 1 | 0 | 1 | 1 | 0 | 0 | 0 | 0 | 0 | 0 | 1 | 0 | 1 | 1 | 1 | 0 | 0 |
| Rengifo 2010 | 0 | 0 | 1 | 0 | 0 | 0 | 1 | 0 | 1 | 1 | 0 | 0 | 0 | 0 | 0 | 0 | 1 | 0 | 0 | 0 | 1 | 0 | 0 |
| Proctor 2010 | 1 | 0 | 1 | 1 | 0 | 0 | 1 | 0 | 1 | 1 | 0 | 0 | 0 | 0 | 0 | 0 | 1 | 0 | 1 | 0 | 0 | 0 | 0 |
| Tsianos 2011 | 0 | 1 | 1 | 1 | 1 | 1 | 1 | 0 | 1 | 1 | 1 | 0 | 0 | 0 | 1 | 0 | 1 | 1 | 1 | 1 | 1 | 0 | 0 |
| Mörl 2012 | 0 | 0 | 1 | 0 | 0 | 0 | 1 | 0 | 1 | 0 | 1 | 1 | 0 | 0 | 0 | 0 | 1 | 0 | 0 | 0 | 1 | 0 | 1 |
| Wang 2012 | 0 | 0 | 1 | 0 | 0 | 0 | 1 | 0 | 1 | 1 | 0 | 0 | 0 | 0 | 0 | 0 | 1 | 0 | 0 | 0 | 1 | 0 | 0 |
| Blümel 2012 | 1 | 0 | 1 | 0 | 0 | 0 | 1 | 1 | 1 | 1 | 0 | 1 | 0 | 0 | 0 | 0 | 1 | 0 | 1 | 1 | 1 | 0 | 0 |
| Maceri 2012 | 1 | 0 | 1 | 0 | 0 | 0 | 1 | 0 | 1 | 1 | 0 | 0 | 0 | 0 | 0 | 0 | 1 | 0 | 1 | 0 | 1 | 0 | 1 |
| Callahan 2013 | 1 | 1 | 1 | 0 | 0 | 0 | 1 | 0 | 1 | 0 | 1 | 1 | 0 | 0 | 0 | 0 | 0 | 0 | 0 | 0 | 1 | 0 | 0 |
| Millard 2013 | 0 | 0 | 1 | 0 | 0 | 0 | 1 | 0 | 1 | 1 | 0 | 0 | 0 | 0 | 0 | 0 | 1 | 0 | 1 | 1 | 1 | 0 | 0 |
| Chand 2013 | 0 | 0 | 1 | 0 | 0 | 0 | 1 | 0 | 1 | 1 | 0 | 1 | 0 | 0 | 0 | 0 | 1 | 0 | 1 | 0 | 1 | 0 | 0 |
| Lee 2013 | 0 | 1 | 1 | 0 | 0 | 0 | 1 | 0 | 1 | 1 | 0 | 0 | 0 | 0 | 0 | 0 | 1 | 0 | 0 | 0 | 0 | 0 | 0 |
| Elias 2014 | 1 | 1 | 1 | | 0 | 0 | 1 | 0 | 1 | 1 | 1 | 0 | 0 | 0 | 0 | 0 | 1 | 0 | 1 | 1 | 1 | 0 | 0 |
| Hamouda 2016 | 0 | 1 | 0 | 0 | 0 | 0 | 1 | 0 | 1 | 1 | 0 | 0 | 0 | 0 | 0 | 0 | 1 | 0 | 0 | 0 | 1 | 0 | 0 |
| Dick 2017 | 0 | 1 | 1 | 0 | 0 | 0 | 1 | 0 | 1 | 1 | 0 | 0 | 0 | 0 | 0 | 0 | 1 | 0 | 0 | 0 | 0 | 0 | 0 |
| Marcucci 2017 | 0 | 0 | 1 | 0 | 0 | 0 | 1 | 0 | 1 | 0 | 0 | 0 | 0 | 0 | 0 | 0 | 1 | 0 | 0 | 0 | 0 | 1 | 0 |
| Ross 2018 | 0 | 0 | 1 | 0 | 0 | 0 | 1 | 0 | 1 | 1 | 0 | 0 | 0 | 0 | 0 | 0 | 1 | 0 | 0 | 1 | 0 | 0 | 0 |
| Kim 2015, 2018 | 1 | 0 | 1 | 1 | 1 | 0 | 1 | 0 | 1 | 1 | 0 | 0 | 0 | 0 | 0 | 0 | 0 | 0 | 0 | 0 | 1 | 0 | 0 |
| Siebert 2018 | 0 | 0 | 1 | 0 | 0 | 0 | 1 | 0 | 1 | 0 | 0 | 0 | 0 | 0 | 1 | 0 | 0 | 0 | 0 | 0 | 1 | 0 | 0 |
| Rockenfeller 2020 | 0 | 0 | 1 | 1 | 1 | 0 | 1 | 0 | 1 | 1 | 1 | 1 | 0 | 0 | 0 | 0 | 1 | 0 | 0 | 0 | 1 | 0 | 1 |
| Warner 2020 | 0 | 0 | 1 | 0 | 0 | 0 | 1 | 0 | 1 | 1 | 0 | 0 | 0 | 0 | 0 | 0 | 1 | 0 | 0 | 0 | 1 | 0 | 0 |
| Hussein 2022 | 0 | 0 | 1 | 1 | 0 | 0 | 1 | 0 | 1 | 0 | 0 | 1 | 0 | 0 | 0 | 0 | 1 | 0 | 1 | 0 | 1 | 0 | 0 |
| Millard 2023 | 0 | 0 | 1 | 0 | 0 | 0 | 1 | 0 | 1 | 1 | 0 | 0 | 0 | 0 | 1 | 0 | 1 | 1 | 1 | 0 | 1 | 1 | 1 |
| Caillet 2023 | 1 | 1 | 1 | 1 | 1 | 0 | 1 | 1 | 0 | 0 | 0 | 0 | 0 | 0 | 0 | 0 | 1 | 0 | 0 | 0 | 1 | 0 | 0 |

| | VALIDATION | | | REUSABILITY | | MODEL. CHOICES & STRATEGY | | | CALIBRATION | |
|---|---|---|---|---|---|---|---|---|---|---|
| Author | Q1 | Q2 | Q3 | Q4 | Q5 | Q6 | Q7 | Q8 | Q9 | Q10 |
| Complete | 2 | 2 | 2 | 2 | 2 | 2 | 2 | 2 | 2 | 2 |
| Bahler 1968 | 2 | 2 | 0 | 1 | 0 | 2 | 2 | 1 | 2 | 2 |
| Hatze 1977 | 1 | 2 | 0 | 2 | 2 | 2 | 2 | 2 | 1 | 1 |
| Hatze 1978 | 2 | 2 | 0 | 2 | 2 | 2 | 2 | 2 | 1 | 2 |
| Wyss 1984 | 2 | 1 | 0 | 0 | 0 | 0 | 2 | 1 | 0 | 0 |
| Pierrynowski 1985 | 0 | 0 | 0 | 2 | 0 | 1 | 2 | 1 | 1 | 1 |
| Otten 1987 | 1 | 1 | 0 | 1 | 0 | 1 | 1 | 2 | 1 | 1 |
| Bobbert 1990 | 2 | 1 | 0 | 1 | 0 | 0 | 2 | 2 | 2 | 2 |
| Giat 1993 | 2 | 1 | 0 | 2 | 0 | 1 | 2 | 1 | 2 | 2 |
| Van Soest 1993 | 0 | 0 | 0 | 1 | 0 | 0 | 1 | 0 | 0 | 2 |
| Happee 1994 | 0 | 0 | 0 | 1 | 2 | 0 | 1 | 2 | 0 | 0 |
| Durfee 1994 | 2 | 2 | 2 | 0 | 1 | 0 | 2 | 2 | 2 | 2 |
| Winters 1995 | 0 | 0 | 0 | 2 | 0 | 1 | 2 | 1 | 1 | 0 |
| Caldwell 1995 | 0 | 0 | 0 | 1 | 0 | 0 | 2 | 2 | 0 | 0 |
| Van Zandwijk 1996 | 2 | 2 | 0 | 1 | 0 | 1 | 2 | 1 | 2 | 2 |
| Dorgan 1997 | 1 | 2 | 0 | 2 | 2 | 2 | 2 | 1 | 0 | 0 |
| Shue 1998 | 2 | 2 | 0 | 1 | 2 | 0 | 2 | 2 | 2 | 2 |
| Nussbaum 1998 | 2 | 1 | 2 | 1 | 0 | 0 | 1 | 1 | 2 | 2 |
| Curtin 1998 | 2 | 2 | 0 | 1 | 0 | 0 | 2 | 2 | 2 | 2 |
| Brown 1999 | 1 | 1 | 0 | 2 | 0 | 0 | 2 | 2 | 2 | 1 |
| Cheng 2000 | 0 | 0 | 0 | 2 | 2 | 2 | 2 | 1 | 2 | 1 |
| Ettema 2000 | 2 | 2 | 2 | 1 | 0 | 0 | 1 | 1 | 2 | 2 |
| De Woody 2001 | 0 | 0 | 0 | 1 | 0 | 1 | 1 | 0 | 0 | 0 |
| Gallucci 2002 | 2 | 1 | 0 | 2 | 0 | 1 | 1 | 2 | 0 | 1 |
| Lloyd 2003 | 2 | 1 | 2 | 0 | 2 | 1 | 1 | 2 | 2 | 2 |
| Thelen 2003 | 1 | 1 | 1 | 2 | 0 | 0 | 2 | 2 | 1 | 1 |
| McLean 2003 | 2 | 1 | 2 | 2 | 2 | 0 | 1 | 0 | 1 | 1 |
| Stelzer 2006 | 1 | 1 | 0 | 2 | 0 | 0 | 1 | 0 | 0 | 0 |
| Kistemaker 2006 | 2 | 1 | 2 | 2 | 2 | 1 | 1 | 0 | 0 | 1 |
| Günther 2007 | 2 | 2 | 0 | 2 | 2 | 1 | 2 | 2 | 2 | 2 |
| Mavritsaki 2007 | 1 | 2 | 0 | 1 | 0 | 2 | 1 | 1 | 2 | 1 |
| Gömmel 2008 | 0 | 0 | 0 | 1 | 0 | 1 | 1 | 1 | 0 | 0 |
| Song 2008 | 0 | 0 | 1 | 2 | 2 | 2 | 2 | 1 | 2 | 0 |
| Menegaldo 2009 | 2 | 1 | 2 | 1 | 0 | 0 | 1 | 0 | 0 | 2 |
| Moody 2009 | 0 | 0 | 0 | 1 | 0 | 1 | 1 | 0 | 0 | 0 |
| Rengifo 2010 | 0 | 0 | 0 | 1 | 2 | 1 | 1 | 0 | 0 | 0 |
| Proctor 2010 | 0 | 0 | 0 | 2 | 0 | 0 | 1 | 1 | 1 | 1 |
| Tsianos 2011 | 1 | 1 | 1 | 2 | 1 | 2 | 2 | 1 | 2 | 0 |
| Mörl 2012 | 1 | 1 | 1 | 2 | 0 | 1 | 2 | 2 | 1 | 2 |
| Wang 2012 | 2 | 1 | 0 | 2 | 0 | 0 | 1 | 0 | 0 | 2 |
| Blümel 2012 | 2 | 2 | 2 | 1 | 2 | 0 | 2 | 2 | 2 | 2 |
| Maceri 2012 | 1 | 1 | 0 | 1 | 2 | 2 | 2 | 1 | 1 | 1 |
| Callahan 2013 | 2 | 1 | 1 | 2 | 0 | 2 | 1 | 1 | 0 | 2 |
| Millard 2013 | 1 | 2 | 1 | 2 | 2 | 2 | 2 | 2 | 2 | 2 |
| Chand 2013 | 2 | 1 | 2 | 2 | 2 | 0 | 2 | 2 | 0 | 2 |
| Lee 2013 | 2 | 2 | 2 | 2 | 0 | 2 | 1 | 2 | 2 | 2 |
| Elias 2014 | 1 | 1 | 0 | 1 | 2 | 1 | 2 | 0 | 0 | 1 |
| Hamouda 2016 | 0 | 0 | 0 | 2 | 0 | 1 | 2 | 1 | 0 | 0 |
| Dick 2017 | 2 | 2 | 2 | 2 | 1 | 2 | 2 | 2 | 2 | 2 |
| Marcucci 2017 | 1 | 1 | 0 | 1 | 0 | 2 | 2 | 2 | 2 | 1 |
| Ross 2018 | 0 | 0 | 0 | 2 | 2 | 2 | 1 | 2 | 0 | 2 |
| Kim 2015, 2018 | 2 | 2 | 2 | 2 | 2 | 2 | 1 | 2 | 2 | 2 |
| Siebert 2018 | 1 | 2 | 1 | 2 | 0 | 1 | 2 | 2 | 2 | 2 |
| Rockenfeller 2020 | 1 | 2 | 2 | 2 | 1 | 0 | 2 | 2 | 2 | 2 |
| Warner 2020 | 0 | 0 | 0 | 2 | 2 | 0 | 1 | 0 | 0 | 1 |
| Hussein 2022 | 1 | 1 | 2 | 1 | 2 | 1 | 2 | 2 | 2 | 1 |
| Millard 2023 | 1 | 2 | 2 | 2 | 2 | 2 | 2 | 2 | 2 | 1 |
| Caillet 2023 | 2 | 1 | 2 | 2 | 2 | 2 | 2 | 2 | 2 | 2 |
